# Directed biosynthesis of fluorinated polyketides

**DOI:** 10.1101/2021.07.30.454469

**Authors:** Alexander Rittner, Mirko Joppe, Jennifer J. Schmidt, Lara Maria Mayer, Elia Heid, David H. Sherman, Martin Grininger

## Abstract

Modification of polyketides with fluorine offers a promising approach to develop new pharmaceuticals. While synthetic chemical methods for site-specific incorporation of fluorine in complex molecules have improved in recent years, approaches for the direct biosynthetic fluorination of natural compounds are still rare. Herein, we present a broadly applicable approach for site-specific, biocatalytic derivatization of polyketides with fluorine. Specifically, we exchanged the native acyltransferase domain (AT) of a polyketide synthase (PKS), which acts as the gatekeeper for selection of extender units, with an evolutionarily related but substrate tolerant domain from metazoan type I fatty acid synthase (FAS). The resulting PKS/FAS hybrid can utilize fluoromalonyl coenzyme A and fluoromethylmalonyl coenzyme A for polyketide chain extension, introducing fluorine or fluoro-methyl disubstitutions in polyketide scaffolds. Addition of a fluorine atom is often a decisive factor toward developing superior properties in next-generation antibiotics, including the macrolide solithromycin. We demonstrate the feasibility of our approach in the semisynthesis of a fluorinated derivative of the macrolide antibiotic YC-17.

## Main Text

The majority of new FDA-approved drugs are small organic molecules, making up over 90 percent of pharmaceuticals on the market today ^1^. Among those, natural products are highly represented ^2^, as their structures are presumed to undergo preselection during evolution to interact with cellular biomacromolecules ^3^. Fluorination has been widely used in medicinal chemistry for lead structure optimization, as its electronegativity and its small size can strongly impact molecular properties, thereby modulating protein–ligand interactions, bioavailability and metabolic stability ^4^. Reflecting the importance of fluorination, about a quarter of all small molecule drugs contain at least one fluorine atom, including the antidepressant Prozac, the cholesterol-lowering drug Lipitor and quinolone antibiotic Ciprobay ^5^. Applications of organofluorine chemistry toward natural product biosynthesis are rare due to the paucity of enzymes that catalyze addition of F atoms in secondary metabolism ^6^. Thus, new methods that enable fluorine derivatization are urgently needed to bridge the gap between the inherent bioactivity of a natural compound and its development as a human therapeutic agent.

Polyketide natural products comprise over 10,000 molecules with a wide range of bioactivities and are among the most prominent classes of approved clinical agents ^7,8^. In nature, polyketides are assembled mainly from simple monomeric acetate and propionate units by polyketide synthases (PKSs). In the type I cis-AT subclass, PKSs occur as multi-functional protein mega-complexes comprising a series of catalytic domains organized in modules on one or few polypeptide chains. Typically, one PKS module requires a minimum of three domains for a two-carbon extension of a growing polyketide intermediate: an acyltransferase (AT) domain that selects an acyl-coenzyme A (CoA) extender unit and transfers the acyl moiety to the acyl carrier protein (ACP) domain, and a ketosynthase (KS) domain that accepts a growing chain from the ACP of the previous module and catalyzes a decarboxylative Claisen condensation to extend the polyketide chain. A canonical PKS module may further contain up to three additional domains, a ketoreductase (KR), a dehydratase (DH) and an enoylreductase (ER) that tailor the β-keto functionality prior to the next round of chain extension. The final module in the biosynthetic assembly line typically contains a thioesterase (TE) domain located at the C-terminus and is responsible for polyketide chain release as a linear chain or a macrocyclic product. Engineering of modular polyketide biosynthesis for the directed assembly of new-to-nature polyketides is a highly aspired aim ^9^, and offers an alternative or complementary approach to organic synthesis. Changing substrate specificity of a single enzymatic domain of a specific PKS module by protein engineering enables, for example, the regioselective modification of the product during biosynthesis.

Enzymes that catalyze direct fluorination of polyketides remain unknown. Previous efforts have demonstrated that engineered PKS assembly lines at even-numbered carbon centers enable loading of non-canonical extender substrates during the chain elongation process (Fig. 1a)^10^. Indeed, also fluoromalonyl-CoA was incorporated into a growing triketide chain followed by spontaneous off-loading lactonization ^11,12^. However, the application of this concept to the formation of a complete macrolide structure had not been demonstrated. In canonical modular PKSs, extender subunits are selected by the AT domains, which act as the “gatekeepers” of polyketide biosynthesis and typically ensure the introduction of a defined acyl-CoA with high substrate specificity (Supplementary Fig. S1). We have recently demonstrated that the promiscuous AT domain from metazoan fatty acid synthase (FAS), termed malonyl-/acetyl transferase (MAT), is able to transfer various acyl-CoA moieties with high efficiencies ^13^, different to AT domains from PKSs ^14^. We hypothesized that this domain may also transfer fluorinated extender substrates into a PKS module, since FASs and PKSs are structurally and biochemically related (Fig 1b) ^15-17^. To test the feasibility of this approach, we chose to work with module 6 of DEBS including its C-terminal TE domain (DEBS M6+TE) (Fig. 1c) ^18^.

**Fig. 1:**
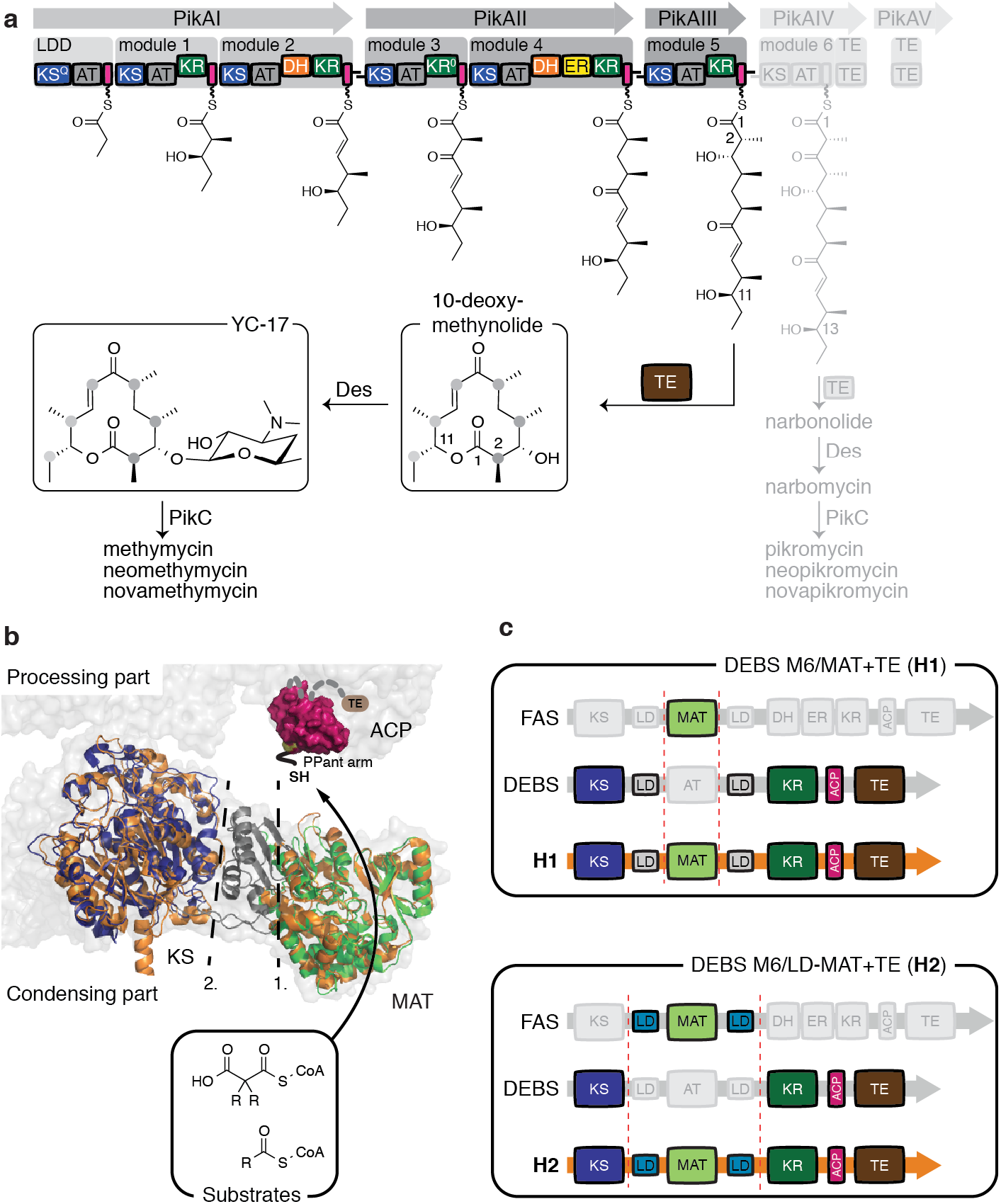
Modular PKSs and hybrid design. **a**, Assembly line like biosynthesis in the methymycin/pikromycin pathway. The modular pikromycin synthase can either produce a 12- or a 14-membered macrolactone. The polyketide products are subsequently glycosylated and oxidized by post-PKS enzymes. **b**, Function of the murine MAT domain and its insertion into the KS-MAT didomain (KS: blue; LD: grey; MAT: green; PDB code: 5my0). Atomic coordinates: porcine FAS (grey; PDB code: 2vz9), the ACP domain (purple; PDB code: 2png) and DEBS module 5 KS-AT didomain (orange; PDB code: 2hg4) ^13,15,29,30^. R – corresponds to various chemical groups ^13^. **c**, Design of DEBS/FAS hybrids **H1** and **H2**.

## Results and discussion

Initially, we tested whether the polyspecific MAT domain of murine FAS is able to select fluoromalonyl-coenzyme A (F-Mal-CoA) as a substrate, and whether it accepts the ACP domain of the DEBS M6 for substrate processing. F-Mal-CoA (**1**) was chemically synthesized following the four-step route to the thiophenyl-ester by Saadi and Wennemers ^19^ with the additional subsequent transacylation of the F-Mal moiety to free CoA (Fig. 2a, Supplementary Fig. S2-S3). In an enzyme-coupled fluorometric assay with the domains as individual proteins ^13^, we observed excellent transfer kinetics of MAT for F-Mal-CoA (*K*_m_/*k*_cat_ = 6.9 10^6^ M^−1^ s^−1^) as well as its ability to catalyze transacylation with DEBS ACP6 (Supplementary Fig. S4-S7). The specificity constant of MAT for loading ACP6 with methylmalonyl moieties (*K*_m_/*k*_cat_ = 6.9 10^6^ M^−1^ s^−1^) was 2 -3 orders of magnitude higher than DEBS AT6 (Supplementary Table S1), which can be explained by the inherently high transacylation rates of the MAT domain.

**Fig. 2:**
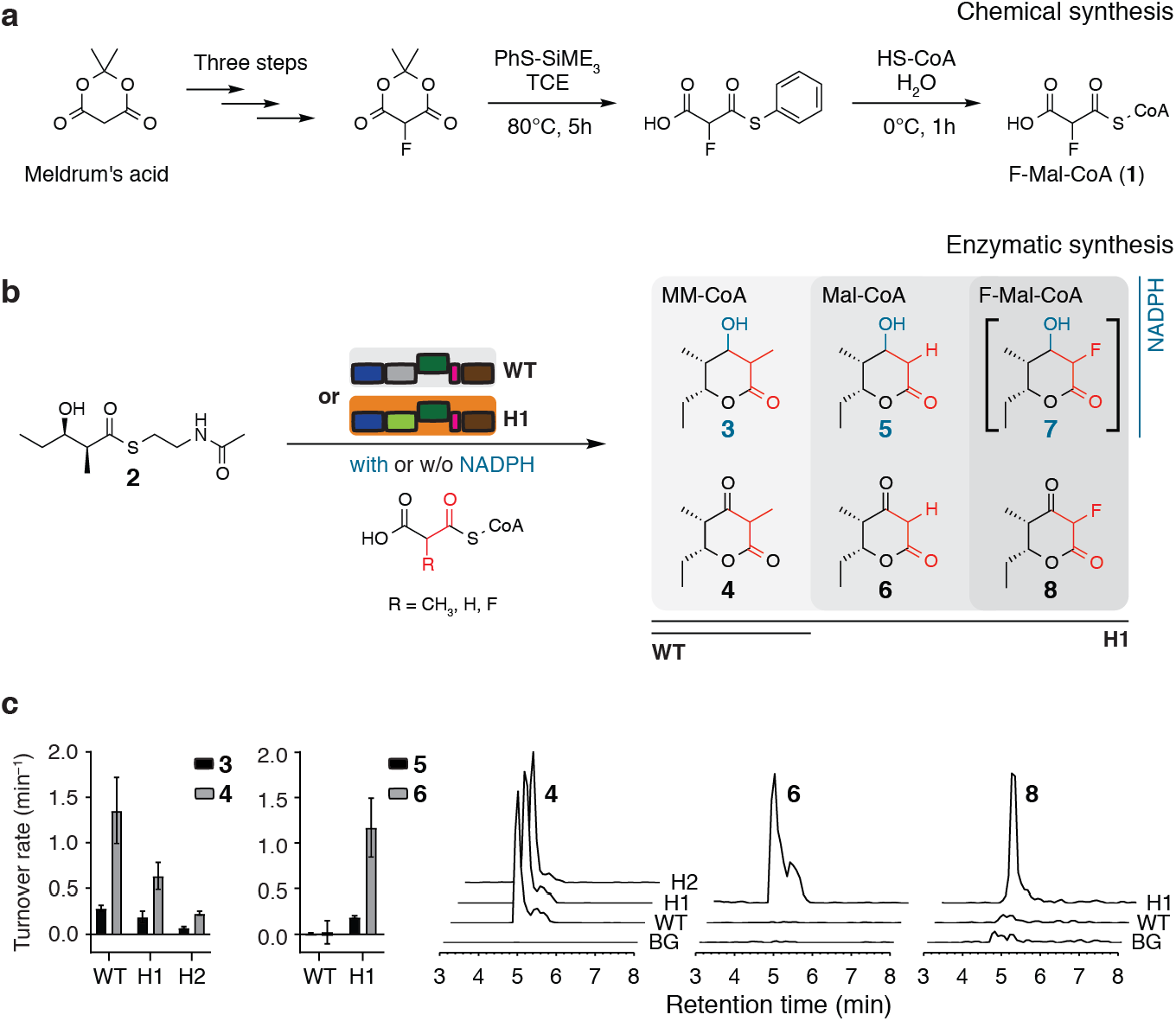
Function of the hybrid DEBS/FAS modules. **a**, Synthetic route to F-Mal-CoA. **b**, Hybrid PKS-mediated synthesis of triketide lactones (TKLs) from **2** and MM-CoA, Mal-CoA or F-Mal-CoA. Compound **7** was only produced in traces (not shown), presumably due to a substrate selective KR domain. **c**, Turnover rates for the **WT**-, **H1**- and **H2**-mediated formation of TKLs and detection by HPLC-MS (EIC: 4 [M-H]^-^ *m*/*z* = 169.12; 6 [M-H]^-^ *m*/*z* = 155.16; 8 [M-H]^-^ *m*/*z* = 173.11).

We constructed two hybrids of *S. erythraea* DEBS and murine FAS, which differed in the DEBS/FAS interface by substituting DEBS AT6 with or without its adjacent linker domain, giving construct **H1** (MAT hybrid) or **H2** (LD-MAT hybrid), respectively (Fig. 1c, Supplementary Fig. S8). The hybrids **H1** and **H2** were produced in *Escherichia coli* in yields similar to the wildtype protein (**WT**), but with different oligomeric stability (Supplementary Fig. S9). This provided overall yields of purified dimeric species of about 4 and 2 mg per liter of cell culture for **H1** and **H2**, respectively (compared to 8 mg of DEBS M6+TE **WT**). Native PAGE further indicated contamination of **H2** preparations with degraded or disassembled PKS proteins. On the basis of **H1** showing significantly higher turnover rates in synthesizing triketide lactones (TKLs) from diketide N-acetylcysteamine compound **2** and MM-CoA (67% of **WT** rate compared to **H2** with 25% of **WT** rate), we decided to pursue further efforts with **H1** only (Fig. 2b-2c, Supplementary Fig. S10 and Supplementary Table S2). Indeed, **H1** was able to produce C-2 derivatives of TKLs with the extender substrates malonyl-CoA (Mal-CoA; the native substrate of MAT) and F-Mal-CoA, demonstrating that the substrate promiscuity of MAT enables substrate elongation with a fluorinated extender unit.

With the catalytically competent hybrid **H1** in hand, we aimed to produce 12-membered macrolactones from the pikromycin pathway that are diversified at the C-2 position (Fig. 1a, Supplementary Fig. S11). These are particularly interesting as exhibiting microbial activity against erythromycin-resistant *Staphylococcus aureus* strains and bind in a unique way to the 50S ribosomal subunit ^20-22^. Elongation of the pentaketide (**9**) with MM-CoA, Mal-CoA and F-CoA, respectively, produced 10-deoxymethynolide (**10**) as well as the respective desmethylated macrocycle (**12**) and the fluorinated analog (**14**) ^23^. The absence of NADPH led to the C-3 oxidized species (**11, 13** and **15**) (compounds **10**–**15** were confirmed by HRMS; Fig. 3a and b, Supplementary Fig. S12 and Supplementary Table S2). The **H1**-mediated conversion rates were faster for Mal-CoA and slightly slower for F-Mal-CoA compared to the native substrate MM-CoA yielding compounds **12, 14** and **10**, respectively (Fig. 3c, Supplementary Table S2). Intriguingly, when seeking to conduct scale-up to isolate milligram quantities, we faced challenges for the reactions to the fluoro-compounds **14** and **15**. Here, very low amounts of products were obtained, and reactions were contaminated with significant levels of side products. When working-up the reaction mixture to target compound **15**, we identified compound **16** as the main product, presumably generated from the hexaketide intermediate via hydrolysis, decarboxylation and cyclohexanone formation (Fig. 3d). This side reaction has been reported previously as originating from the narrow substrate specificity of the DEBS TE domain, preventing macrolactonization to the 12-membered ring when a keto-group is present at C-3 ^24^. We elected to pause further analysis of the origin for the low yields of compound **14**, and reasoned that chemical instability relates to the C-2 HFC group.

**Fig. 3:**
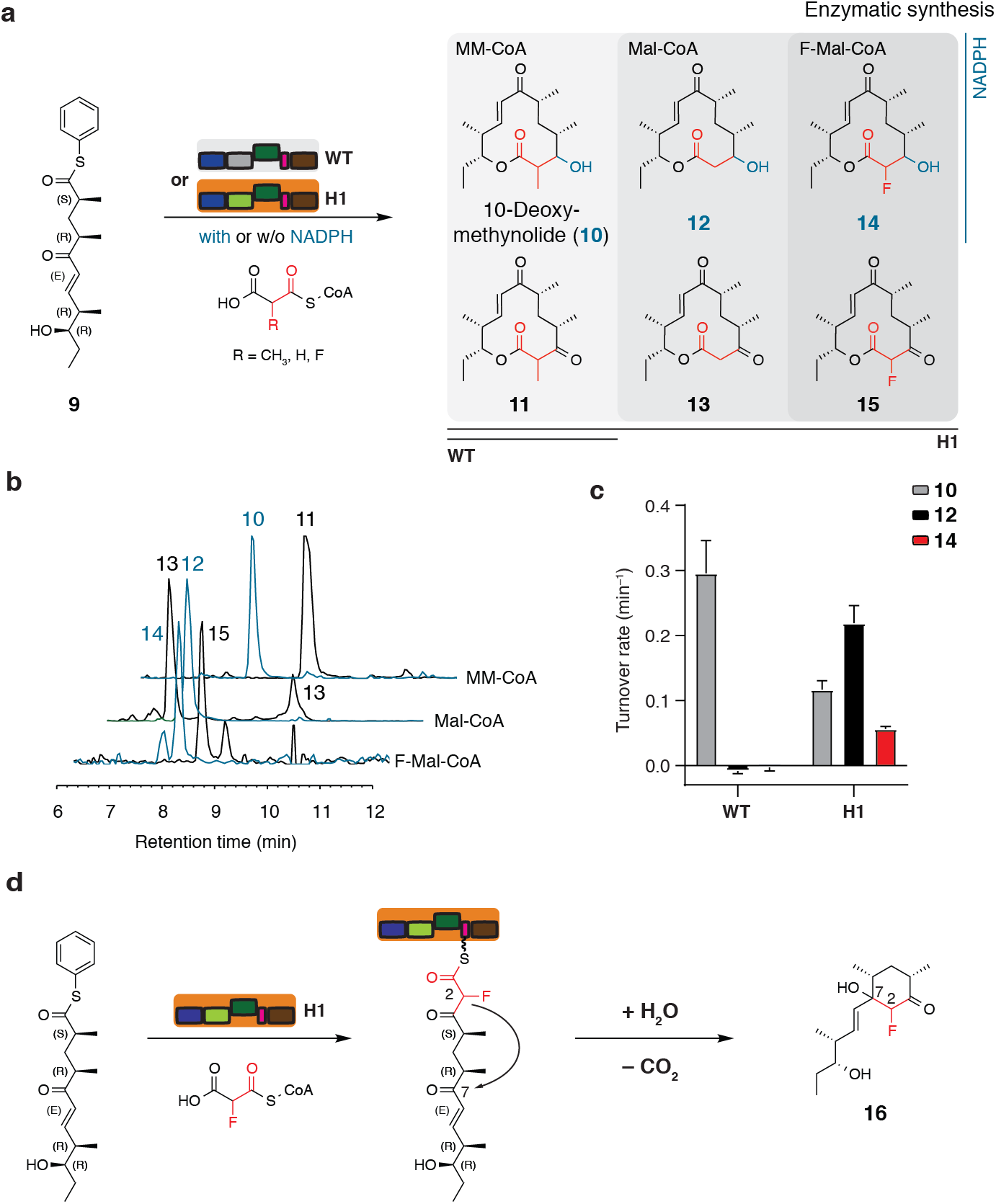
Enzymatic synthesis of 10-deoxymethynolide derivatives. **a**, Reaction scheme for the **H1**-mediated conversion of pentaketide **9** to new derivatized keto- and macrolactones **10**-**15** (see also Table S2). **b**, Detection of macrolactones by HPLC-MS (EIC: **10** [M+Na]^+^ *m*/*z* = 319.11; **11** [M+Na]^+^ *m*/*z* = 317.09; **12** [M+H]^+^ *m*/*z* = 305.09; **13** [M+Na]^+^ *m*/*z* = 303.08; **14** [M+Na]^+^ *m*/*z* = 323.08 and **15** [M+Na]^+^ *m*/*z* = 321.07). **c**, Turnover rates for **H1**-mediated macrolactone formation in comparison with the **WT** turnover rate. **d**, Formation of side product **16** during the synthesis of fluoro-compound **15**.

As a next step, we postulated that direct installation of a C-2 fluoro-methyl disubstitution (MeFC group) might increase stability by abstracting the acidic proton while maintaining a similar size compared to the natural compound. Notably, this would give direct access to fluorination patterns of the erythromycin derivatives flurithromycin and solithromycin (Supplementary Fig. S13), two examples of semisynthetic next-generation macrolides, in which acidic protons have been replaced by fluorine ^25^. For solithromycin, the fluorine induces improved binding to the ribosome and modulates pharmacokinetics leading to superior antibiotic properties (23S rRNA binding) ^26,27^.

Natural polyketides with MeFC moieties are not known and just a handful of polyketides with a *gem*-dimethyl substitution have been discovered to date, mainly ascribed to the methylation of the condensation product by C-methyltransferases. Recently, Keasling and coworkers demonstrated that modules of yersiniabactin and epothilone PKSs are capable of elongating the growing acyl-chain with dimethylmalonyl moieties ^28^, however disubstituted malonyl moieties have not yet been employed in directed biosynthesis. In order to incorporate the MeFC group into macrolides, we established a route for chemical synthesis of fluoromethylmalonyl-CoA (**17**, F-MM-CoA) following a similar strategy as for F-Mal-CoA (Fig. 4a). Finally, F-MM-CoA proved to be accepted by **H1** for elongation of the pentaketide (**9**) in presence of NADPH to produce 2-fluoro-10-deoxymethynolide (**18**) (for mechanistic implications, see Supplementary Fig. S14). Conversion of the extender unit was verified by the NADPH consumption assay (turnover rate: 0.14 ± 0.02 min^−1^) as well as HRMS, and reaction scale-up provided full structural analysis by NMR (Fig. 4b). Macrolactone **18** features a (2*S*,3*S*)-configuration identical to solithromycin (see Supplementary Note). The stereoselectivity of this reaction indicates that the DEBS M6-derived construct **H1** accommodates the F-MM moiety for substrate elongation with fluorine at the hydrogen position of the natively used methylmalonyl moiety (for formation of a cyclohexanone side product in absence of NADPH, see Supplementary Fig. S15). As a final step, we biotransformed the semisynthetic compound **18** for desosaminylation, and isolated the 2-fluoro derivative of YC-17 (**19**) (Supplementary Fig. S16), which demonstrated that post-PKS processing is not hindered by the fluorine atom. Using this approach, we have succeeded in introducing a fluorine atom into the macrolide antibiotic YC-17 and, with compound **18**, to create the precursor molecule for a plethora of fluorinated 12-membered ring macrolides with potential activity against erythromycin-resistant strains ^20^.

**Fig. 4:**
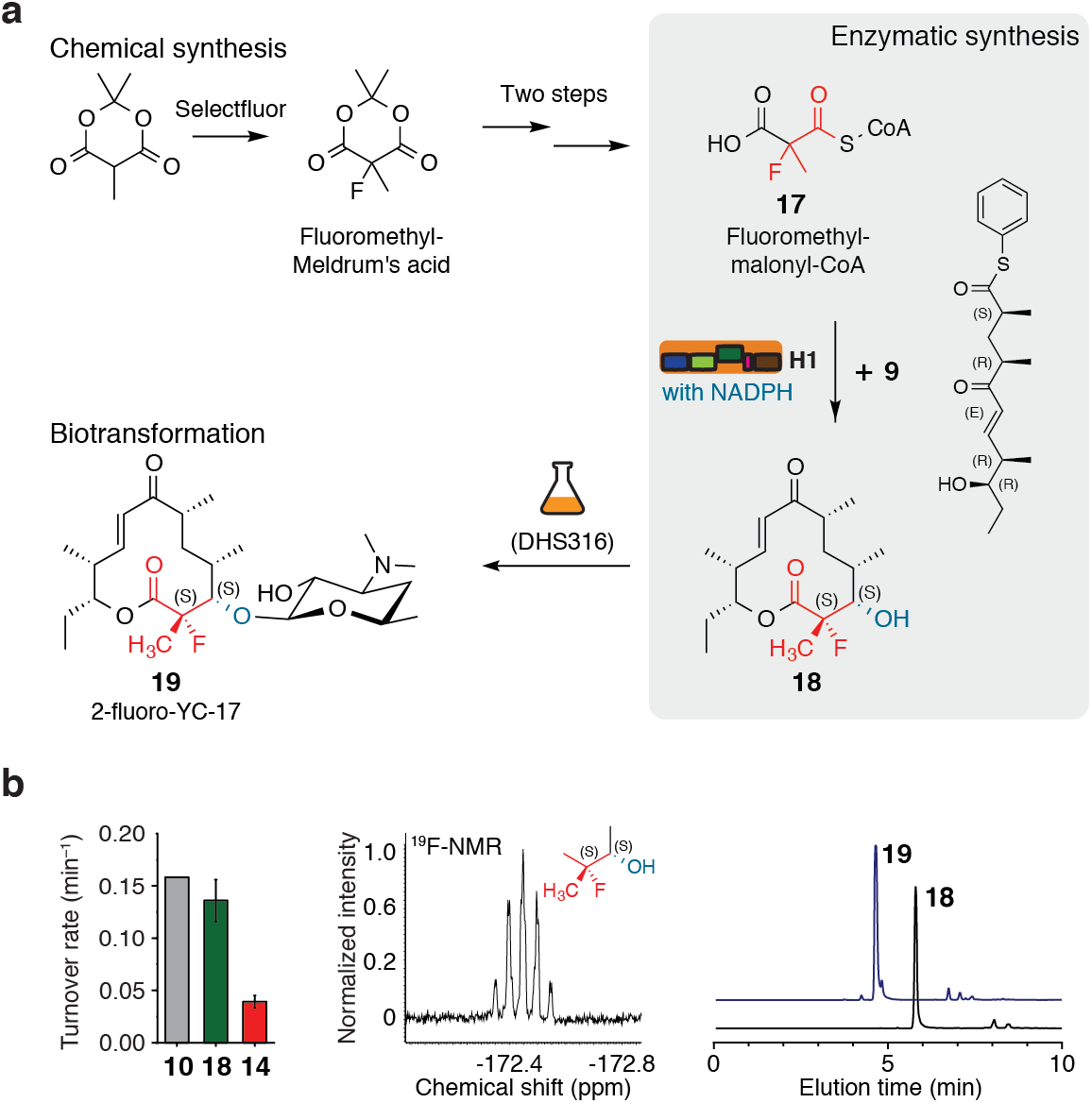
Synthesis of a new fluorinated macrolide antibiotic. **a**, Semi-synthetic approach to establish the fluoro-methyl disubstitution (MeFC group) at position C-2 in the macrolactone **10**. Chemical synthesis was performed analogously to **1** from the respective Meldrum’s acid and the product was converted enzymatically with pentaketide **9** and NADPH to macrolactone **18**. Compound **18** was transformed to the fluorinated derivative of the antibiotic YC-17 (**19**) using the strain DHS316. **b**, Selected data on target compounds and enzymatic turnover. Turnover rates for the **H1**-mediated conversion of F-MM-CoA, MM-CoA and F-Mal-CoA yielding compounds **18, 10** and **14**, respectively (left panel). Elongation using the substrate F-MM-CoA with subsequent reduction to compound **18** can be verified by the multiplicity in ^19^F-NMR as a quintet (middle panel). The conversion of compound **18** to 2-fluoro-YC-17 (**19**) by biotransformation was demonstrated by HPLC-HRMS (EIC: **18** [M+H]^+^ *m*/*z* = 315.1961; **19** [M+H]^+^ *m*/*z* = 472.3082 (right panel).

In conclusion, we present a new strategy to derivatize polyketides with fluorine by utilizing the promiscuous MAT domain of metazoan FAS, integrated as a domain in bacterial modular PKSs. This approach extends previous findings of Chang and coworkers, who introduced F-Mal-CoA metabolism in the cell ^11,12^. The AT exchange strategy maintains the overall protein architecture and integrity of the vectorial synthesis of type I PKSs, which paves the way for regioselective incorporation of fluorine in a diversity of polyketides during biosynthesis. Specifically, we demonstrate the relevance of this fluorination strategy by producing the new antibiotic 2-fluoro-YC-17 with the MeFC group selectively introduced in the *S*-configuration (stereochemistry as in solithromycin). Overall, this approach expands the toolbox for biosynthetic and semi-synthetic polyketide synthesis and offers a broadly applicable approach for the fluorination of complex PKS-derived macrolide compounds.

## Methods

Methods, additional references and spectra are available in the supplementary information.

### General synthesis of fluorinated CoA-Ester

Fluoro-Meldrum’s acid was synthesized from Meldrum’s acid in three steps using a previously described method utilizing Selectfluor^®^. Fluoromethyl-Meldrum’s acid was directly received from Methyl-Meldrum’s acid with Selectfluor^®^. The meldrum’s acids were treated with trimethylsilylthiophenol to produce the respective thiophenyl halfesters, and fluorinated CoA-Esters were eventually synthesized by transacylation from thiophenyl halfesters to free coenzyme A (CoASH). The CoA-Esters were purified by precipitation with acetone (−20 °C).

### General procedure for the biosyntheses

Small scale biosynthesis of TKLs and macrolactones were carried out with 4-6.1 µM enzyme, 5 mM **2** or 1 mM **9**, 200 µM X-CoA and 60 µM NADPH in the assay buffer (400 mM phosphate buffer, 20 % (v/v) glycerol, 1 mM EDTA, 0.8 % DMSO, pH 7.2) or the reaction buffer (250 mM potassium phosphate, 10 % glycerol, pH 7) at 25 °C. The reactions were followed fluorometrically by monitoring the consumption of NADPH. Products were extracted with EtOAc and confirmed by HPLC-MS.

### General procedure for the up-scaled syntheses of macrolactones

In order to receive larger amounts of macrolactones, reactions were carried out in 10-50 mL scale with the final concentrations of 5-10 µM **H1**, 300-600 µM **9**, 400-4000 µM X-CoA and 500-1000 µM NADPH in the reaction buffer at 25 °C. After at least 4 h of incubation, products were extracted with EtOAc and purified on a silica column. The macrolactones **15** and **16** were additionally purified by HPLC on a C18 column.

## Abbreviations

PKS: polyketide synthase
FAS: fatty acid synthase
(DEBS): 6-deoxyerythronolide synthase
AT: acyltransferase
MAT: malonyl-/acetyltransferase
ACP: acyl carrier protein
KS: -ketoacyl synthase
KR: ketoreductase
DH: dehydratase
ER: enoylreductase
TE: thioesterase
LD: linker domain
TKL: triketide lactone.

## Acknowledgments

This work was supported by a Lichtenberg grant of the Volkswagen Foundation to M.G. (grant number 85701). Further support was received from the LOEWE program (Landes-Offensive zur Entwicklung wissenschaftlich-ökonomischer Exzellenz) of the state of Hesse conducted within the framework of the MegaSyn Research Cluster. We would like to thank Khanh Vu Huu and Kudratullah Karimi for MS-analysis of acyl carrier proteins and Karthik S. Paithankar for proofreading the manuscript. Further, we are grateful to the Bode group for the extensive support in HPLC-MS analysis and Julia Wirmer-Bartoschek and Gabriele Sentis for support in NMR analysis. D.H.S. is grateful to NIH grant R35 GM118101 and the Hans W. Vahlteich Professorship for support.

## Funding

LOEWE program (Landes-Offensive zur Entwicklung wissenschaftlich-ökonomischer Exzellenz) of the state of Hesse (MG)

Lichtenberg grant of the Volkswagen Foundation (MG)

NIH grant R35 GM118101 (DHS)

## Author contributions

A.R. conceived and supervised the project. M.G. and D.S. designed the research. A.R. and D.H. performed the expression, purification and mutagenesis of murine KS-MAT constructs. L.M.M. performed global kinetic experiments and analyzed corresponding data under supervision of A.R.. A.R. and M.J. designed DEBS/FAS hybrids. M.J. performed the expression, purification and analysis of DEBS M6 constructs with respective MS analysis. M.J. and E.H. performed substrate consumption assays by HPLC-UV. F-Mal-CoA and F-MM-CoA were synthesized by A.R. and the diketide SNAC was synthesized by M.J. The pentaketide substrate was synthesized by J.J.S.. A.R. and M.J. performed semi-synthesis and analysis of compound 12, (14, 15), 16, and 18 and analyzed all data. J.J.S performed the biotransformation of compound 18. A.R., M.J., J.J.S., D.H.S. and M.G. wrote the manuscript.

## Competing interests

Authors declare that they have no competing interests.

## Data and materials availability

All data are available in the main text or the supplementary information.

## Supplementary Materials

Materials and Methods

Figs. S1 to S16

Tables S1 to S4

Supplementary Note

Supplementary Note Figures S1-S4

Supplementary Note Tables 1-2

Spectra

References

## Notes

### Competing Interest Statement

The authors have declared no competing interest.

