## Supplemental Information for "Directed biosynthesis of fluorinated polyketides"

**This PDF file includes:**

Materials and Methods
Figs. S1 to S16
Tables S1 to S4
Supplementary Note
Supplementary Note Figures S1-S4
Supplementary Note Tables 1-2
Spectra
References

#### 1. Materials

All CoA-esters (except F-Mal-CoA), β-$NAD^+$, NADH, α-ketoglutarate dehydrogenase (porcine heart) (αKGDH), α-ketoglutaric acid, thiamine pyrophosphate (TPP), and EDTA were purchased from Merck. BSA was from Serva. Restriction enzymes were bought from NEB biolabs. IPTG was from Carl Roth. Ni-NTA affinity resin was from Qiagen and 5 mL Strep-Tactin® columns were purchased from IBA technologies. Purity of CoA-esters was confirmed by HPLC-UV analysis before use.

Plasmid encoding DEBS M6+TE (pBL18) was kindly provided by the Khosla laboratory at Stanford University and the plasmid encoding the 4'-phosphopantetheinyl transferase Npt (UniProt code: A0A1Y2MXW0) was kindly provided by the Erb laboratory at the Max-Planck-Institute for terrestrial Microbiology, Marburg.

#### 2. General molecular biology experiments

##### 2.1 Cloning

Vectors encoding hybrid DEBS/FAS proteins (pMJD076 (**H2**) and pMJD077 (**H1**)) were produced by sequence and ligation independent cloning using the In-Fusion HD Cloning Kit (Takara Bio, USA). Briefly, pBL18 was amplified with primers: MJD101 and MJD102 or primers: MJD105 and MJD106. The corresponding inserts were generated by amplification of pAR2641 with primers: MJD087 and MJD088 or primers: MJD091 and MJD092. PCR products were treated with Dpn1 (NEB), purified by gel electrophoresis and DNA was extracted with the Wizard® SV Gel and PCR Clean-Up System (Promega). Purified vector and insert DNA were combined in the In-Fusion reaction following the manufacturer's instructions. DNA was transformed into E. coli Stellar™ Competent Cells, 5 mL LB cultures were grown and plasmids were isolated with the PureYield™ Plasmid Miniprep System (Promega) or the GeneJET Plasmid Miniprep Kit (Thermo Scientific). Sequences of all plasmids, listed in Table S3, were confirmed with the "dye terminator" method.

Vectors pAR432, pMJD091 and pMJD094 were also generated by sequence and ligation independent cloning using the In-Fusion HD Cloning Kit. For pAR432, the two fragments were obtained by amplification of pBL18 with primers: AR719 and AR722 or primers: AR721 and AR720. For pMJD091, the insert was generated by amplification of pET21a_Sppt with primers: MJD138 and MJD139 and the vector by amplification of pAR357 with primers: MJD136 and MJD137. For pMJD094, the insert was generated by amplification of pBL18 with primers: MJD145 and MJD146 and the linearized vector by digestion of pET-28a (Merck Millipore) with enzymes NdeI and EcoRI resembling the design of Kim *et al.*.[2]

##### 2.2 Heterologous expression and purification of murine KS-MAT

The plasmid was transformed into chemically competent *E. coli* BL21 Gold (DE3) cells. Transformed cells were grown overnight at 37 °C in 20 mL LB (100 µg $mL^{-1}$ ampicillin (amp) and 1 % (w/v) glucose) medium. Pre-cultures were used to inoculate 1 L TB medium (100 µg $mL^{-1}$ amp). Cultures were grown at 37 °C until they reached an optical density ($OD_{600}$) of 0.5–0.6. After cooling at 4 °C for 20 min, cultures were induced with 0.25 mM IPTG, and grown for additional 16 h at 20 °C and 180 rpm. Cells were harvested by centrifugation (4,000 rcf for 20 min). The cell pellets were resuspended in lysis buffer (50 mM potassium phosphate, 200 mM potassium chloride, 10 % (v/v) glycerol, 1 mM EDTA, 30 mM imidazole (pH 7.0)) and lysed by French press. After centrifugation at 50,000 rcf for 30 min, the supernatant was mixed with 1 M $MgCl_2$ to a final concentration of 2 mM. The cytosol was transferred to Ni-NTA-columns and washed with 5 column volumes (CV) wash buffer (lysis buffer without EDTA). Bound protein was eluted with 2.5 CV elution buffer (50 mM potassium phosphate, 200 mM potassium chloride, 10 % (v/v) glycerol, 300 mM imidazole (pH 7.0). The eluent was transferred to Strep-Tactin-columns, and washed with 5 CV strep-wash buffer (250 mM potassium phosphate, 10 % (v/v) glycerol, 1 mM EDTA, (pH 7.0)). Proteins were eluted with 2.5 CV elution buffer (strep-wash buffer containing 2.5 mM D-desthiobiotin). After concentration to 10–20 mg $mL^{-1}$, the proteins were frozen in liquid nitrogen and stored at -80 °C. Samples were thawed at 37 °C for 30 min and further polished by size-exclusion chromatography (SEC) using a Superdex 200 GL10/300 column equilibrated with the strep-wash buffer. Fractions containing dimeric protein were pooled and concentrated to 10-20 mg $mL^{-1}$ to be frozen in liquid nitrogen and stored in aliquots at –80 °C.

##### 2.3 Heterologous expression and purification of FAS ACP and DEBS ACP6

FAS ACP for the activity assay was produced by co-expressing FAS ACP with Sfp from *Bacillus subtilis* bicistronically (pAR352) in *E. coli* BL21gold(DE3) cells.[1] Overnight cultures were grown in LB (100 µg/mL ampicillin and 1 % glucose) at 37 °C. 2 L TB medium (100 µg/mL amp) was inoculated and incubated at 37 °C until an optical cell density ($OD_{600}$) of 0.5-0.6 was reached. After cooling at 4 °C for 20 min, cultures were induced with 0.25 mM IPTG and cells were grown for 16 h at 20 °C. Cells were harvested by centrifugation, resuspended in lysis buffer (50 mM sodium phosphate, 200 mM NaCl, 20 % (v/v) glycerol, 1 mM EDTA, 30 mM imidazole (pH 7.4) and lysed by French press. After centrifugation (50,000 rcf for 30 min), the supernatant (supplemented with 2 mM $MgCl_2$) was transferred to Ni-NTA-columns and washed with 5 CV wash buffer (lysis buffer without EDTA). The protein was eluted with 2.5 CV elution buffer (wash buffer containing 300 mM imidazole) and concentrated. Pooled fractions, were separated on a Superdex 200 HiLoad 16/60 or 26/60 SEC column equilibrated with buffer (50 mM sodium phosphate, 200 mM NaCl, 10 % (v/v) glycerol, 1 mM EDTA, 1 mM DTT). All fractions containing monomeric ACP were pooled and concentrated to 10-20 mg $mL^{-1}$ (common yield was 20 mg purified protein per 1 L culture).

DEBS ACP6 was produced by co-expressing DEBS ACP6 (pMJD094) with Sfp from *Bacillus subtilis* (pAR357) or with 4'-phosphopantetheinyl transferase Npt from *Streptomyces platensis* (pMJD091) in *E. coli* BL21gold(DE3) cells. Expression and purification was performed with the FAS ACP protocol, except that kanamycin (50 µg/mL kan) and spectinomycin (50 µg/mL spc) were used as antibiotics for cultivation. Phosphopantetheinylation of DEBS ACP6 was investigated by electrospray ionization (ESI) mass spectrometry (Figure S4), which showed that only co-expression with Npt yielded sufficiently activated protein. All assays were performed with this protein.

##### 2.4 Heterologous expression and purification of DEBS M6+TE, its variants and DEBS $KS^{C1661G}$-AT

DEBS M6+TE and all its variant were produced by co-expression of DEBS M6+TE (pBL18, pMJD076, pMJD077) with Sfp from *Bacillus subtilis* (pAR357) in *E. coli* BL21gold(DE3) cells. Expression and purification was performed with the FAS ACP protocol, except that ampicillin (100 µg/mL amp) and spectinomycin (50 µg/mL spc) were used as antibiotics for cultivation. Further, the protein was purified in a different lysis buffer (50 mM sodium phosphate, 450 mM NaCl, 20 % (v/v) glycerol, 1 mM EDTA, 20 mM imidazole (pH 7.6)). The protein on the Ni-NTA column was washed with 5 CV washing buffer 1 (50 mM sodium phosphate, 450 mM NaCl, 20 % (v/v) glycerol, 20 mM imidazole (pH 7.6)) and 2 CV washing buffer 2 (50 mM sodium phosphate, 450 mM NaCl, 20 % (v/v) glycerol, 1 mM EDTA, 60 mM imidazole (pH 7.6)). The proteins were eluted with 2.5 CV elution buffer (50 mM sodium phosphate, 450 mM NaCl, 20 % (v/v) glycerol, 300 mM imidazole (pH 7.6)) and further polished and analyzed by size-exclusion chromatography (SEC) using a Superdex 200 Increase 10/300 GL, HiLoad 16/600 Superdex 200 or Superose 6 Increase 10/300 GL column equilibrated with the washing buffer 1. Fractions containing dimeric protein were pooled and concentrated to 1–10 mg $mL^{-1}$ to be frozen in liquid nitrogen and stored in aliquots at –80 °C.

DEBS $KS^{C1661G}$-AT (pAR432) was expressed without Sfp in *E. coli* BL21gold(DE3) cells and purified following the protocol for the whole DEBS module 6.

2.5 <u>Protein concentration</u>

Protein concentrations were calculated from the absorbance at 280 nm, which was recorded on a NanoDrop 2000c (Thermo Scientific). Extinction coefficients were calculated from the primary sequence without *N*-formylmethionine with CLC Main workbench (Qiagen). Absorbance 1 g/L at 280 nm (10 mm): 1.053 for FAS KS-MAT; 0.475 for FAS ACP, 1.009 for DEBS M6+TE (**WT**), 0.474 for DEBS ACP6, 0.899 for DEBS KS6-AT6, 1.087 for **H1** and 1.069 for **H2**.

2.6 <u>Thermal shift assay</u>

Thermal shift assays were performed as previously reported.[3] Briefly, 2 µL of protein solution (5-6 µM) were mixed with 21 µL of the respective buffer and 2 µL of SYPRO Orange protein gel stain (80 × diluted), then fluorescence was measured from 5 °C to 95 °C with a step gradient of 0.5 °C $min^{-1}$, with excitation wavelength set to 450-490 nm, and emission wavelength to 560-580 nm. Data was analyzed with the software CFX Maestro 1.0.

2.7 <u>α-Ketoglutarate dehydrogenase coupled activity assay</u>

The enzyme-coupled assay was performed as previously published.[1] Assays were run in 384-well Small Volume HiBase Microplates (Greiner Bio-one) with following settings for the microplate reader (ClarioStar, BMG labtech): 348-20 nm; emission: 476-20 nm; gain: 1500; focal height: 11.9 mm; flashes: 17; orbital averaging: off.

Briefly, four different solutions were prepared as 4-fold concentrated stocks in assay buffer (50 mM sodium phosphate, 10 % (v/v) glycerol, 1 mM EDTA (pH 7.6), filtered and degased). Solution 1 (Sol 1) contained the acyltransferase, supplemented with 0.1 mg $mL^{-1}$ BSA. Solution 2 (Sol 2) contained 8 mM α-ketoglutaric acid, 1.6 mM $NAD^+$, 1.6 mM TPP and 60 mU/100 µL αKGDH. Solution 3 (Sol 3) contained the CoA-esters and Solution 4 (Sol 4) finally contained the respective ACP as standalone protein. The components (5 µL) were pipetted in order: Sol 1, Sol 2 and Sol 3, followed by manual mixing. The transfer reaction was initiated by injection of 5 µL Sol 4 with the dispenser. The final concentrations of all ingredients were 50 mM sodium phosphate, pH 7.6, 10 % (v/v) glycerol, 1 mM EDTA, 2 mM α-ketoglutaric acid, 0.4 mM $NAD^+$, 0.4 mM TPP, 15 mU/100 µL αKGDH, 0.03 mg $mL^{-1}$ BSA, 1-5 nM FAS MAT or 250 nM DEBS AT6, 10–400 µM ACP, 0.1–25 µM X-CoA for FAS MAT and 2-130 µM for DEBS AT6 (where X refers to the respective acyl-moiety of the assay). Equidistant kinetic measurements were taken every 5 s for 5 min at 25 °C. Every data point was recorded in technical triplicates and the respective background noise of the assay set-up (assay buffer supplemented with 0.1 mg $mL^{-1}$ BSA) was subtracted. The enzyme-mediated hydrolysis rate was not subtracted, as it is relatively low compared to transfer rates.

2.8 Analysis of AT-mediated transfer by global fitting

For every global fit, initial velocities were determined for eight different CoA-ester concentrations at four fixed ACP concentrations. Every data point reflects the result from one biological replicate and measurements were performed in at least two biological replicates ($n \geq 2$). Relative fluorescent units were converted into concentrations using a NADH calibration curve. Series of response curves were globally fit using all data without any parameter constraints. The global fit was performed with OriginPro 8.5 (OriginLab, USA) using the following equations for the ping-pong mechanism:

$$\boldsymbol{v} = \frac{\boldsymbol{k}_{\mathbf{cat}}[\mathbf{AT_0}][\mathbf{XCoA}][\mathbf{ACP}]}{[\mathbf{XCoA}]\boldsymbol{K}_{\mathbf{ACP}} + [\mathbf{ACP}]\boldsymbol{K}_{\mathbf{S}} + [\mathbf{XCoA}][\mathbf{ACP}]} \qquad [\ 1\ ]$$

2.9 Specific ketoreductase activity

The specific ketoreductase activity of DEBS KR6 was measured fluorometrically by monitoring the consumption of NADPH at 25 °C according to previous reports.[4] Assays were performed in 384-well Small Volume HiBase Microplates (Greiner Bio-one) with following settings for the microplate reader (ClarioStar, BMG labtech): 348-20 nm; emission: 476-20 nm; gain: 1301; focal height: 12.4 mm; flashes: 17; orbital averaging: off. Two solutions were prepared as 4-fold concentrated stocks and *trans*-1-decalone was prepared as 2-fold concentrated stock, in the assay buffer (400 mM phosphate buffer, 20 % (v/v) glycerol, 2 mM DTT, 1 mM EDTA, 0.8 % DMSO (pH 7.2)).[5] Solution 1, 2 and 3 contained the variants of DEBS M6+TE (1.2 µM), the NADPH (240 µM) and the *trans*-1-decalone (4 mM in assay buffer), respectively. 5 µL of enzyme and NADPH solution were mixed with 10 µL of *trans*-1-decalone solution (all incubated at 25 °C) to final concentrations of 0.3 µM enzyme, 60 µM NADPH and 2 mM *trans*-1-decalone. The consumption was monitored fluorometrically for 3 min and converted to concentrations using a NADPH calibration curve. Each of the biological triplicates was measured in technical replicates and the slope was corrected by the background noise of the assay set up (NADPH and *trans*-1-dealone in the respective concentrations without enzyme).

#### 2.10 NADPH consumption assays following reduced triketide lactone (TKL) and reduced macrolactone production

The production rate of reduced TKLs and macrolactones were monitored fluorometrically by observing the consumption of NADPH. Assays were performed in 384-well Small Volume HiBase Microplates (Greiner Bio-one) with following settings for the microplate reader (ClarioStar, BMG labtech): 348-20 nm; emission: 476-20 nm; gain: 1301; focal height: 12.4 mm; flashes: 17; orbital averaging: off. Four solutions were prepared as 4-fold concentrated stocks and the X-CoA and NADPH solutions were combined to yield a 2-fold concentrated stock, in the assay buffer (400 mM phosphate buffer, 20 % (v/v) glycerol, 1 mM EDTA, 0.8 % DMSO (pH 7.2)). Solution 1, 2 and 3 contained the variants of DEBS M6+TE (16 µM), the diketide SNAC **2** (20 mM) or pentaketide **9** (4 mM), MM-CoA or Mal-CoA or F-Mal-CoA (400 µM) and the NADPH (120 µM), respectively. 5 µL of priming substrates and enzyme solutions were mixed with 10 µL of the X-CoA, NADPH stock (all incubated at 25 °C) to final concentrations of 4 µM enzyme, 5 mM **2** or 1 mM **9**, 200 µM X-CoA and 60 µM NADPH. The fluorescence was monitored for 13-20 min and converted into concentrations by using a NADPH calibration curve. Each of the biological triplicates was measured in technical replicates and the slope was corrected by the background noise of the assay set up (NADPH, X-CoA and **2** without enzyme for TKL production and NADPH, **9** and enzyme without elongation substrate for macrolactones). For analyzing the turnover rate of FMM-CoA with H1 and pentaketide **9** and to compare them with MM-CoA and F-Mal-CoA conditions were slightly changed to 6.4 µM H1 and to the buffer composition (250 mM potassium phosphate, 10 % glycerol, pH 7). Assays with these conditions are highlighted with an asterisk in table S2.

#### 2.11 Chromatographic assay following non-reduced TKL production

Alternatively, we measured the production rates for non-reduced TKLs in the absence of NADPH by quantifying the consumption of CoA-ester substrates and liberation of CoA with HPLC-UV. The four stock solutions were prepared as described in the previous section in the assay buffer (400 mM phosphate buffer, 20 % (v/v) glycerol, 1 mM EDTA, 0.8 % DMSO (pH 7.2)). 80 µL of all solutions were mixed and incubated at 25 °C for 60 min. After 0, 5, 10, 15, 20, 30 and 60 min, 40 µL samples were taken and the reactions were stopped by the addition of 5 µL perchloric acid (70 %). Reaction mixtures were mixed thoroughly and subsequently neutralized with 5 µL NAOH (10 M). Hydoxybutyryl-CoA (final concentration: 50 µM) was added as an internal standard. Reaction mixtures were spun at 20000 rcf for 10 min. 30 µL of the supernatant was transferred into HPLC vials and measured by HPLC-UV.

HPLC-UV analysis of CoA esters was performed using a Dionex UltiMate 3000 RSLC with a UV detector. Chromatographic separation was performed on a Synchronis aQ-c18 column (4.6 × 250 mm, particle size 5 µm, ThermoFisher Scientific) with a mobile-phase system consisting of buffer A (0.2 M ammonium acetate (pH 6.0)) and buffer B (methanol). The column was equilibrated with 95 % buffer A and 5 % buffer B at a flow rate of 0.8 mL/min. X-CoA was purified using a linear gradient from 5 to 20 % B over 15 min and an increased gradient from 20 to 60 % B over 8 min (The column was regenerated 3 min at 60 % B and re-equilibrated again for the next sample). Absorbance of X-CoA was monitored at 260 nm and assigned by the elution time of purchased CoA ester references. Peak areas were correlated to the internal standard peak and converted into concentrations. Concentrations were plotted versus the time and fit with an exponential decay function in OriginPro 8.5 (OriginLab, USA). Initial velocities were obtained from the deviation of the function and its slope at 0 min. Background measurements were performed in the absence of diketide SNAC **2**.

$$y = y_0 + A_1 e^{-x/t_1} \qquad [2]$$

2.12 HPLC-MS analysis of TKL and macrolactone products

All reaction mixtures were extracted using EtOAc (2 times 400 µL for TKLs, 3 times 300 µL for macrolactones). Combined organic phases were evaporated in a SpeedVac *in vacuo*. Samples were dissolved in 50 µL methanol and spun at 20000 rcf for 20 min. 40 µL of the supernatant was transferred into HPLC vials and measured by HPLC-MS using the Ultimate 3000 LC (Dionex) system connected to a Acquity UPLC BEH C18 (2.1 × 50 mm, particle size 1.7 µm, Waters) for separation. After equilibration with 5 % acetonitrile in water, samples were purified using a linear gradient from 5-95 % within 16 min and subsequently injected into the AmaZonX (Bruker) or Impact II qTof (Bruker) for ESI.

2.13 HPLC-UV analysis and purification of macrolactone products

HPLC-UV analysis of macrolactones was performed using a Dionex UltiMate 3000 RS UHPLC with a RS Diode Array UV detector. Chromatographic separation was performed on a Synchronis aQ-c18 column (4.6 × 250 mm, particle size 5 µm, ThermoFisher Scientific) with a mobile-phase system consisting of buffer A (water) and buffer B (acetonitrile). The column was equilibrated with 95 % buffer A and 5 % buffer B at a flow rate of 0.8 mL/min. Macrolactones were purified using a linear gradient from 5 to 95 % B over 15 min and another 5 min at 95 % B. A linear gradient to 5 % B was used within 1 min and the column was re-equilibrated for five more minutes at 5 % B. Fractionation was performed with the UltiMate 3000 Autosampler in between 3 and 16 min in 20 s steps.

2.14 Enzymatic synthesis and analysis of compound **12**

In order to conduct NMR analysis of compound **12**, the reaction volume was scaled up to 10 mL and a purification strategy was established. Final concentrations of 300 µM pentaketide substrate **9** (1.05 mg, dissolved in DMSO), 400 µM Mal-CoA and 500 µM NADPH were dissolved in assay buffer (250 mM potassium phosphate, 10 % glycerol, pH 7) and 5 µM **H1** (9.16 mg, dissolved in 250 mM potassium phosphate, 10 % glycerol, pH 7) was added to a final volume of 10 mL. The reaction mixture (slightly cloudy emulsion) was incubated for 16 h at 25 °C and the progress of the reaction was monitored by HPLC (samples were prepared by quenching 20 µL mixture with 60 µL methanol). The reaction mixture was transferred to a 50 mL falcon tube and the aqueous phase was extracted with EtOAc (5 × 10 mL) by spinning for 1 min at 3000 rcf to separate the phases. The combined organic phases were transferred to a 250 mL round bottom glass flask, the solvent was evaporated *in vacuo* and the residual oil was dried by azeotropic evaporation with toluene (2 × 1 mL). After running a TLC (hexane/EtOAc 1:1 stained with $KMnO_4$-solution), the crude product was adsorbed to silica gel and purified by flash chromatography (silica column: 1.3 cm diameter and 15 cm height). A gradient was used for elution (hexane:EtOAc 20:1 → 10:1 → 7:1 → 5:1 → 3:1 → 2:1 → 1:1) and every fraction was carefully analyzed by TLC. All fractions containing the product were pooled, filtered and the solvent was evaporated *in vacuo* to yield compound **12** as white solid.

**$R_F$-value** (TLC; hexane/EtOAc: 1/1): 0.44
**Retention time** (HPLC; water/ACN): 13.74 min

**$^{1}$H NMR** (600 MHz, Chloroform-*d*) δ 6.76 (dd, *J* = 15.7, 5.4 Hz, 1H), 6.44 (dd, *J* = 15.8, 1.2 Hz, 1H), 5.02 (ddd, *J* = 8.3, 6.0, 2.3 Hz, 1H), 4.05-4.02 (ddd, *J* = 11.2, 5.2, 1.2 Hz, 1H), 2.71-2.54 (m, 4H), 1.74-1.56 (m, 4H), 1.30-1.28 (m, 1H), 1.25 (d, J = 7.0 Hz, 3H), 1.13 (d, J = 6.9 Hz, 3H), 1.03 (d, J = 6.2 Hz, 3H), 0.93 (t, J = 7.4 Hz, 3H) ppm.
**$^{13}$C NMR** (500 MHz, Chloroform-*d*) δ 170.3, 147.5, 125.5, 74.5, 72.9, 45.3, 39.0, 38.1, 33.0, 32.5, 25.0, 17.7, 17.2, 10.3, 9.4 ppm.
**MS (HR-ESI+)** found 283.1900, for $[M+H]^+$ calculated 283.1910; found 305.1720, calculated for $[M+Na]^+$ 305.1729; found 265.1796, calculated for $[M-H_2O+H]^+$ 265.1804.

2.15 Enzymatic synthesis and analysis of compound **15/16**

In order to conduct NMR analysis of compound **15**/**16**, the reaction volume was scaled up to 50 mL and a purification strategy was established. Final concentrations of 600 µM pentaketide substrate **9** (10.46 mg, dissolved in DMSO), 4000 µM F-Mal-CoA and 1000 µM NADPH were dissolved in assay buffer (250 mM potassium phosphate, 10 % glycerol, pH 7) and 10 µM **H1** (90.57 mg, dissolved in 250 mM potassium phosphate, 10 % glycerol, pH 7) was added to a final volume of 50 mL. The reaction mixture (slightly cloudy emulsion) was incubated for 22.5 h at 25 °C and the progress of the reaction was monitored by HPLC (samples were prepared by quenching 20 µL mixture with 60 µL methanol). The reaction mixture was transferred to two 50 mL falcon tube and the aqueous phase was extracted with EtOAc (4-6 × 25 mL) by spinning for 1 min at 3000 rcf to separate the phases. The combined organic phases were transferred to a 250 mL round bottom glass flask, the solvent was evaporated *in vacuo*. The residual oil was dissolved in MeOH/CHCl3 (9/1) and purified with an uHPLC system (Thermo Fisher) on a Synchronis aQ-C18-LC column. Fractions containing the major fluorinated compound, based on $^{19}$F-NMR analysis, were further purified with flash chromotagraphy. The crude product was adsorbed to silica gel and purified (silica column: 1.3 cm diameter and 12 cm height) using a gradient for elution (hexane:EtOAc 20:1 → 15:1 → 10:1 → 7:1 → 5:1 → 4:1 → 3:1 → 1:1) and every fraction was carefully analyzed by TLC. All fractions containing product were pooled, filtered and the solvent was evaporated *in vacuo* to yield compound **16** as white solid.
**$R_F$-value** (TLC; hexane/EtOAc: 3/1): 0.22
**Retention time** (HPLC; water/ACN): 13.23 min
**$^{1}$H NMR** (500 MHz, Chloroform-*d*) δ 5.70 (dd, *J* = 15.6, 8.0 Hz, 1H), 5.51 (dd, *J* = 15.7, 0.8 Hz, 1H), 4.84 (d, *J* = 48.4 Hz, 1H), 3.41 (ddd, *J* = 9.0, 5.5, 3.7 Hz, 1H), 2.54-2.47 (m, 1H), 2.41-2.34 (m, 1H), 2.07-2.00 (m, 1H), 1.92-1.87 (m, 1H), 1.66 (q, *J* = 12.8 Hz, 1H), 1.61-1.55 (m, 1H), 1.44-1.35 (m, 1H), 1.11 (d, *J* = 6.4 Hz, 3H), 1.08 (d, *J* = 6.8 Hz, 3H), 0.98-0.95 (ovlp m, 6H)
**$^{19}$F NMR** (500 MHz, Chloroform-*d*) δ -206.21(d, *J* = 46,8 Hz)
**$^{13}$C NMR** from HSQC (500 MHz, Chloroform-*d*) δ 134.1, 131.5, 94.8/96.4, 76.5, 42.4, 42.2, 38.7, 36.9, 27.0, 13.4, 15.2, 10.4, 13.6
**MS (HR-ESI+)** found 273.1859, for $[M+H]^+$ calculated 273.1866; found 295.1677, calculated for $[M+Na]^+$ 295.1686; found 255.1754, calculated for $[M-H_2O+H]^+$ 255.1761; found 567.3462, calculated for $[M_2+Na]^+$ 567.3474

2.16 Enzymatic synthesis and analysis of compound **18**

In order to conduct NMR analysis of compound **18**, the reaction volume was scaled up to 50 mL and a purification strategy was established. Final concentrations of 600 µM pentaketide substrate **9** (10.46 mg, dissolved in DMSO), 800 µM F-MM-CoA and 1000 µM NADPH were dissolved in assay buffer (250 mM potassium phosphate, 10 % glycerol, pH 7) and 7 µM **H1**

(63.40 mg, dissolved in 250 mM potassium phosphate, 10 % glycerol, pH 7) was added to a final volume of 50 mL. The reaction mixture (slightly cloudy emulsion) was incubated for at least 4 h at 25 °C and the progress of the reaction was monitored by HPLC (samples were prepared by quenching 20 µL mixture with 60 µL methanol). The reaction mixture was transferred to two 50 mL falcon tube and the aqueous phase was extracted with EtOAc (5 × 25 mL) by spinning for 1 min at 3000 rcf to separate the phases. The combined organic phases were transferred to a 250 mL round bottom glass flask, the solvent was evaporated *in vacuo*. The residual oil was purified with flash chromotagraphy. After running a TLC (hexane/EtOAc 2:1 stained with $KMnO_4$-solution), the crude product was adsorbed to silica gel and purified (silica column: 1.3 cm diameter and 12 cm height) using a gradient for elution (hexane:EtOAc 30:1 → 15:1 → 10:1 → 7:1 → 5:1 → 4:1 → 3:1 → 2:1 → 1:1) and every fraction was carefully analyzed by TLC. All fractions containing the product were pooled, filtered and the solvent was evaporated *in vacuo* to yield compound **18** as white solid. The educt could not be separated by this procedure as co-eluting with compound **18**.

**$R_F$-value** (TLC; hexane/EtOAc: 3/1): 0.83

**Retention time** (HPLC; water/ACN): 14.65 min

**$^1$H NMR** (500 MHz, Chloroform-*d*) δ 6.78 (dd, *J* = 15.8, 5.4 Hz, 1H), 6.49 (dd, *J* = 15.8 ,1.2 Hz, 1H), 5.09 (ddd, 8.0, 5.4, 1.8 Hz, 1H), 3.68 (dd, *J* = 26.3, 0.9 Hz, 1H), 2.73-2.66 (m, 1H), 2.57-2.48 (m, 1H), 2.03-1.98 (m, 1H), 1.85-1.77 (m, 1H), 1.69-1.62 (m, 1H), 1.66 (d, *J* = 22 Hz, 3H), 1.34-1.25 (m, 2H), 1.22 (d, *J* = 7 Hz, 3H), 1.19 (d, *J* = 6.9 Hz, 3H), 1.04 (d, *J* = 6.7 Hz, 3H), 0.96 (t, *J* = 7.3 Hz, 3H)

**$^{19}$F NMR** (500 MHz, Chloroform-*d*) δ -172.45 (quintet, *J* = 22,5 Hz)

**$^{13}$C NMR** from HSQC δ 146.0, 125.9, 78.2, 75.2, 45.0, 37.7, 32.7, 31.8, 25.1, 22.7, 17.7, 17.5, 10.2, 9.3

**MS (HR-ESI+)** found 315.1961, for $[M+H]^+$ calculated 315.1972; found 337.1781, calculated for $[M+Na]^+$ 337.1791; found 297.1857, calculated for $[M-H_2O+H]^+$ 297.1866

**Yield:** 6%, note on the yield: Without any optimization, we yielded about 6% of the fluorinated 10-deoxymethynolide (**18**) utilizing shortened and non-canonical starter substrate. Recently, a similar yield was reported for the elongation of the pentaketide (**9**) with the native MM-CoA by wildtype DEBS M6, and the TE was identified as the bottleneck hindering macrolactonization.[6] These results indicate that the AT domain exchange is not responsible for the low yield.

##### 2.17 Biotransformation of compound 18

The following biotransformation reaction was carried out in an analogous manner to the published procedures with minor modifications.[7,8] A 3 mL seed culture of SCM medium (20 g soytone, 15 g soluble starch, 10.5 g MOPS, 1.5 g yeast extract, 0.1 g $CaCl_2$, per 1 L water, pH 7.2 ) in 15 mL snap cap tube was inoculated with 3 µL of *Streptomyces venezuelae* strain DHS316 spore stock and shaken overnight at 28 °C (180 rpm). The OD600 was measured and this was used to inoculate a 10 mL biotransformation culture of SCM medium to an OD600 of ~0.1. The culture was incubated at 28 °C (180 rpm) for 1h, prior to the addition of 2 µL of acetyl-narbonolide as a DMSO solution (20 mg/mL) followed by the macrolactone **18** as a DMSO solution (~75 µL). The cultures were incubated at 28 °C for 18 h (180 rpm) and centrifuged at 4,000 x g for 10 min to remove cell debris. The remaining aqueous solution was saturated with NaCl prior to adjusting the pH to 11 with 10N NaOH. The solution was extracted with 3 x 10 mL of ethyl acetate, and the combined organic layers were dried over anhydrous $Na_2SO_4$. Solvent was removed under reduced pressure to yield crude product mixture. The reactions were purified by HPLC using a HydroRP C18 column, 250 x 10.0 mm, 4 micron, with 3 mL/min flow rate. A gradient of solvent A (Water + 0.1% Formic Aicid) and solvent B (Acetonitrile) was as follows: isocratic 10% B for 2 minutes, then a linear gradient from 10% - 95% B over 38 minutes, then isocratic 95% for 10 minutes. Product eluted between 11.5 – 13.5 minutes.

**MS (HR-ESI+)** found 472.3082, for $[M+H]^+$ calculated 472.3069; found 494.2877, calculated for $[M+Na]^+$ 494.2888

#### 3. Substrate synthesis

3.1 Chemicals

Reagents were purchased from Merck, Carl-Roth and abcr GmbH. All the reagents were used as purchased without any further purification. Reactions were carried out in oven-dried glassware under an inert Argon gas atmosphere. Reactions were magnetically stirred and monitored by thin layer chromatography (TLC) on 0.25 mm E. Merck silica gel plates (60F-254) using UV light as visualizing agent and $KMnO_4$ stain & heat as developing agents. Room temperature when mentioned ranges from 22 to 25 °C. E. Merck silica gel (60, particle size 0.040-0.060mm) was used for column chromatography. The NMR spectra were determined on Bruker DPX250, on Bruker AV400, on Bruker AV500 or on Bruker DRX600 using deuterated solvents from the company Deutero GmbH, Kastellaun. Chemicals shifts values are expressed in parts per million (ppm) relative to chloroform (δ 7.27), DMSO (δ 2.51) or water (δ 4.79). Multiplicities are explained using following abbreviations: s = singlet, d = doublet, t = triplet, q = quartet, m = multiplet, dd = doublet of doublet, dq = doublet of quartet, ddd = doublet of doublet of doublet. High resolution Electrospray Ionization mass spectra were obtained on ThermoFisher Surveyor MSQ.

3.2 Fluoromalonyl-CoA

Fluoro-Meldrum's acid (S1)

Fluoro-Meldrum's acid (S1) was synthesized from Meldrum's acid in three steps using a previously described method.[9]

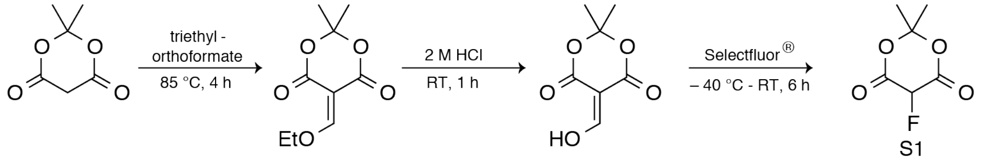

**Yield** (over three steps): 44 % (5.1 g), colorless solid.

**$^1$H NMR** (400 MHz, $C_2D_6OS$) δ = 6.69 (d, *J* = 41.6 Hz, 1H), 1.82 (s, 3H), 1.71 (s, 3H) ppm.

**$^{19}$F NMR** (300 MHz, $C_2D_6OS$) δ = -207.9 (d, *J* = 42 Hz, 1H) ppm.

Fluoromalonic acid thiophenyl halfester (S2)

The thiophenyl halfester (S2) was synthesized from fluoro-Meldrum's acid S1 and trimethylsilylthiophenol, which was produced utilizing the butyllithium method.[9]

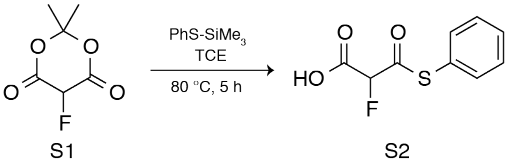

**Yield**: 53 % (0.7 g), yellowish powder.

**$^1$H NMR** (400 MHz, $C_2D_6OS$) δ = 7.64-7.50 (m, 3H), 7.49-7.46 (m, 2H), 5.95 (d, *J* = 47.2 Hz, 1H) ppm.

**$^{19}$F NMR** (300 MHz, $C_2D_6OS$) δ = -192.5 (d, *J* = 48 Hz) ppm.

Fluoromalonyl-CoA (1)

Fluoromalonyl-CoA (1) was synthesized via transacylation from the fluoromalonyl thiophenyl halfester S2 to free coenzyme A (CoASH), adapted from Dunn *et al.*[10]

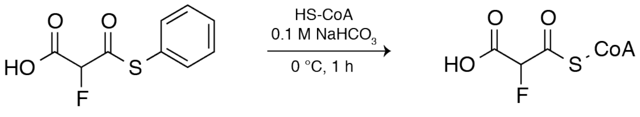

S2 **1**

CoASH (30 mg; 38 µmol) and the fluoromalonyl thiophenyl halfester (44 mg; 205 µmol) were dissolved in 1 mL ice cold $NaHCO_3$ (0.1 M) at 0 °C. The pH was adjusted back to pH 8-9 with 1 mL NaOH (0.2 M) and the reaction was continued for 1 h at 0 °C. Then, the reaction mixture (2 mL) was poured into 15 mL acetone (-20 °C) in a 50 mL falcon and spun for 10 min at 3000 rcf. The supernatant was discarded and the pellet was washed with acetone (-20 °C) and dried by air stream. The pellet was dissolved in water and the yield was determined by the absorbance at 260 nm (ε = 16,400 $M^{-1}$ $cm^{-1}$).[11]

**Yield**: 81 %, white crystals.
**$^1$H NMR** (400 MHz, $D_2O$) δ = 8.54 (s, 1H), 8.25 (s, 1H), 6.18-6.13 (m, 1H), 5.33 (d, *J* = 50.3 Hz, 1H), 4.83-4.79 (m, 2H), 4.58-4.53 (m, 1H), 4.25-4.17 (m, 1H), 3.99 (s, 1H), 3.83-3.76 (m, 1H), 3.55-3.47 (m, 1H), 3.47-3.39 (m, 2H), 3.37-3.30 (m, 2H), 3.10-3.03 (m, 2H), 2.47-2.37 (m, 2H), 0.85 (s, 3H), 0.70 (s, 3H) ppm.
**$^{19}$F NMR** (300 MHz, $D_2O$) δ = -177.4 (d, *J* = 54 Hz), -182.0 (M) ppm.
**HRMS (ESI+)** found 872.1138, calculated for $[M+H]^+$ 872.1141.

3.3 Diketide SNAC (2)

(4*S*, 2’*S*, 3’*R*)-3-(2’-Methyl-3’-hydroxypentanoyl)-4-benzyl-2-oxazolidinone (S4)

Compound S4 was synthesized from Evans auxiliary S3 using the method of Sharma *et al.*.[12] The product was triturated with EtOAc, *n*-hexane and *n*-pentane after column chromatography.

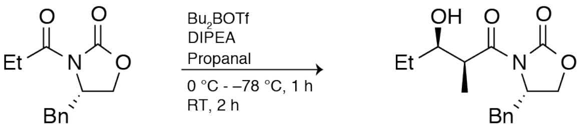

S3 S4

**Yield**: 69 % (4.3 g), white crystals.
**$^1$H NMR** (400 MHz, $CDCl_3$) δ = 7.37-7.21 (m, 5H), 4.75-4.69 (m, 1H), 4.27-4.19 (m, 2H), 3.90-3.86 (m, 1H), 3.81 (dq, *J* = 2.7, 7.0 Hz, 1H), 3.27 (dd, *J* = 3.4, 13.5 Hz, 1H), 2.81 (dd, *J* = 9.4, 13.4 Hz, 1H), 1.65-1.43 (m, 2H), 1.27 (d, *J* = 7.0 Hz, 3H), 0.99 (t, *J* = 7.5 Hz, 3H) ppm.

(2S, 3R) 3-hydroxy-2-methylpentanoyl-*S*-*N*-acetylcysteamine thioester (2)

Diketide SNAC **2** was synthesized in two steps from compound S4 following protocols of Sharma *et al.* and Peter *et al.*.[12,13]

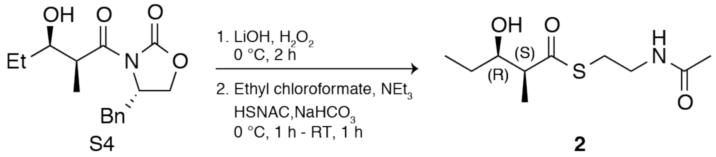

3-hydroxy-2-methyl-pentanoic acid was obtained almost quantitatively following the instructions of Sharma *et al*. and was used without further purification. To the acid (220 mg, 1.66 mmol) in 5 ml THF cooled to 0 °C, 148 µL ethyl chloroformate (168 mg, 1.55 mmol) and 215 µL $NEt_3$ (157 mg, 1.55 mmol) were added at 0 °C. The reaction mixture was stirred excessively for 45 min at 0 °C. Then, 224 µL *N*-acetylcysteamine (251 mg, 2.00 mmol) and 54 mg $NaHCO_3$ dissolved in 2 mL $H_2O$ were added and the reaction mixture was further stirred for 1 h at RT. The aqueous phase was extracted with EtOAc (3 × 20 mL) and the combined organic phases were washed with $H_2O$ (20 mL). The organic phase was dried over anhydrous $MgSO_4$, filtered and the solvent was evaporated *in vacuo*. The crude product was purified by column chromatography (silica gel, DCM:Acetone 10:1 → 8:1 → 6:1 → 4:1) to yield 128 mg (33 %) product **2** as clear liquid.

**Yield**: 33 % (128 mg), clear liquid.
**$^1$H NMR** (400 MHz, $CDCl_3$) δ = 5.76 (s, br, 1H), 3.88-3.83 (m, 1H), 3.53-3.41 (m, 2H), 3.10-2.99 (m, 2H), 2.78-2.72 (m, 1H), 1.98 (s, 3H), 1.57-1.43 (m, 2H), 1.23 (d, *J* = 7.0 Hz, 3H) 0.99 (t, *J* = 7.4 Hz, 3H) ppm.
**MS (ESI+)** found 234.12, for $[M+H]^+$ calculated 234.12; found 256.24, calculated for $[M+Na]^+$ 256.10.

3.4 Fluoromethylmalonyl-CoA (17)
Fluoromethyl-Meldrum's acid (S5)

2,2,5-Trimethyl-1,3-dioxane-4,6-dione (Methyl Meldrum's acid) was either synthesized from methylmalonic acid after the protocol of Bravo-Rodriguez *et al*. or purchased from Merck.[14]
**Yield**: 73 % (1.96 g), white crystals.
**$^1$H NMR** (400 MHz, $CDCl_3$ δ = 3.58 (q, *J* = 7.0 Hz, 1H), 1.81 (s, 3H), 1.77 (s, 3H), 1.58 (d, *J* = 7.1 Hz, 3H) ppm.

Fluoromethyl-Meldrum's acid (S5) was synthesized from compound methyl Meldrum's acid.

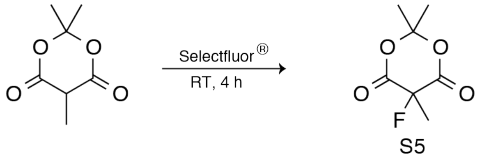

A dry, nitrogen-flushed round bottom flask (500 mL) equipped with a septum was charged with Methyl Meldrum's acid (5 g, 31.6 mmol), the finely powdered Selectfluor (11.8 g, 33.2 mmol;) and 250 mL dry acetonitrile was added. The mixture was stirred for 4 h at RT and solids dissolved after 90 min. The solvent was evaporated and ca. 100 mL DCM and 50 mL water were added. The phases were separated and the aqueous phase was extracted with DCM (3 × 50 mL). Combined organic phases were washed with brine, dried over $MgSO_4$, filtered and the solvent was evaporated *in vacuo* to yield crude Fluoromethyl-Meldrum's acid (5.1 g; 92 %, purity 90 %) as fine white crystals. The crude product still containing educt was adsorbed on silica and

purified by column chromatography (silica gel, hexane:ethylacetate 10:1 → 5:1 → 3:1 → 2:1 → 1:1 → 1:2 → 1:3) to yield 2.8 g (48 %) product S5 as white crystals.
**Yield**: 48 % (2.8 g), white crystals.
**$R_F$-value** (H:EE 1:1): 0.77 (slightly visible with $KMnO_4$-staining solution).
**$^1$H NMR** (400 MHz, $CDCl_3$) δ = 1.93 (d, *J* = 24 Hz, 3H), 1.85 (s, 3H), 1.80 (s, 3H) ppm.
**$^{19}$F NMR** (300 MHz, $CDCl_3$) δ = −153.50 (q, *J* = 24 Hz) ppm.

Fluoromethylmalonic acid thiophenyl halfester (S6)

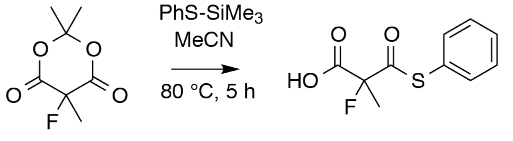

S5 S6

Fluoromethyl-Meldrum's acid (S5) (1.95 g, 11.1 mmol) was covered with ((phenyl)thio)-trimethylsilane (2.4 g, 2.5 mL, 13.2 mmol) and 10 mL dry acetonitrile in an argon-purged round bottom flask. The solution was stirred at reflux for 5 h. The solvent was evaporated and then 10 mL ice-cold saturated $NaHCO_3$ aq. was added on ice with strong gas evaporation. 10 mL water and 80 mL DCM were added, the phases were separated and the aqueous phase was extracted with DCM (3 × 30 mL). The combined organic phases were dried over $MgSO_4$, filtered and the solvent was evaporated *in vacuo*. The crude product was adsorbed on silica and purified by column chromatography (silica gel, dichlormethane:methanol:acetic acid 100:1:0.1) to yield 220 mg (9 %) pure product S6 as colorless oil. The poor yield results from contamination of the product fractions with the respective defluorinated thioester and only the latter fractions containing the pure product were pooled.
**Yield**: 9 % (220 mg), colorless oil
**$R_F$-value** (DCM/MeOH/AcOH: 100/1/0.1): 0.2 ($KMnO_4$-staining).
**$^1$H NMR** (250 MHz, MeOD) δ = 7.48-7.41 (m, 5H), 1.77 (d, *J* = 22 Hz, 3H) ppm.
**$^{19}$F NMR** (300 MHz, $CDCl_3$) δ = −156.4 (q, *J* = 22 Hz, 1H) ppm.

Fluoromethylmalonyl-CoA (17)

Fluoromalonyl-CoA 17 was synthesized via transacylation from the fluoromalonyl thiophenyl halfester S6 to free coenzyme A (CoASH), adapted from Dunn *et al.*[10]

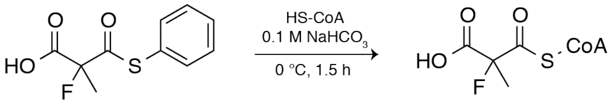

S6 **17**

CoASH (200 mg; 255 µmol) and the fluoromethylmalonyl thiophenyl halfester S6 (200 mg; 876 µmol) were dissolved in 1 mL ice cold $NaHCO_3$ (0.1 M) at 0 °C. The pH was adjusted back to pH 8-9 with 1 mL NaOH (0.2 M) and the reaction was continued for 1.5 h at 0 °C. Then, the reaction mixture (ca. 2 mL) was poured into 20 mL acetone (-20 °C) in a 50 mL falcon and spun for 10 min at 3000 rcf. The supernatant was discarded and the pellet was washed with 1 mL acetone (-20 °C) and dried *in vacuo*. The pellet was dissolved in water and the yield was determined by the absorbance at 260 nm (ε = 16,400 $M^{-1}$ $cm^{-1}$).[11]
**Yield**: 93 %, white crystals.

**$^{1}$H NMR** (300 MHz, $D_2O$) δ = 8.57 (s, 1H), 8.28 (s, 1H), 6.19 (d, *J* = 6.9 Hz , 1H), 4.62-4.56 (m, 1H), 4.28-4.21 (m, 2H), 4.02 (s, 1H), 3.86-3.80 (m, 1H), 3.58-3.52 (m, 1H), 3.48-3.44 (m, 2H), 3.40-3.33 (m, 2H), 3.09-3.03 (m, 2H), 2.48-2.42 (m, 2H), 1.71 (d, *J* = 22 Hz, 3H), 0.88 (s, 3H), 0.73 (s, 3H) ppm.
**$^{19}$F NMR** (300 MHz, $D_2O$) δ = −146.7 (q, *J* = 22 Hz, 1H) ppm.

#### 4. Supplementary Figures:

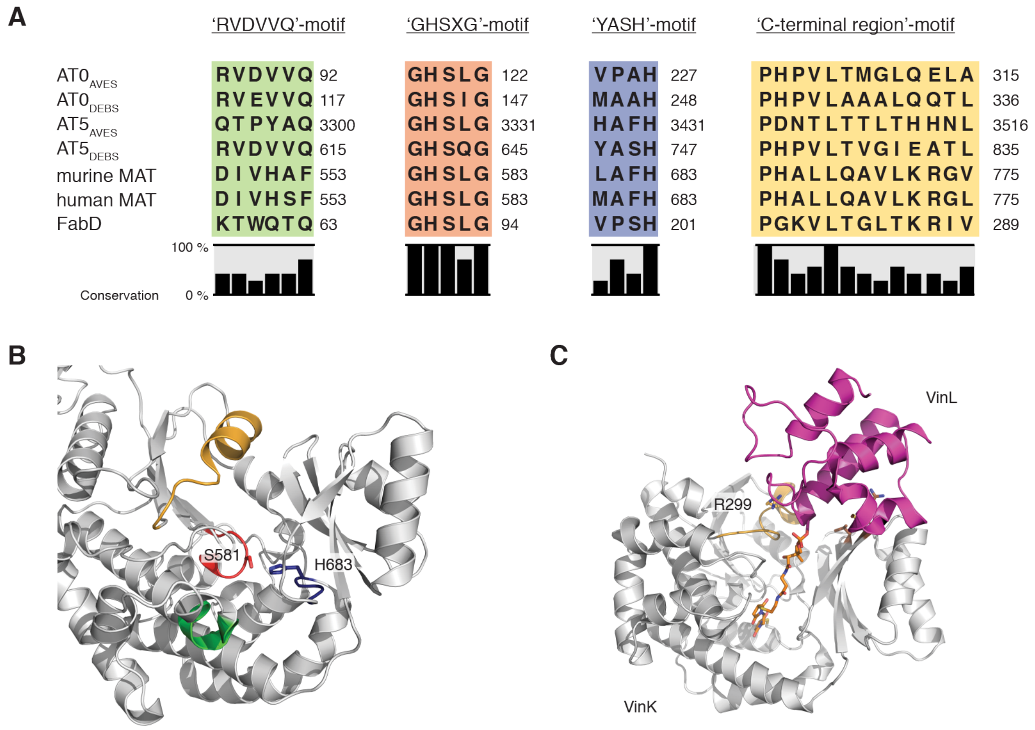

**Fig. S1: Substrate specificity of ATs.**

(A) Sequence alignments of various ATs in regard to the four sequence motifs characterizing substrate specificity. Utilized primary sequences refer to loading-ATs: $AT0_{AVES}$ (Q79ZN1) and $AT0_{DEBS}$ (Q03131), specialized extender-ATs: $AT5_{AVES}$ (Q9S0R7), $AT5_{DEBS}$ (Q03133) and FabD (P0AAI9), and polyspecific ATs: murine MAT (P19096) and human MAT (P49327). The Uniprot code is given in brackets. (B) Cartoon depiction of the human MAT (PDB code: 3hhd) with sequence motifs colored as above. Active site serine (S581) is shown in sticks representation.[15] (C) Crystal structure of the VinK-VinL complex (PDB code: 5czd) in cartoon representation.[16] The ACP (VinL) was chemically cross-linked (sticks, orange) to the AT (VinK). The authors characterized the binding interface and identified crucial residues including R299 in the 'C-terminal' region (brightorange).

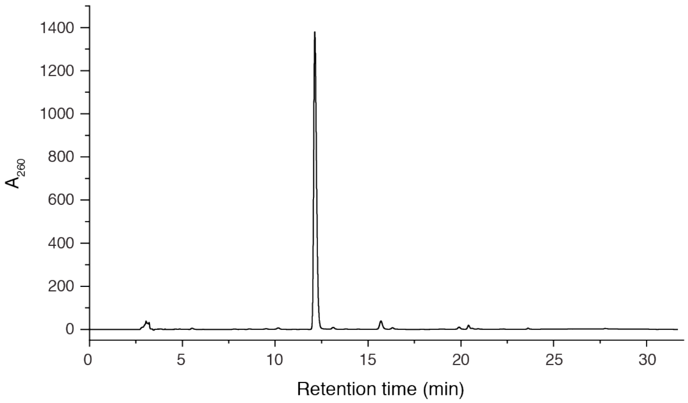

**Fig. S2: Quality control of precipitated F-Mal-CoA.**

F-Mal-CoA was purified with HPLC-UV using the gradient described in method section, showing almost quantitative conversion of CoA (15.5 min). F-Mal-CoA (12.1 min) has a comparable elution time to Mal-CoA (12.0 min).

A

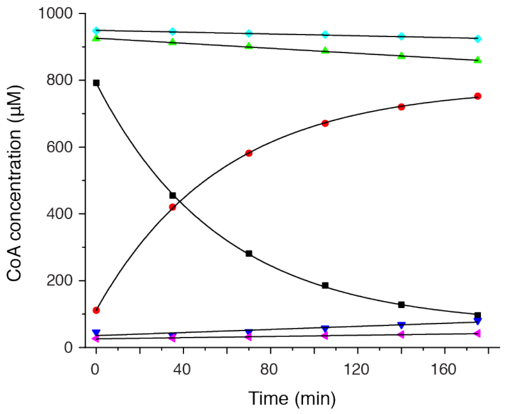

B

| Hydrolyse | F-Mal-CoA | Temperature |
|---|---|---|
| with DTT ($\mu$M min$^{-1}$) | 12 ± 5 | 22 °C |
| with DTT ($\mu$M min$^{-1}$) | 0.11 ± 0.01 | 5 °C |
| w/o DTT ($\mu$M min$^{-1}$) | 0.31 ± 0.01 | 22 °C |

C

| Hydrolyse | Mal-CoA | MM-CoA | F-Mal-CoA |
|---|---|---|---|
| MAT-mediated (s$^{-1}$) | 1.13E-03 | 7.20E-04 | 1.81E-02 |
| Total ($\mu$M min$^{-1}$) | 0.03 | 0.04 | 0.35 |

**Fig. S3: Further information on F-Mal-CoA.**

(A) F-Mal-stability was monitored with HPLC-UV using the gradient described in the method section. Peak areas were converted to concentration using a CoA calibration curve. F-Mal-CoA degradation (black) and its corresponding CoA liberation (red) in buffer with DTT at 22 °C was fitted with an exponential decay function in OriginPro 8.5. Degradation without DTT at 5 °C (cyan: F-Mal-degradation, purple: CoA-liberation) and at 22 °C (green: F-Mal-degradation, blue: CoA-liberation) was fitted by linear regression (buffer: (400 mM phosphate buffer, 20 % (v/v) glycerol, 2 mM or 0 mM DTT, 1 mM EDTA, 0.8 % DMSO (pH 7.2)). (B) Initial slope average of F-Mal-degradation and CoA-liberation of the corresponding condition (C) Hydrolysis rates of the extender substrates determined by the αKGDH assay. Final substrate concentrations in the assays were 20 µM X-CoA and 0.2 µM KS°-MAT. Fast F-Mal-CoA degradation by DTT was the reason why DTT was omitted in assays when using this substrate.

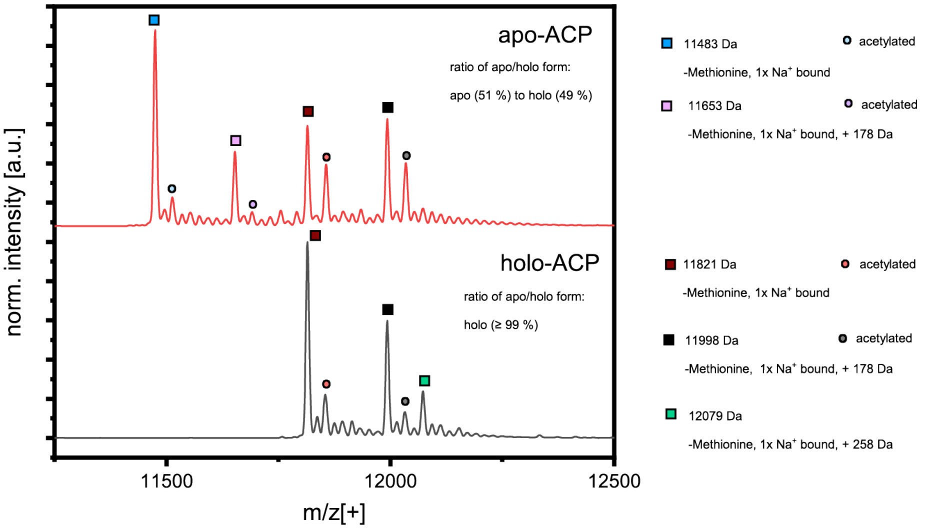

**Fig. S4: Quality control of DEBS ACP6.**

Mass spectroscopic analysis of DEBS ACP6, co-expressed with Sfp (red) or Npt (black), was performed with ESI in positive mode (SYNAPT G-2, waters). Co-expression with Sfp led to a mixture of 49 % of phosphopantetheinlated (holo-form, brown square, theoretical mass: 11816.10 Da) and 51 % apo-form (blue square, theoretical mass = 11475.76 Da), while co-expression with Npt resulted in quantitative conversion to the holo-form. A mass shift of 178 Da (purple square for the apo-form and black square for the holo-form) and 258 Da (cyan square for the holo-form) could be explained by His-tag phosphogluconoylation in *E. coli*, whereas a mass shift of 42 Da corresponds to the acetylated form (circles with the corresponding color).

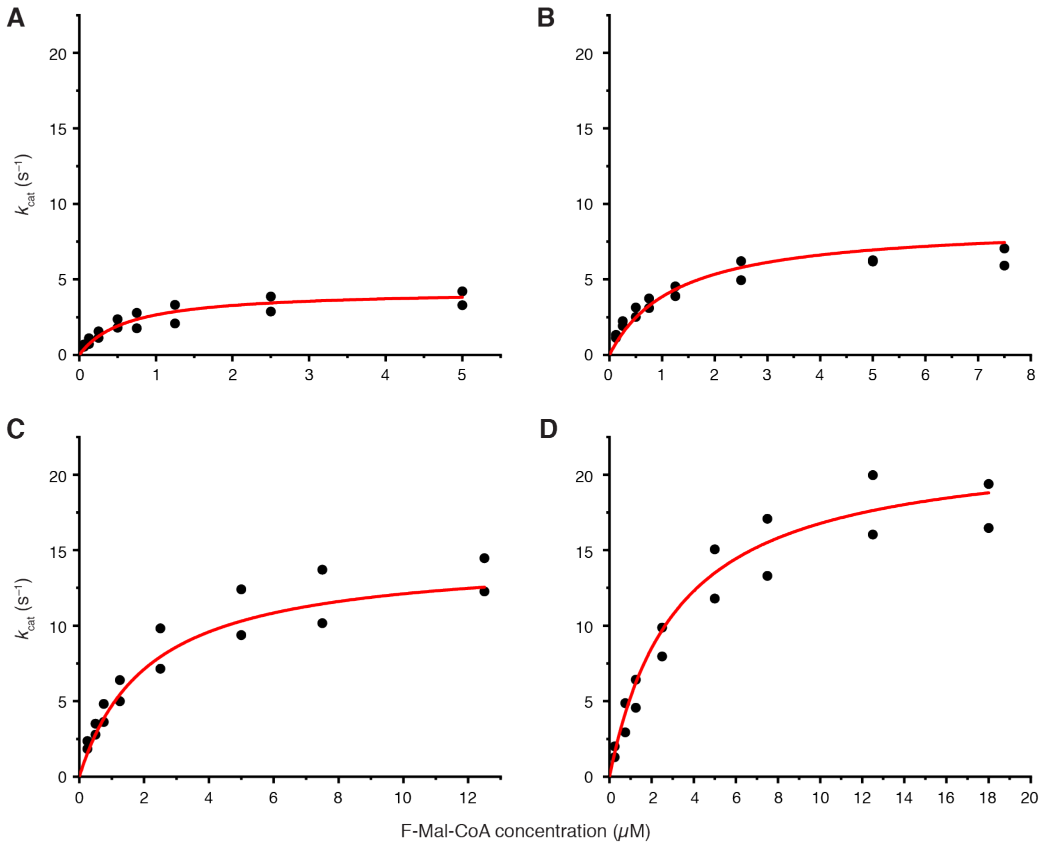

**Fig. S5: Global Michaelis-Menten fit of FAS MAT-mediated transacylation of fluoromalonyl moieties to FAS ACP.**

Initial velocities were plotted against fluoromalonyl-CoA (F-Mal-CoA) concentrations at four fixed ACP concentrations: (A) 11 μM, (B) 24 μM, (C) 49 μM and 100 μM ACP. Data were fit globally with the Michaelis-Menten equation assuming a ping-pong bi-bi mechanism (n = 2).

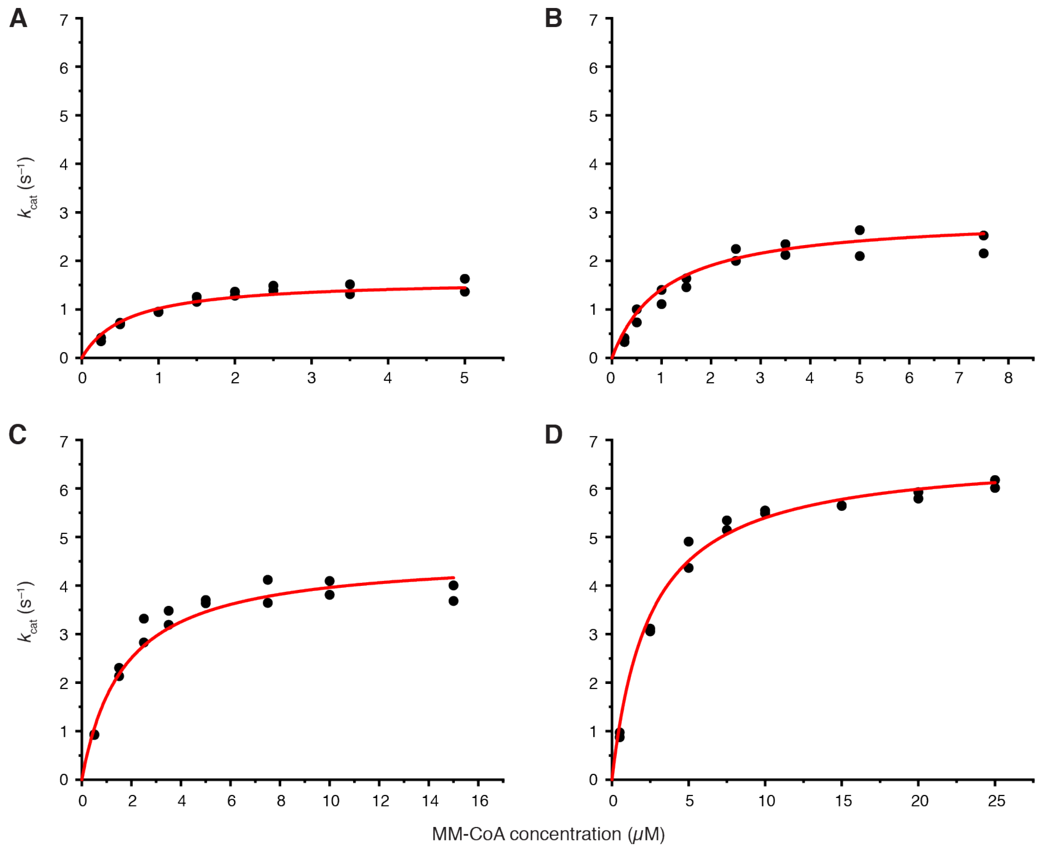

**Fig. S6: Global Michaelis-Menten fit of FAS MAT-mediated transacylation of methylmalonyl moieties to DEBS ACP6.**

Initial velocities were plotted against methylmalonyl-CoA (MM-CoA) concentrations at four fixed ACP concentrations: (A) 53 µM, (B) 107 µM, (C) 200 µM and 373 µM ACP. Data were fit globally with the Michaelis-Menten equation assuming a ping-pong bi-bi mechanism (n = 2).

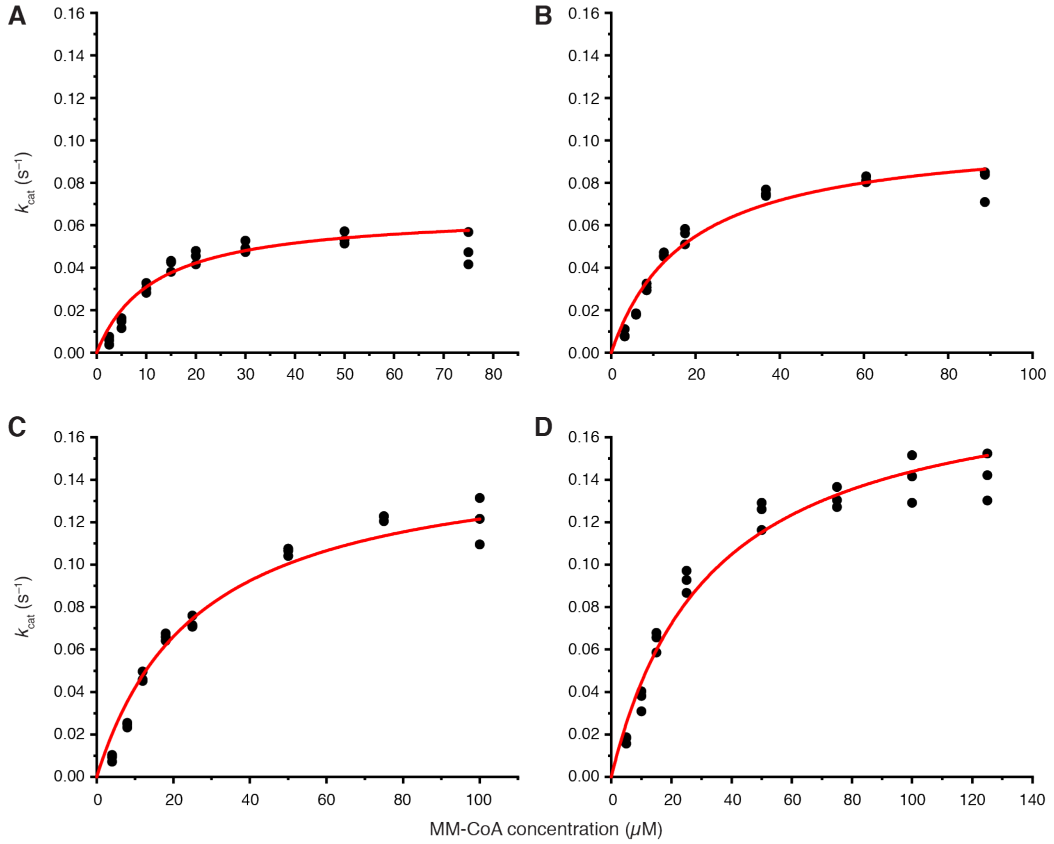

**Fig. S7: Global Michaelis-Menten fit of DEBS AT6-mediated transacylation of methylmalonyl moieties to DEBS ACP6.**

Initial velocities were plotted against methylmalonyl-CoA (MM-CoA) concentrations at four fixed ACP concentrations: (A) 64 µM, (B) 119 µM, (C) 238 µM and 406 µM ACP. Data were fit globally with the Michaelis-Menten equation assuming a ping-pong bi-bi mechanism (n = 3).

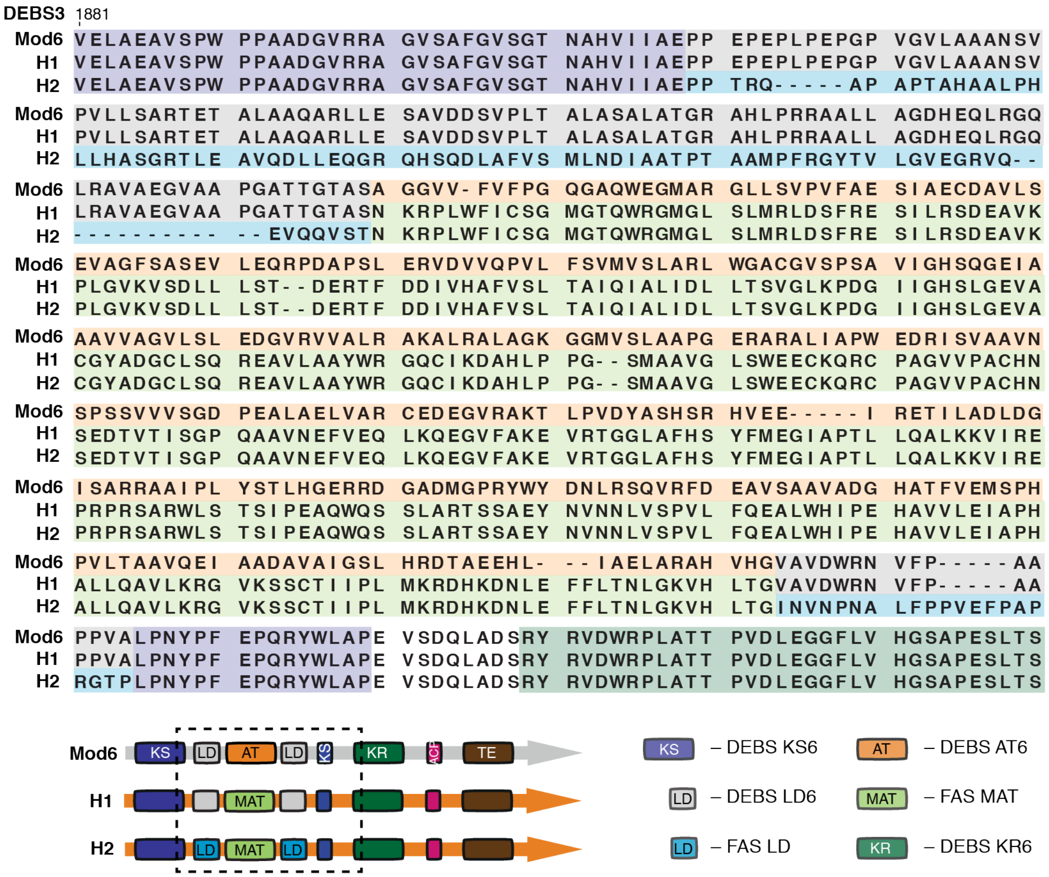

**Fig. S8: Design of DEBS/FAS hybrids.**

Sequence alignment of wildtype DEBS M6+TE (**WT**) with both hybrid DEBS/FAS constructs **H1** and **H2**. In construct **H1**, the AT6 domain of DEBS was exchanged by the MAT domain of murine FAS. For construct **H2**, the AT6 domain of DEBS plus the adjacent linker domain was exchanged with LD-MAT from murine FAS. The sequence alignment is colored according to the color code of the attached domain architecture.

An AT-exchange was published by Yuzawa *et al.* before and our hybrid **H1** is similar to their **D2**, although we defined the terminal long α-helix as part of MAT and not as part of the post-AT linker. Similarly, our hybrid **H2** equals the design of **D1**.[5]

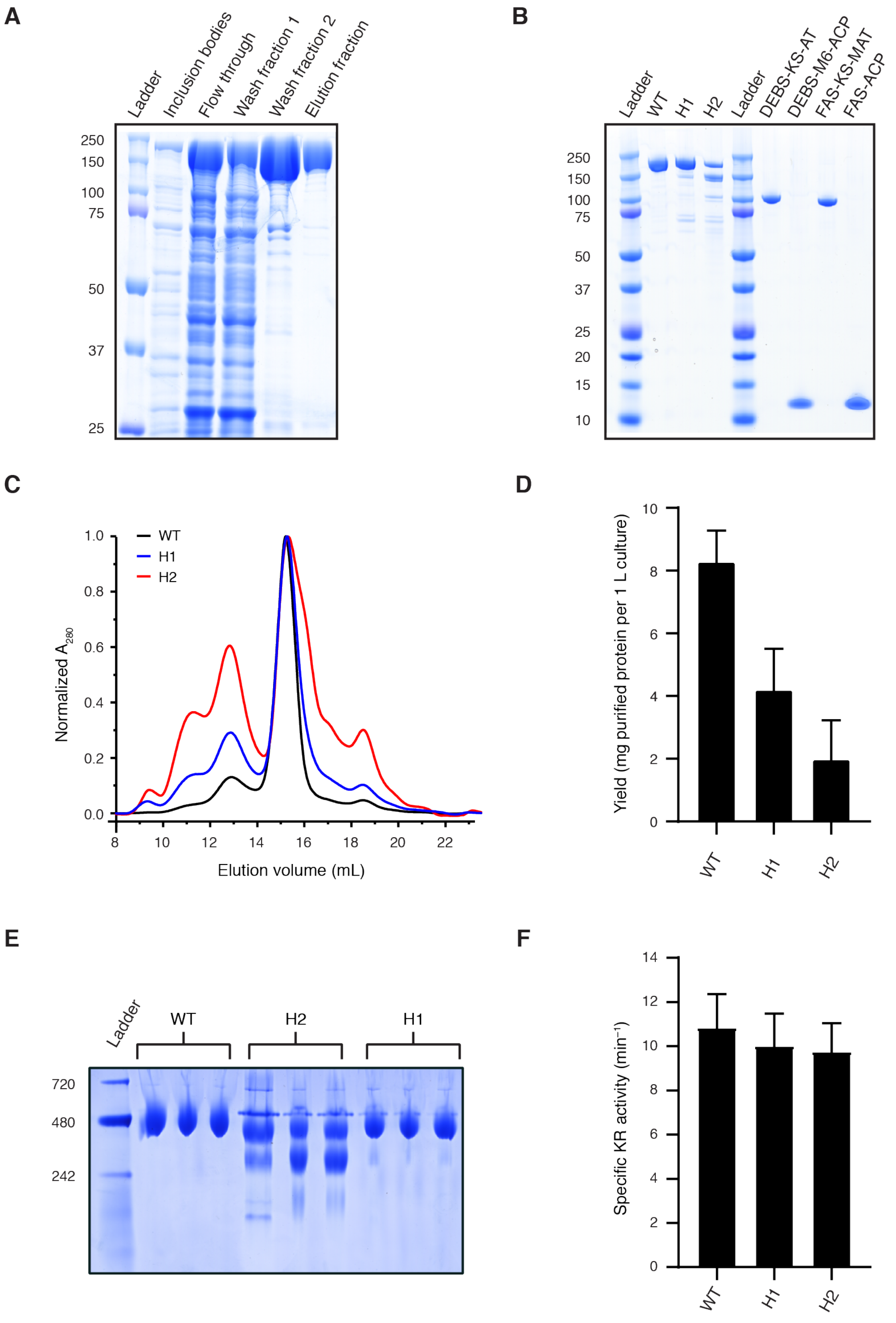

**Fig. S9: Purification and quality control of DEBS M6+TE and DEBS/FAS hybrids H1 and H2.**

(A) Purification of DEBS M6+TE (WT) co-expressed with Sfp (SDS-PAGE (7.5 % Tris-glycine, 1 % SDS buffer). After centrifugation, supernatant was purified with Ni-chelating chromatography and the pellet was analyzed for inclusion bodies. DEBS M6+TE has little tendency to aggregate and was received as a pure protein. (B) Purity of used enzymes after SEC (SDS-PAGE (NuPAGE 4-12 % Bis-Tris, Thermo Fisher)). (C) Oligomeric state of purified WT, **H1** and **H2** analyzed by SEC with absorbance normalized to the highest peak. While the chromatogram of H1 is similar to the WT, the main peak shoulder of H2 indicates the presence of monomers. (D) Comparison of yields after purification (IMAC, SEC) of three different expression cultures of DEBS-M6+TE and variants (n = 3). (E) Native gel electrophoresis (3-12 % Bis-Tris, Thermo Fisher) of DEBS-M6+TE and variants showing dimer for **WT**, **H1**, and **H2** as a mixture of dimer and monomer. Each enzyme is shown in biological triplicates (n = 3) (F) KR specific assay was performed with *trans*-1-decalone indicating 92 % KR-activity for **H1** and 90 % KR-activity for **H2** in biological and technical triplicates (n = 3) compared to the **WT**. Final substrate concentrations in the assay were 0.3 µM enzyme, 2 mM *trans*-1-decalone and 60 µM NADPH.

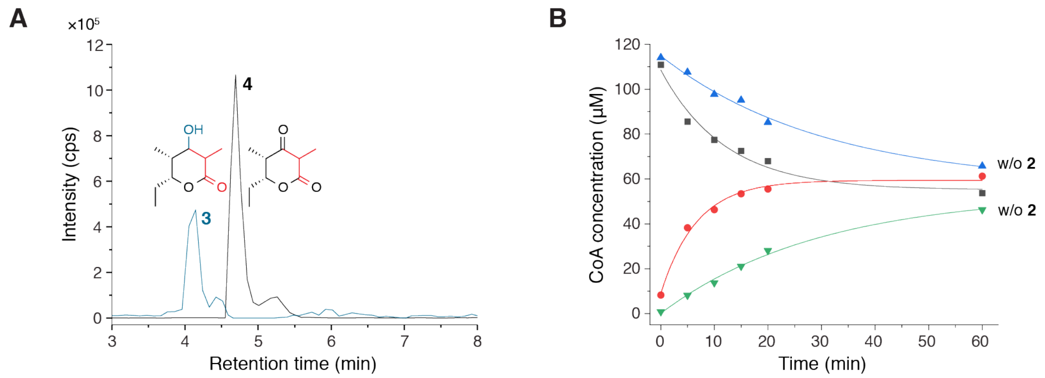

**Fig. S10: Functionality of DEBS M6+TE.**

(A) Reductive WT-mediated chain extension of diketide-SNAC 2 with MM-CoA monitored by HPLC-MS [EIC: 3 $[M+Na]^+$ $m/z$ = 194.98 and 4 $[M–H]^–$ $m/z$ = 169.12]. (B) Non-reductive enzyme-mediated chain extension of diketide-SNAC 2 with MM-CoA monitored by HPLC-UV exemplified by **WT**. MMal-CoA consumption and CoA liberation was tracked at $A_{260}$ at defined time points. Enzyme-mediated hydrolysis was measured without 2. CoA concentrations of each time point were determined by using the internal standard HyBu-CoA and fitted with an exponential decay function in OriginPro 8.5. The sum of MMal-CoA and CoA concentrations of each sample was used as quality control and outliers were excluded.

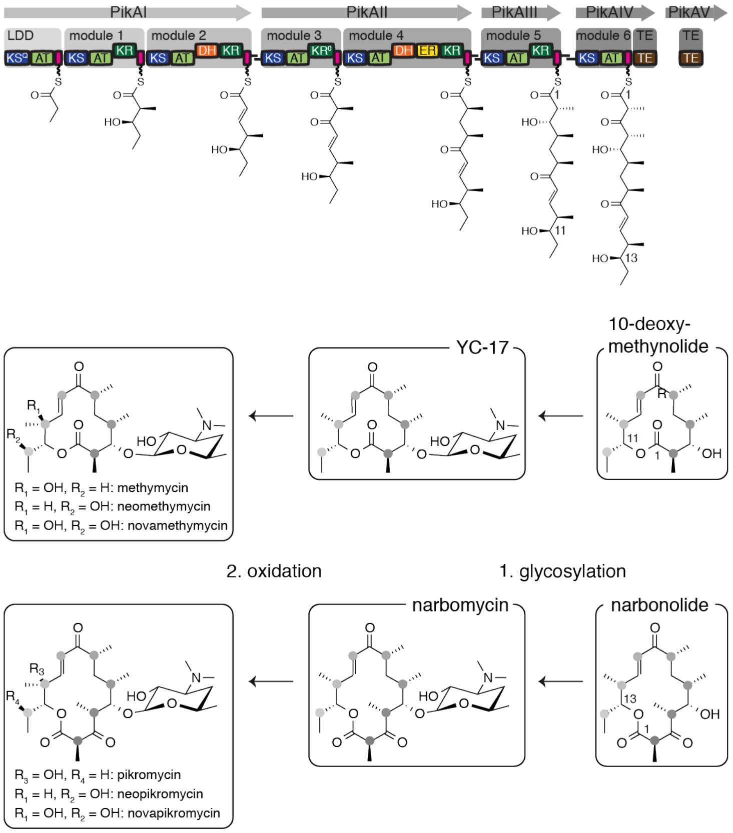

**Fig. S11: Biosynthesis of the pikromycin/methymycin precursor molecules.**

The pikromycin synthase catalyze the assembly like biosynthesis of 10-deoxymethynolide and narbonolide, precursor molecules for YC-17 and narbomycin, which can be oxidized to various compounds. The enzymes are distributed on five peptide chains including a N-terminal loading module and two C-terminal thioesterases.

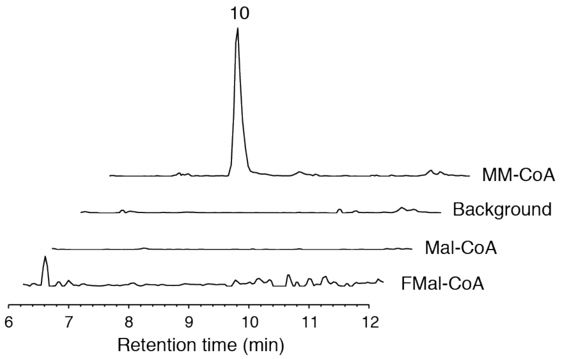

**Fig. S12: Extender substrate specificity of DEBS M6+TE (WT).**

Reductive **WT**-mediated chain extension of pentaketide 9 with MM-CoA, Mal-CoA and F-Mal-CoA monitored by HPLC-MS. The background was performed without elongation substrate and only the EIC of the negative control is shown for the mass of compound 10 (elongation product with MM-CoA). EIC of compound 12 (elongation product of Mal-CoA) and 14 (elongation product of F-Mal-CoA) are shown although the masses were not found. Data are normalized with respect to the highest peak of H1. [EIC: 10 $[M+Na]^+$ *m*/*z* = 319.11; 12 $[M+Na]^+$ *m*/*z* = 305.09 and 14 $[M+Na]^+$ *m*/*z* = 323.08).

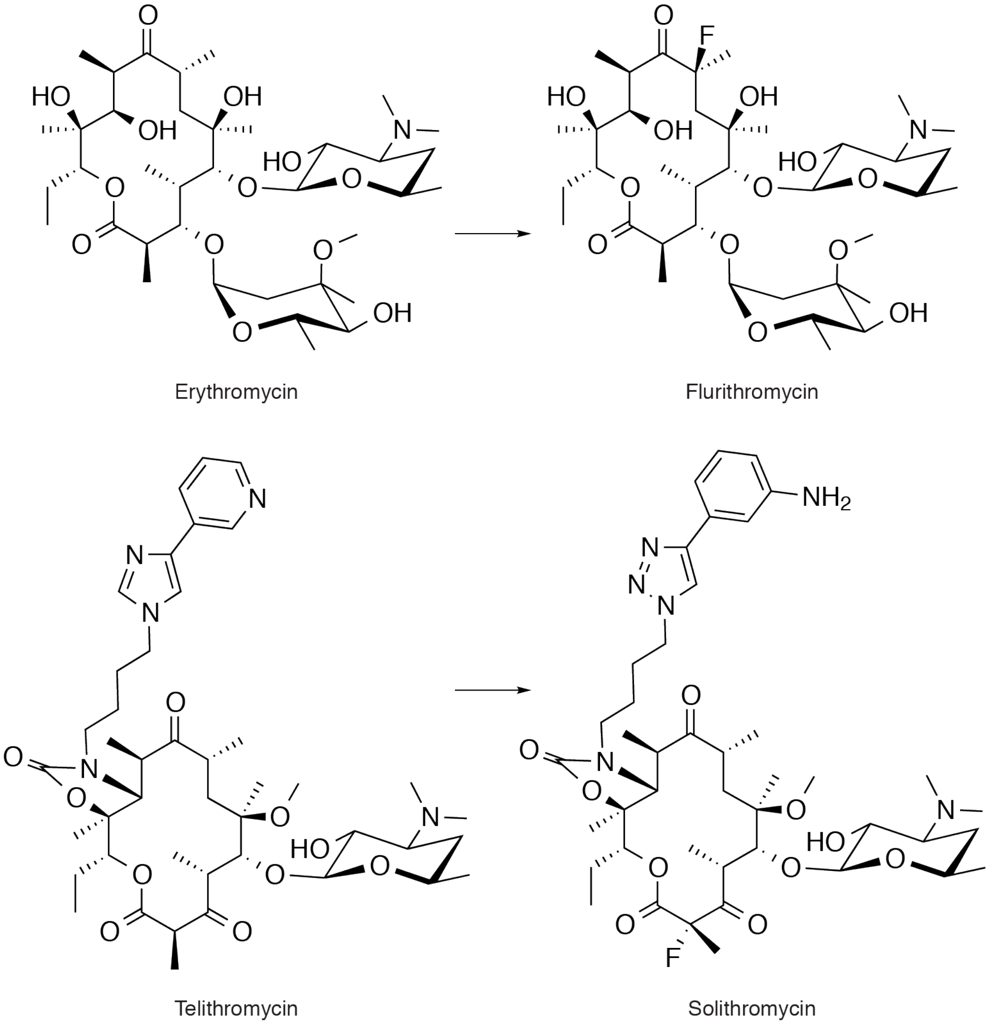

**Fig. S13: Chemical structures of erythromycin and derivatives.**

The structures of the natural product erythromycin and the semi-synthetic second generation fluoro-derivative flurithromycin are shown. Telithromycin is also a semi-synthetic erythromycin derivative, which is a FDA-approved third generation antibiotic. Due to side-effects of telithromycin the fourth generation antibiotic solithromycin was developed, which is under review for approval.

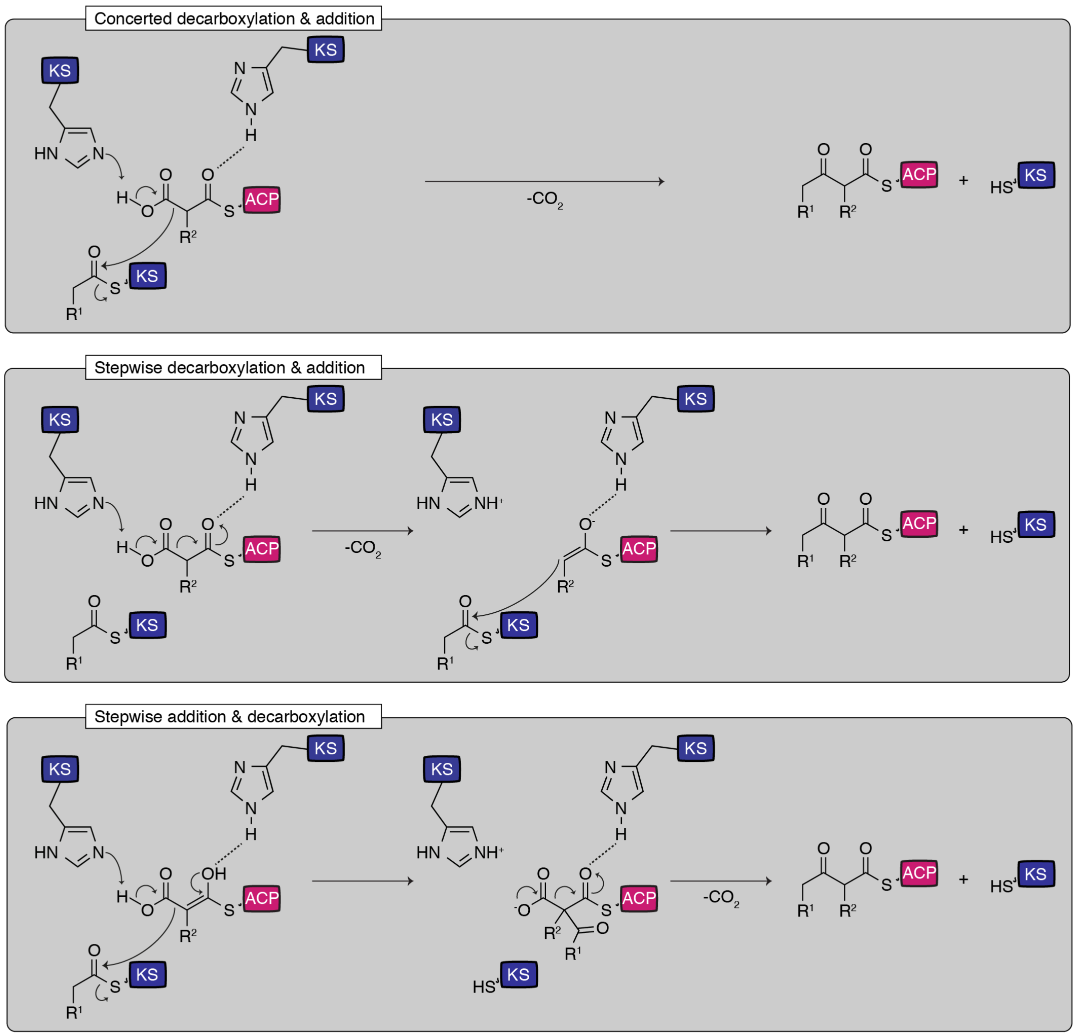

**Fig. S14: Possible mechanisms for the KS-catalyzed Claisen condensation.**

After acylation of the KS with the starter substrate or intermediate and acylation of the ACP with the elongation substrate, three distinct mechanisms for the KS-catalyzed chain elongation are possible. In the first mechanism, the decarboxylation and C-C bond formation occur simultaneously. In contrast, two mechanisms are possible where either the decarboxylation or the C-C bond formation precede the other. Adapted from Blaquiere *et al.*[17].

We note that the biosynthesis of compound **18** is indicative of a Claisen condensation reaction that proceeds via decarboxylation to the enolate and nucleophilic C-C bond formation either occurring in concerted manner or in two steps by C-C bond formation succeeding decarboxylation. Demonstrating the direct condensation of disubstituted malonyl-CoA substrates by a conventional KS-domain expands the potential of PKS engineering strategies and enlarges the spectrum of accessible compounds.

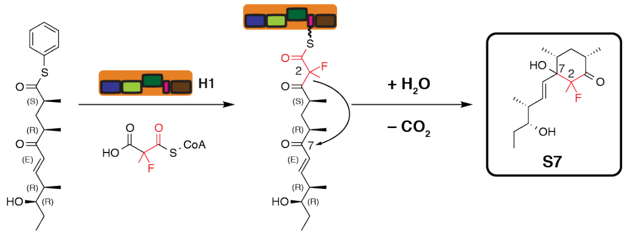

**Fig. S15: Postulated mechanism for the production of S7.**

We note that elongating the pentaketide with F-MM-CoA in the absence of NADPH led to product **S7**, which is the respective derivative to the previously identified compound **16**. This data indicates that cyclohexanone ring formation of **S7** occurs after TE-catalyzed hydrolysis with subsequent spontaneous decarboxylation and ring closure, similar to the proposed formation of pacificanone, a side product of the rosamicin PKS.[18]

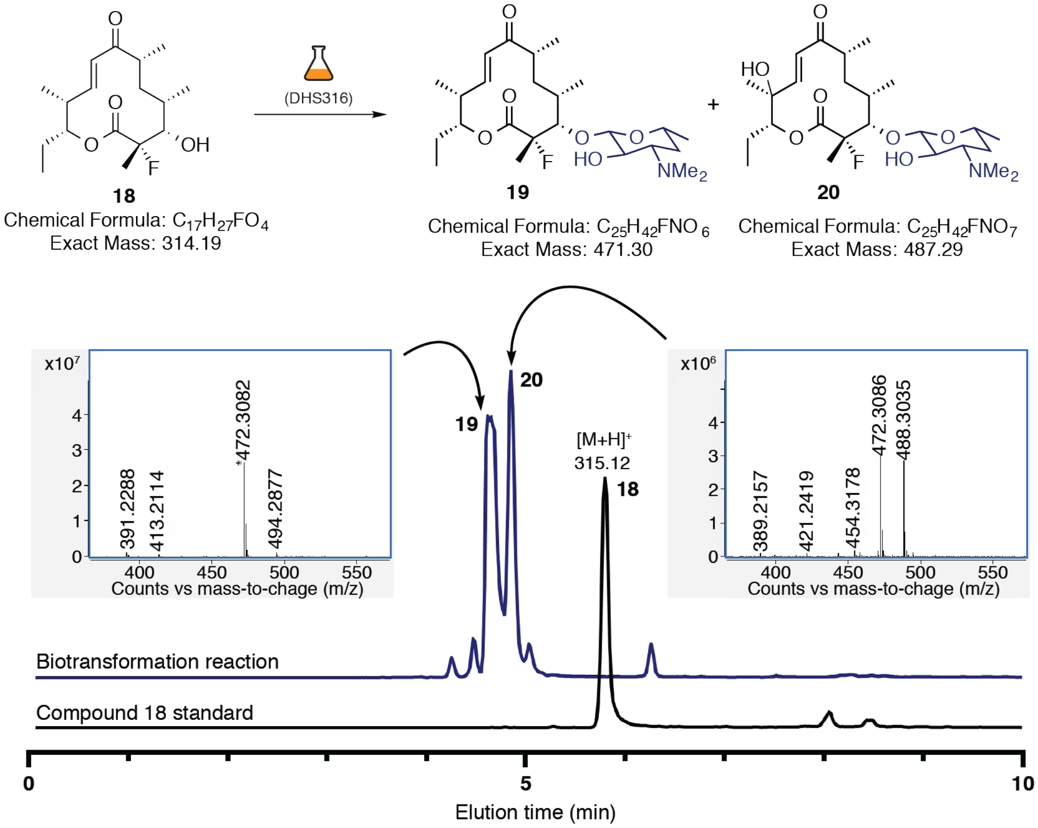

**Fig. S16: Biotransformation of compound 18.**

Overlaid LC-MS traces of compound **18** and compounds after biotransformation. Mass spectra were taken in positive mode. We note that glycosylated and oxidized species with m/z of 487 is likely not C2-fluoromethylated methymycin, because the *Streptomyces venezuelae* strain used for biotransformation is PikC deficient (the cytochrome P450 monooxygenase required for methymycin production).

#### 5. Supplementary Tables:

**Table S1: Absolute kinetic parameters for MAT- and DEBS AT6-mediated transfer.**

| AT/MAT | Substrate | ACP | $k_{cat}$ ($s^{-1}$) | $K_S$ (μM) | $K_{ACP}$ (μM) | $\frac{k_{cat}}{K_S}$ ($M^{-1}$ $s^{-1}$) |
|---|---|---|---|---|---|---|
| FAS | F-Mal-CoA | FAS | 43.28 ± 3.04 | 6.3 ± 0.6 | 95 ± 11 | $6.9 \times 10^6$ |
| FAS | MM-CoA | DEBS M6 | 14.08 ± 0.89 | 5.2 ± 0.4 | 408 ± 43 | $2.7 \times 10^6$ |
| DEBS M6 | MM-CoA | DEBS M6 | 0.29 ± 0.02 | 50.8 ± 5.9 | 217 ± 32 | $5.8 \times 10^3$ |

#### Table S2: Compounds table.

| compound number | chemical formula | turnover rate ($min^{-1}$) | calculated mass | observed mass | observed mass (HR) | retention time [min] |
|---|---|---|---|---|---|---|
| 3 | C9H16O3 | WT = 0.29 ± 0.03<br>H1 = 0.19 ± 0.06<br>H2 = 0.07 ± 0.02 | $[M+H]^+$ = 173,1178<br>$[M+Na]^+$ = 195,0997<br>$[M-H_2O+H]^+$ = 155,1072<br>$[M-H]^-$ = 171,1021 | $[M+H]^+$ = 173,03<br>$[M+Na]^+$ = 194,98<br>$[M-H_2O+H]^+$ = 155,01 | nd | 4,1 |
| 4 | C9H14O3 | WT = 1.4 ± 0.4<br>H1 = 0.6 ± 0.1<br>H2 = 0.23 ± 0.02 | $[M+H]^+$ = 171,1021<br>$[M+Na]^+$ = 193,0841<br>$[M-H_2O+H]^+$ = 153,0916<br>$[M-H]^-$ = 169,0865 | $[M+H]^+$ = 171,00<br>$[M-H_2O+H]^+$ = 152,96<br>$[M-H]^-$ = 169.12 | nd | 4,8 |
| 5 | C8H14O3 | WT = 0.012 ± 0.003<br>H1 = 0.19 ± 0.02 | $[M+H]^+$ = 159,1021<br>$[M+Na]^+$ = 181,0841<br>$[M-H_2O+H]^+$ = 141,0916<br>$[M-H]^-$ = 157,0865 | $[M+Na]^+$ = 181,03 | nd | 3,3 |
| 6 | C8H12O3 | WT = 0.0 ± 0.1<br>H1 = 1.2 ± 0.3 | $[M+H]^+$ = 157,0865<br>$[M+Na]^+$ = 179,0684<br>$[M-H_2O+H]^+$ = 139,0759<br>$[M-H]^-$ = 155,0708 | $[M-H]^-$ = 155,16 | nd | 4,6 |
| 7 | C8H13FO3 | nd | $[M+H]^+$ = 177,0927<br>$[M+Na]^+$ = 199,0747<br>$[M-H_2O+H]^+$ = 159,0822<br>$[M-H]^-$ = 175,0771 | - | - | - |
| 8 | C8H11FO3 | nd | $[M+H]^+$ = 175,0771<br>$[M+Na]^+$ = 197,0590<br>$[M-H_2O+H]^+$ = 157,0665<br>$[M-H]^-$ = 173,0614 | $[M-H]^-$ = 173,11 | nd | 4,8 |
| 10 | C17H28O4 | WT = 0.30 ± 0.05<br>H1 = 0.12 ± 0.01<br>H1 = 0.158 ± 0.001* | $[M+H]^+$ = 297,2066<br>$[M+Na]^+$ = 319,1886<br>$[M-H_2O+H]^+$ = 279,1960<br>$[M-H]^-$ = 295,1910 | $[M+H]^+$ = 297,11<br>$[M+Na]^+$ = 319,11<br>$[M-H_2O+H]^+$ = 279,11 | $[M+H]^+$ = 297,2056<br>$[M+Na]^+$ = 319,1876<br>$[M-H_2O+H]^+$ = 279,1950 | 8,1 |
| 11 | C17H26O4 | nd | $[M+H]^+$ = 295,1910<br>$[M+Na]^+$ = 317,1729<br>$[M-H_2O+H]^+$ = 277,1804<br>$[M-H]^-$ = 293,1753 | $[M+H]^+$ = 295,09<br>$[M+Na]^+$ = 317,09<br>$[M-H_2O+H]^+$ = 277,08 | $[M+H]^+$ = 295.1904<br>$[M+Na]^+$ = 317,1720<br>$[M-H_2O+H]^+$ = 277,1790 | 9,1 |
| 12 | C16H26O4 | WT = -0.009 ± 0.004<br>H1 = 0.22 ± 0.03 | $[M+H]^+$ = 283,1910<br>$[M+Na]^+$ = 305,1729<br>$[M-H_2O+H]^+$ = 265,1804<br>$[M-H]^-$ = 281,1753 | $[M+Na]^+$ = 305,09<br>$[M-H_2O+H]^+$ = 256,09 | [M+H]+ = 283,1899<br>$[M+Na]^+$ = 305,1719<br>[M-H2O+H]+ = 265,1796 | 7,5 |
| 13 | C16H24O4 | nd | $[M+H]^+$ = 281,1753<br>$[M+Na]^+$ = 303,1573<br>$[M-H_2O+H]^+$ = 263,1647<br>$[M-H]^-$ = 279,1597 | $[M+H]^+$ = 281,07<br>$[M+Na]^+$ = 303,08<br>$[M-H_2O+H]^+$ = 263,08 | $[M+H]^+$ = 281,1741<br>$[M+Na]^+$ = 303,1559<br>$[M-H_2O+H]^+$ = 263,1637 | 7.2 / 9.5 |
| 14 | C16H25FO4 | WT = 0.003 ± 0.005<br>H1 = 0.056 ± 0.004<br>H1 = 0.04 ± 0.01* | $[M+H]^+$ = 301,1815<br>$[M+Na]^+$ = 323,1635<br>$[M-H_2O+H]^+$ = 283,1710<br>$[M-H]^-$ = 299,1659 | $[M+H]^+$ = 301,10<br>$[M+Na]^+$ = 323,08<br>$[M-H_2O+H]^+$ = 283,08 | $[M+H]^+$ = 301,1807<br>$[M+Na]^+$ = 323,1625<br>$[M-H_2O+H]^+$ = 283,1700 | 7.7 / 8.0 |
| 15 | C16H23FO4 | nd | $[M+H]^+$ = 299,1659<br>$[M+Na]^+$ = 321,1478<br>$[M-H_2O+H]^+$ = 281,1553<br>$[M-H]^-$ = 297,1502 | $[M+H]^+$ = 299,08<br>$[M+Na]^+$ = 321,07<br>$[M-H_2O+H]^+$ = 281,06 | $[M+H]^+$ = 299,1642<br>$[M+Na]^+$ = 321,1457<br>$[M-H_2O+H]^+$ = 281,1537 | 8.4 / 8.9 |
| 16 | C15H25FO3 | nd | $[M+H]^+$ = 273.1866<br>$[M+Na]^+$ = 295.1686<br>$[M-H_2O+H]^+$ =255.1761<br>$[M_2+Na]^+$ = 567.3474 | nd | $[M+H]^+$ =273.1859<br>$[M+Na]^+$ = 295.1677<br>$[M-H_2O+H]^+$ =255.1754<br>$[M_2+Na]^+$ = 567.3462 | 7.0 |
| 18 | C19H31FO5 | 0.14 ± 0.02* | $[M+H]^+$ = 315,1972<br>$[M+Na]^+$ = 337,1791<br>$[M-H_2O+H]^+$ = 297,1866<br>$[M-H]^-$ = 313,1815 | nd | [M+H]+ = 315.1961<br>$[M+Na]^+$ = 337.1781<br>$[M-H_2O+H]^+$ = 297.1857 | 7.8 |
| 19 | C25H42FNO6 | nd | [M+H]+ = 472.3075<br>[M+Na]+ = 494.2894 | nd | [M+H]+ = 472.3086<br>[M+Na]+ = 494.2877 | 4.6 |
| 20 | C25H42FNO7 | nd | [M+H]+ = 488.3024 | nd | [M+H]+ = 488.3035 | 4.8 |
| S7 | C16H27FO3 | nd | [M+H]+ = 287,2023<br>[M+Na]+ = 309,1842<br>[M-H2O+H]+ = 269,1917<br>$[M_2+Na]$+ = 595,3787 | nd | [M+H]+ = 287.2017<br>[M+Na]+ = 309.1836<br>[M-H2O+H]+ =269.1913<br>$[M_2+Na]$+ = 595.3779 | 7.0 |

Turnover rates highlighted with an asterisk were performed with slightly different conditions (see chapter 1.2.10).

**Table S3: Used plasmids**

| Construct | Name | Ref. |
|---|---|---|
| $KS^{C161G}$-MAT (WT) | pAR70_StrepI_m(KS(C161G)_MAT)_H8_pET22b | 1 |
| FAS ACP | pAR352_StrepII_mACP_H8_RBS_SFP_pET22b | 1 |
| DEBS ACP6 | pMJD094_DEBS-M6-H8-ACP_W pET28a | This study |
| Sfp | pAR357_SFP_pCDF-1b | 1 |
| Npt | pMJD091_Sppt_pCDF | This study |
| DEBS $KS6^{C1661G}$-AT6 | pAR432_DD2_KS6_C1661G_AT6_H6_pET | This study |
| DEBS M6+TE | pBL18_DEBS_M6_TE | Khosla Lab |
| FAS | pAR264_StrepI_Not1_mFASm_H8_pET22b | 1 |
| **H2** | pMJD076_DEBS_M6_TE_mfASm_LDAT | This study |
| **H1** | pMJD077_DEBS_M6_TE_mfASm_AT | This study |

**Table S4: Primers**

| Nr. | Name | Sequence (5'3') | Template |
|---|---|---|---|
| AR26 | AMP_infusion_for | GAG GAC CGA AGG AGC TAA CC | pBL18 |
| AR27 | AMP_infusion_rev | GGT TAG CTC CTT CGG TCC TC | pBL18 |
| AR719 | LE_H4_for | CTCGAGCACCACCACCAC | pBL18 |
| AR720 | DEBS_AT6_H4_rev | GTGGTGGTGCTCGAGGCTGT CGGCGAGCTG | pBL18 |
| AR721 | DEBS_AT6_C1661G_for | CACGGTGGACACGGCGGGC TCGTCGTCGTTGGTGG | pBL18 |
| AR722 | DEBS_AT6_C1661G_rev | CGCCGTGTCCACCG | pBL18 |
| MJD087 | mFASm_LDAT_fwd | acacggcaggccc | pAR264 |
| MJD088 | mFASm_LDAT_rev | aggagtccctcgggg | pAR264 |
| MJD101 | DEBS_LDAT_fwd | ccccgagggactcctCTGCCCAACT ACCCGTTCGAG | pBL18 |
| MJD102 | DEBS_LDAT_rev | caggggcctgccgtgtCGGGGGCTC GGCGAT | pBL18 |
| MJD091 | mFASm_AT_fwd | aacaagcgcccactctg | pAR264 |
| MJD092 | mFASm_AT_rev | tgtgaggtgcaccttgc | pAR264 |
| MJD105 | DEBS_AT_fwd | caaggtgcacctcacaGGCGTGGCC GTGGAC | pBL18 |
| MJD106 | DEBS_AT_rev | gagtgggcgcttgttGGAGGCGGTTC CGGTG | pBL18 |
| MJD136 | pCDFBB_fwd | TAATTAACCTAGGCTGCTGC CAC | pAR357 |
| MJD137 | pCDFBB_rev | GGTATATCTCCTTATTAAAG TTAAACAAAATTATTTC | pAR357 |
| MJD138 | Sppt_fwd | ATAAGGAGATATACCATGAT TGAGAAGTTACTCC | pET21a_Sp pt |
| MJD139 | Sppt_rev | AGCCTAGGTTAAttaCTCGAG TGCGGCCG | pET21a_Sp pt |
| MJD145 | ACP6_W_for | CCGCGCGGCAGCCATATGTG GGCGGCCCCG | pBL18 |
| MJD146 | ACP_W_rev | ACGGAGCTCGAATTCAGAGC TGCTGTCCTATGTGGTCG | pBL18 |

#### 6. Supplementary note: Stereochemistry of 18

During the biosynthesis of compound **18**, two new stereo centers at position C2 and C3 were formed. $^{1}$H-NMR and $^{19}$F-NMR showed that only one stereoisomer was synthesized. To determine the stereochemistry of compound **18**, we predicted the 3D structures of all possible diastereomers using the software Avogadro with UFF force field (Steepest decent) algorithm and no constraints (see supplementary note figure 1). The template of the models was the structure of 10-deoxymethynolide, solved by crystallization of the pikromycin thioesterase (PDB = 2hfk, E4H). We used the models to calculated characteristic distances (d1-d5) in the stereoisomers, more precisely the distance of the hydrogen of the alkene group at C8 to the nearest neighbors (methyl groups at C6 and C9, methylene group at C5) and to the methyl group at C2, as well as the distance of the methyl group at C2 to the hydrogen of the methylene group at C5 (see supplementary note figure 1-2). Furthermore, we calculated the dihedral angel between the hydrogen of the CHOH group at C3 to the hydrogen of the $CHCH_3$ group at C4 (HCCH angle, blue arrow) and to the fluorine at C2 (HCCF angle, red arrow).

For the first analysis, we used the coupling constants of the $^{1}$H-NMR to determine which stereoisomer was produced, more precisely the signal at 3.68 ppm, corresponding to the hydrogen at C3 (dd, $^{3}J_{FH} = 26.41$, $^{3}J_{HH} = 0.9$ Hz, 1H). With an improved Karplus equation, taking electronegativity's of the substituents into account, we calculated the coupling constant of each stereoisomer model (supplementary note table 1, equation 3).[19,20] We received the values for the substituents electronegativity from Altona and used the value of $CH_2F$ for the $CCH_3FC$ group and the value of $CH_2X$ (0.65) for the $CH_2C$ group.[19] Using that analysis we could clearly exclude stereoisomer 2 and 4 as the calculated coupling constants vary widely from the experimental ($^{3}J_{HH} = 0.9$ Hz).

Next, we used the $^{3}J_{FH}$ coupling constant of 26.41 Hz to further distinguish between stereoisomer 1 and 3. We considered a study, which developed a Karplus equation in respect to the CHHF torsion angle (equation 4).[21] We again received the values for the substituents electronegativity from Altona and used the value of $CHCH_3X$ for the $CHCH_3C$ group (0.6) (supplementary note table 2).[19] The results confirmed the $^{3}J_{HH}$ coupling constant analysis as the $^{3}J_{HF}$ coupling constant of stereoisomer 2 and 4 did not correlate with the experimental ($^{3}J_{FH} = 26.41$ Hz). Although the calculated $^{3}J_{FH}$ coupling constant for stereoisomer 1 fits much better to the experimental, we could not exclude stereoisomer 3, especially because we used structural models. Therefore, we performed a NOESY experiments after another purification of the product in higher yields. Unfortunately, we received a mixture of product and educt (see supplementary note figure 3). However, we focused on the hydrogen of the alkene group at C8 ($^{1}$H NMR peak at 6.47 ppm) as the signal of the methyl group at C2 overlapped with the methylene group at C12. In the NOESY experiment, we found NOE signals to 1.18 ppm, 1.21 ppm and 2.00 ppm, which correspond to the methyl group at C10 (distance about 2.3 Å), to the methyl group of C6 (distance about 2.6 Å) and to the methylene group at C5 (distance about 2.2 Å, supplementary note figure 4-5). In contrast we did not found a NOE signal of the methyl group at C2 ($^{1}$H NMR peak at 1.63 ppm) to the alkene group at C8 (distance 2.45 Å in stereoisomer 3) and no signal to the methylene group at C5 (distance 2.03 Å in stereoisomer 3). Therefore, we could exclude stereoisomer 3, as these signals should be observed for that case. The results show that hybrid 1 (**H1**) synthesize stereoisomer 1 with a 2S, 3S configuration.

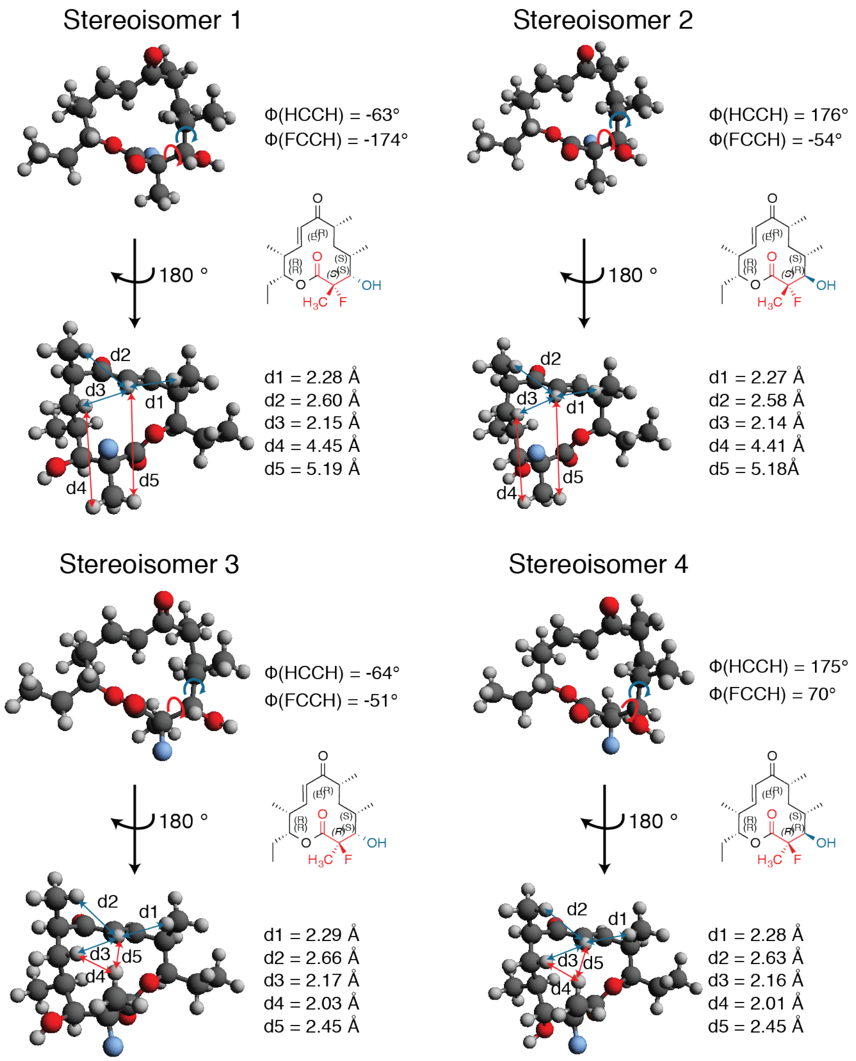

**Supplementary note figure 1: 2D and 3D structure of all possible stereoisomers, created by chemdraw and Avogadro.**

Dihedral torsion angles between HCCH (C3-C4, blue arrow) and HCCF (C2-C3, red arrow) were calculated with Avogadro. Characteristic distances were also calculated by Avogadro and shown with blue arrows (d1-d3) for observed NOE signals and red arrows for non-existent NOE signals.

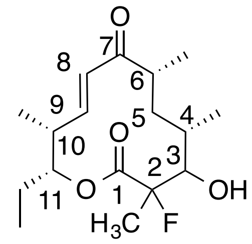

**Supplementary note figure 2: Chemical structure of 18 and numbering of the carbon atoms.**

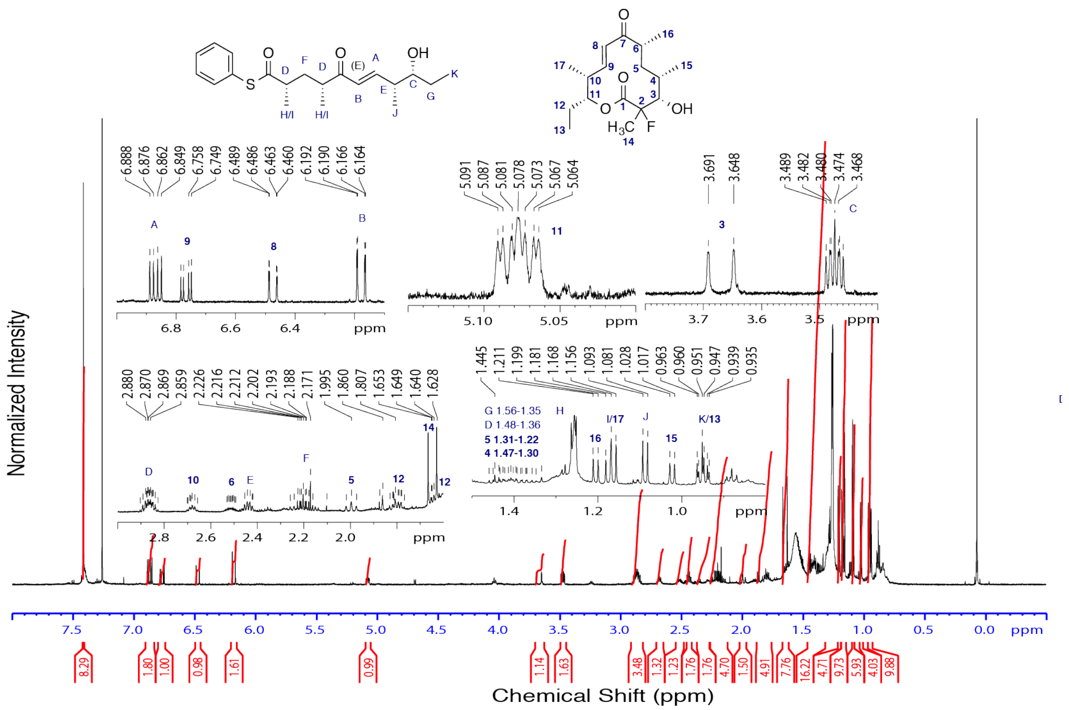

**Supplementary note figure 3: <sup></sup>¹H-NMR analysis of the product/educt mixture and assignment of the peaks.**

The assignment was done with the help of the COSY and HSQC experiment (data not shown). NMR spectrum was processed with TopSpin (version 4.1.1) and structures were created by ChemDraw (version 14.0)

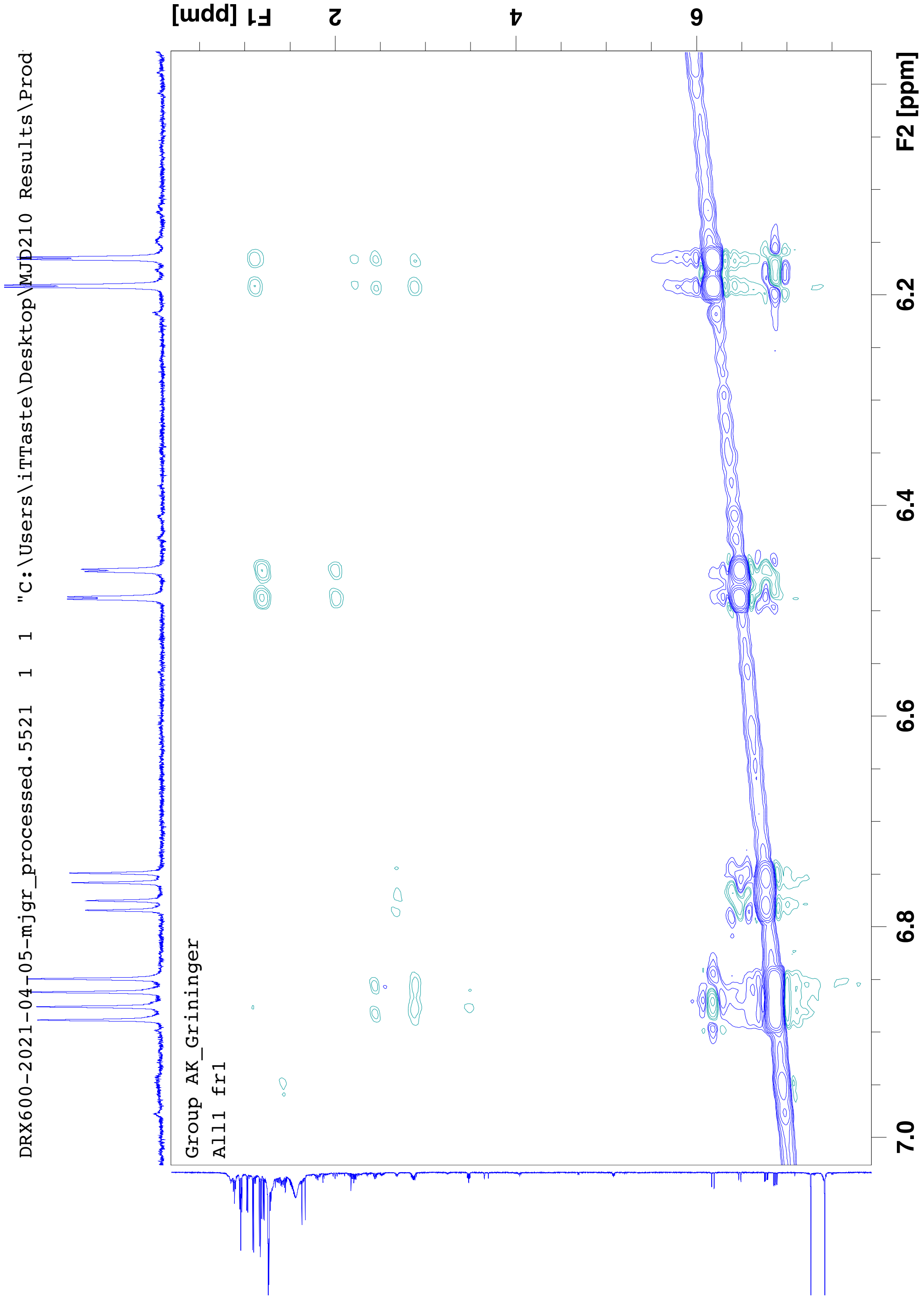

**Supplementary note figure 4: Selected view of the NOESY experiment with compound 18.** NMR spectrum was processed with TopSpin (version 4.1.1).

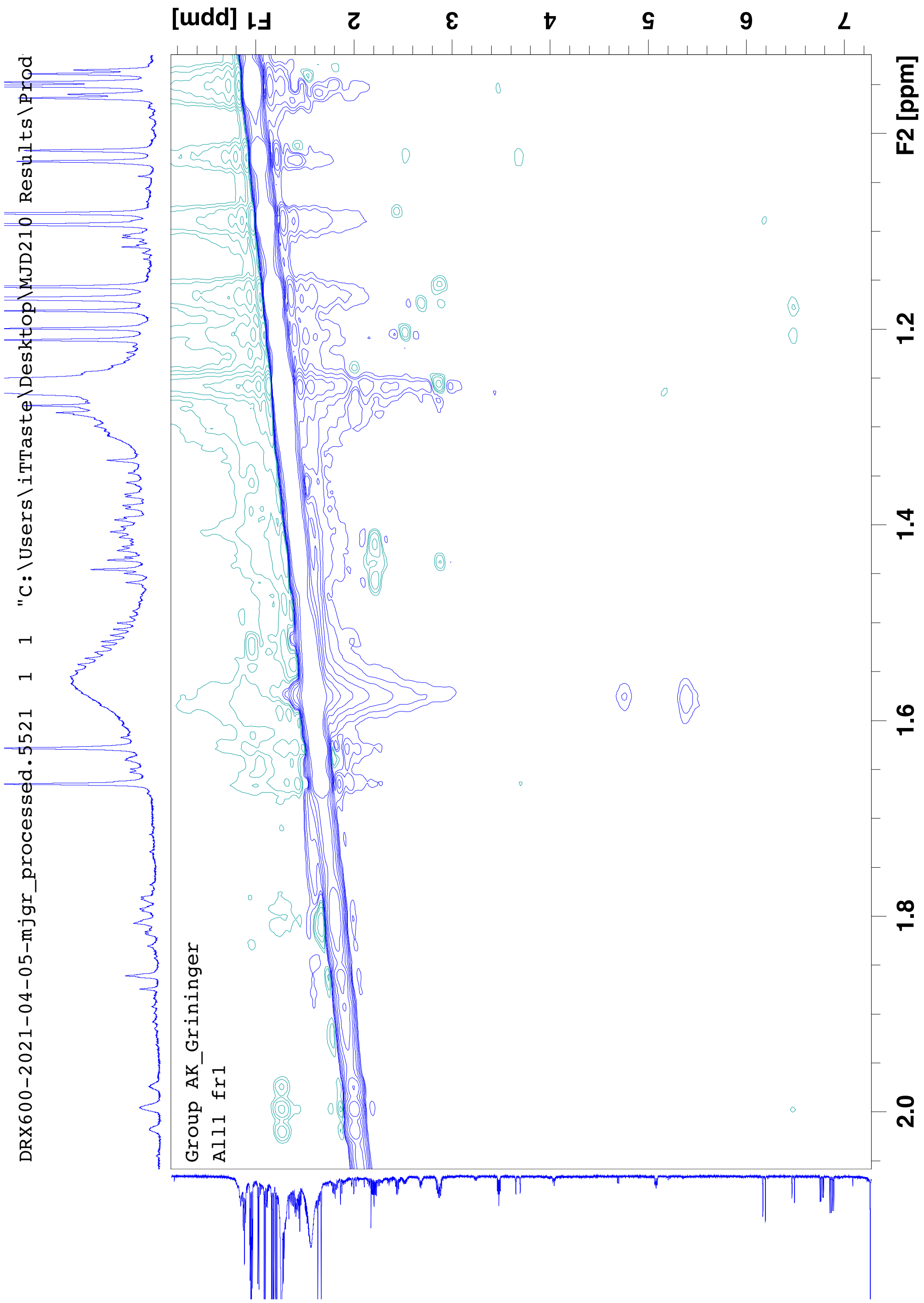

**Supplementary note figure 5: Selected view of the NOESY experiment with compound 18.** NMR spectrum was processed with TopSpin (version 4.1.1).

**Supplementary note table 1: HCCH dihedral angles (Φ) between C3 and C4 and experimental and calculated $^3J_{HH}$ coupling constants.**

$\lambda_i$ is the electronegativity and $s_i$ the sign factor of the substituent.[19,20]

| | $\phi$(HCCH) | $^3J_{HH}$ | $OH: \lambda_1; s_1$ | $CCH_3FC: \lambda_2; s_2$ | $CH_2C: \lambda_3; s_3$ | $CH_3: \lambda_4; s_4$ |
|---|---|---|---|---|---|---|
| Exp. | | 0.9 | | | | |
| Isomer 1 | -63 | 1.28 | 1.33; 1 | 0.65; -1 | 0.65; 1 | 0.8; -1 |
| Isomer 2 | 176 | 12.47 | 1.33; -1 | 0.65; 1 | 0.65; 1 | 0.8; -1 |
| Isomer 3 | -64 | 1.09 | 1.33; 1 | 0.65; -1 | 0.65; 1 | 0.8; -1 |
| Isomer 4 | 175 | 12.40 | 1.33; -1 | 0.65; 1 | 0.65; 1 | 0.8; -1 |

$$3J(H,H) = 14.64\, cos^2(\phi) - 0.78\cos(\phi) + 0.58 + \sum_i \lambda_i\, [0.34 - 2.31\, cos^2(s_i(\phi) + 18.40\, |\lambda i|)] \qquad [3]$$

with $\lambda_i$ = the electronegativity of the substituent, $s_i$ = sign factor of the substituent (see ref 1-2).[19,20]

**Supplementary note table 2: HCCF dihedral angles (Φ) between C2 and C3 and experimental and calculated $^3J_{HF}$ coupling constants.**

$\lambda_i$ is the electronegativity, $\xi_i$ the sign factor of the substituent and $a_{FCC}$ and $a_{HCC}$ the respective bond angles.[19,21]

| | $\phi$(HCCF) | $^3J_{HF}$ | *OH:* $\lambda_1;s_1$ | *CHCH$_3$C:* $\lambda_2;s_2$ | *COOR:* $\lambda_3;s_3$ | *CH$_3$:* $\lambda_4;s_4$ | $a_{FCC};a_{HCC}$ (°) |
|---|---|---|---|---|---|---|---|
| Exp. | | 26.41 | | | | | |
| Isomer 1 | -174 | 26.37 | 1.33; -1 | 0.6; 1 | 0.42; 1 | 0.8; -1 | 111; 104.9 |
| Isomer 2 | -54 | 11.34 | 1.33; 1 | 0.6; -1 | 0.42; 1 | 0.8; -1 | 111; 107.5 |
| Isomer 3 | -51 | 22.41 | 1.33; -1 | 0.6; 1 | 0.42; -1 | 0.8; 1 | 105.9; 102.7 |
| Isomer 4 | 70 | 7.99 | 1.33; 1 | 0.6; -1 | 0.42; -1 | 0.8; 1 | 105.9; 107.8 |

$$3J(H,F) = 40.61\, cos^2(\phi) - 4.22\cos(\phi) + 5.88 + \sum_i \lambda_i\,[-1.27 - 6.20\, cos^2(\xi_i(\phi) + 0.20\,\lambda_i)] - 3.72\left[\frac{(a_{FCC} + a_{HCC})}{2} - 110\right] cos^2(\phi) \qquad [4]$$

with $\lambda_i$ = the electronegativity of the substituent, $\xi_i$ = sign factor of the substituent (see ref 1-2), $a_{FCC}$ and $a_{HCC}$ = bond angles.[19,21]

### 7. Spectra

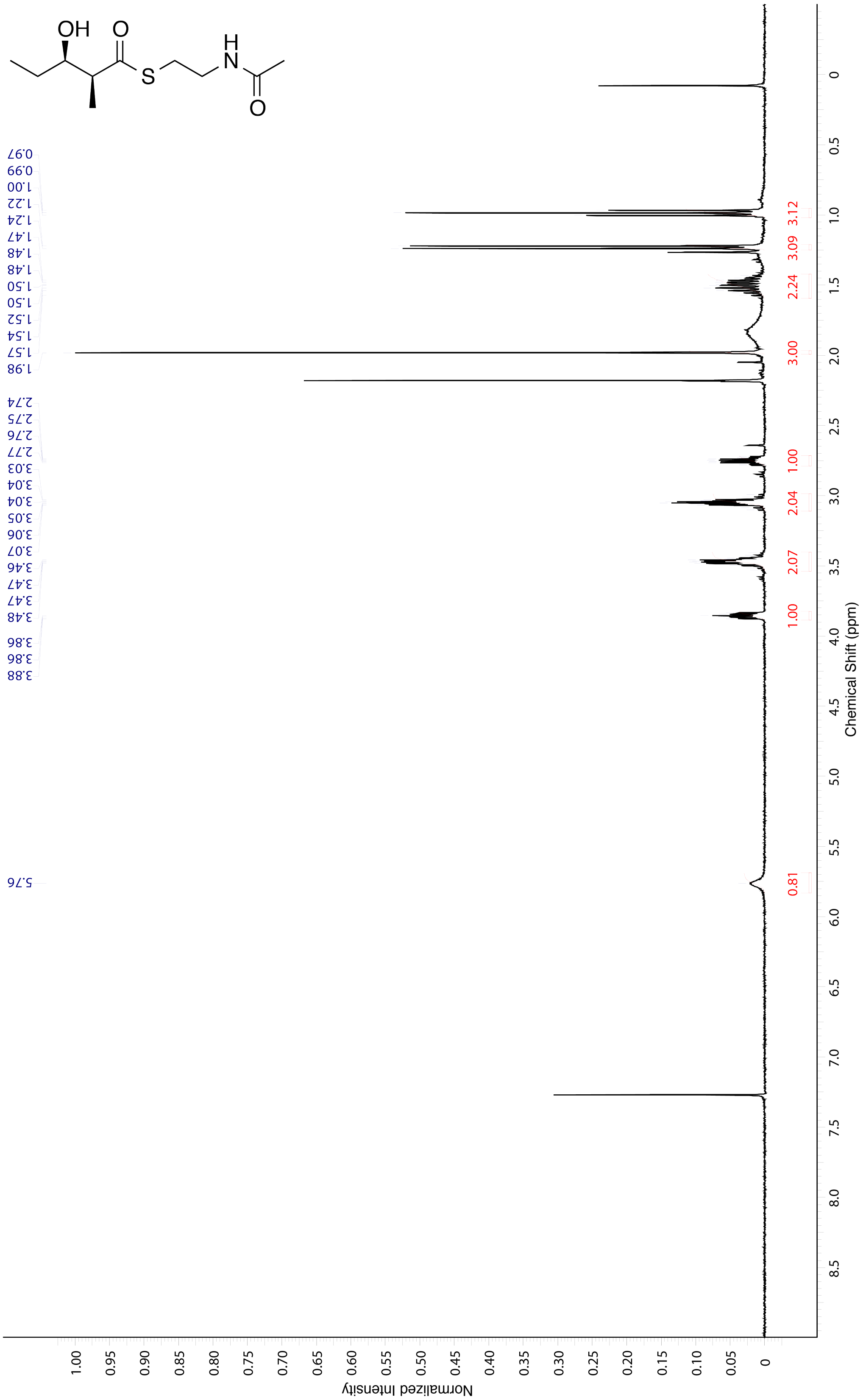

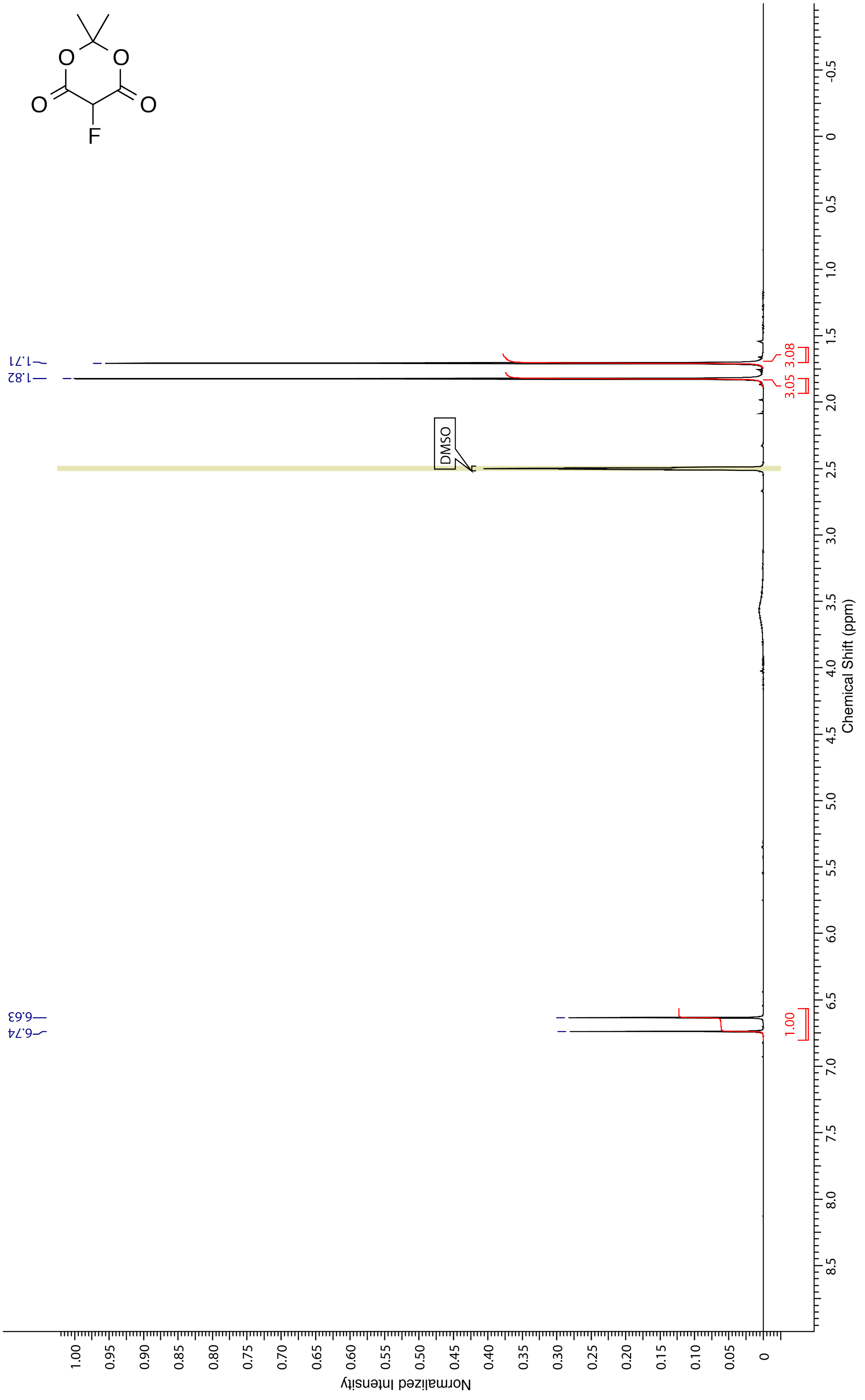

1.71
1.82
6.63
6.74
DMSO
3.08
3.05
1.00
Chemical Shift (ppm)
Normalized Intensity
O
O
O
O
F

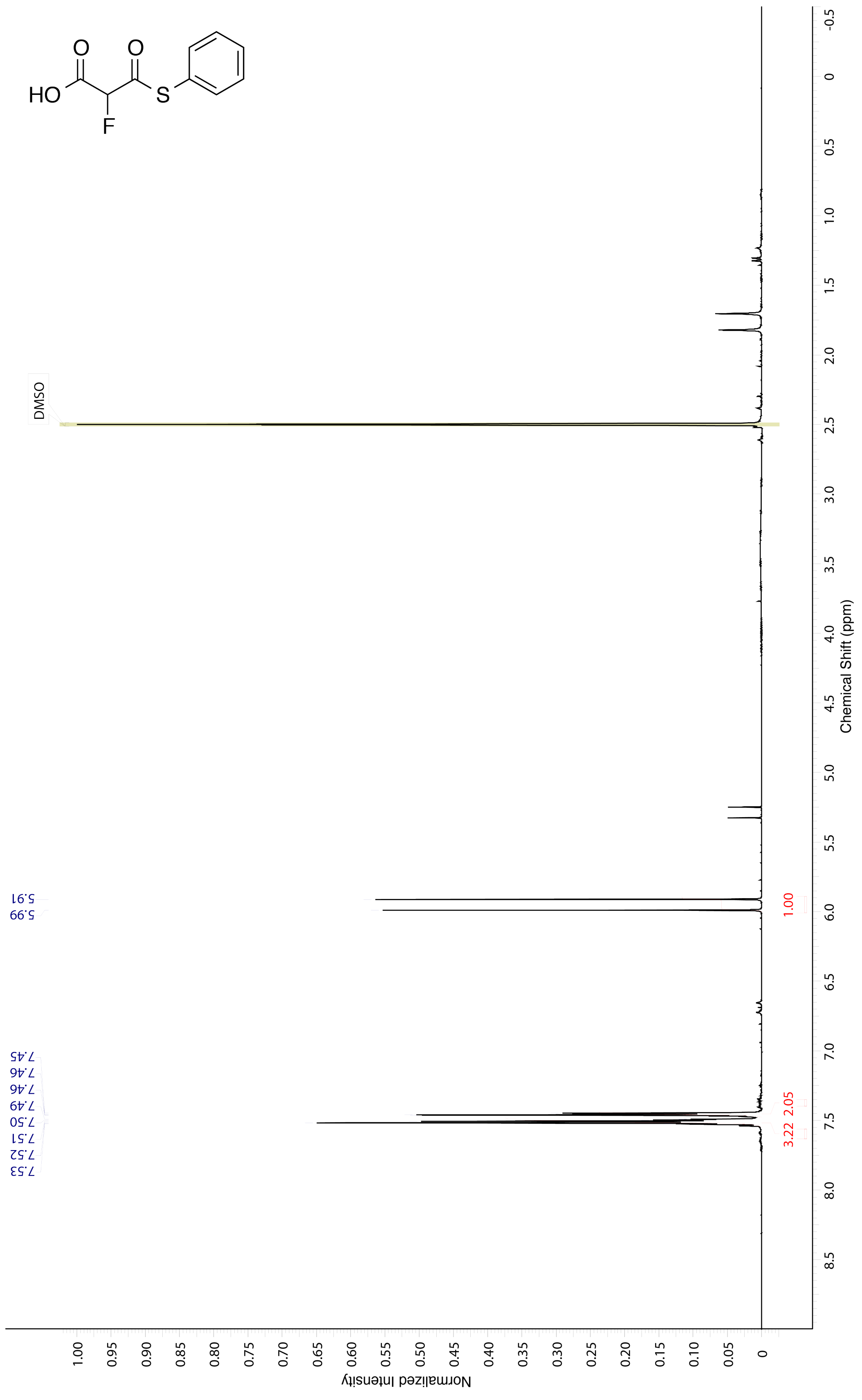

HO
O
F
O
S
DMSO
5.91
5.99
7.45
7.46
7.46
7.49
7.50
7.51
7.52
7.53
1.00
3.22
2.05
Chemical Shift (ppm)
Normalized Intensity

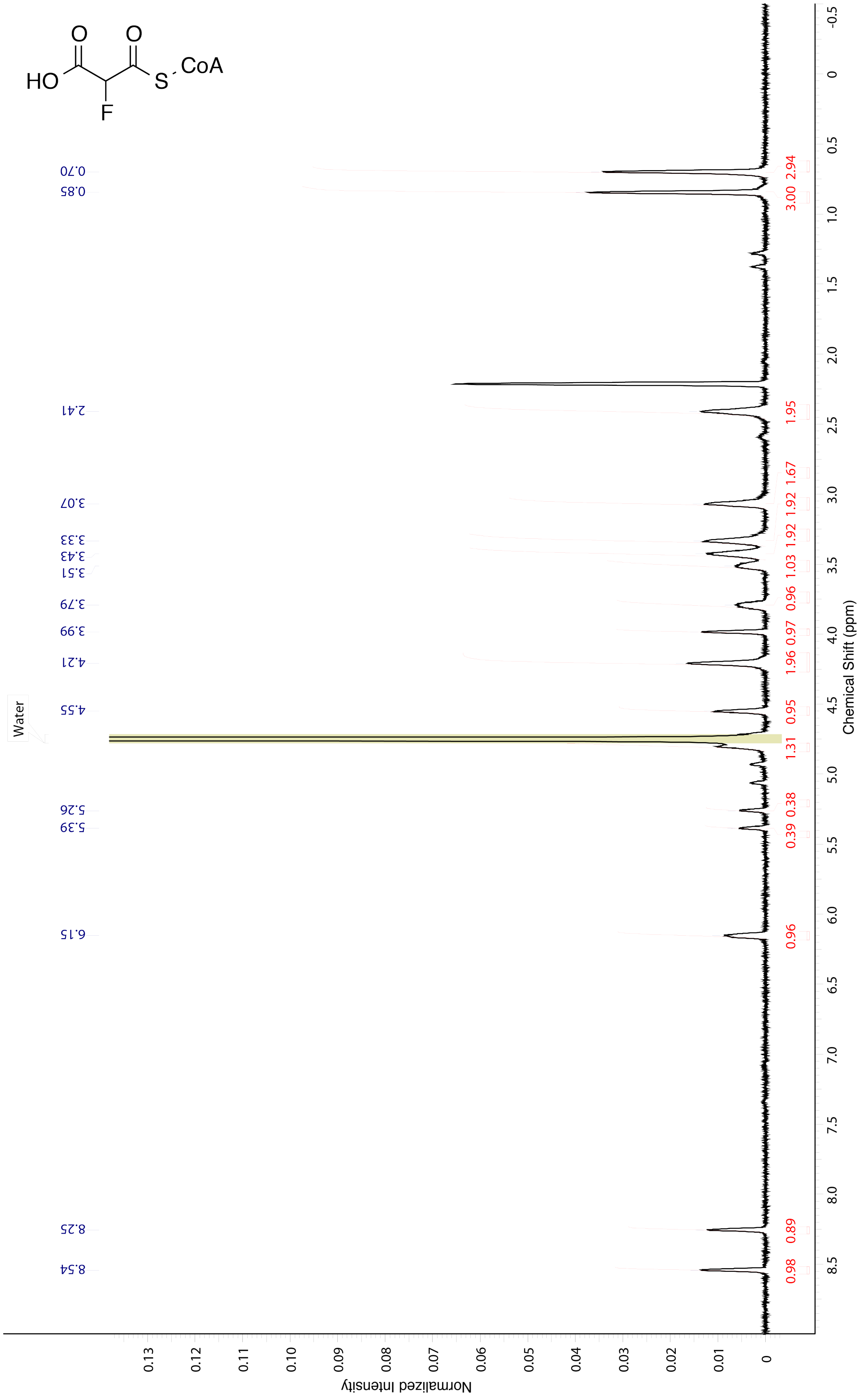

HO
O
O
S
CoA
F
Water
0.70
0.85
2.41
3.07
3.33
3.43
3.51
3.79
3.99
4.21
4.55
5.26
5.39
6.15
8.25
8.54
3.00 2.94
1.95
1.92 1.67
1.92
1.03
0.96
0.97
1.96
0.95
1.31
0.38
0.39
0.96
0.89
0.98
Chemical Shift (ppm)
Normalized Intensity

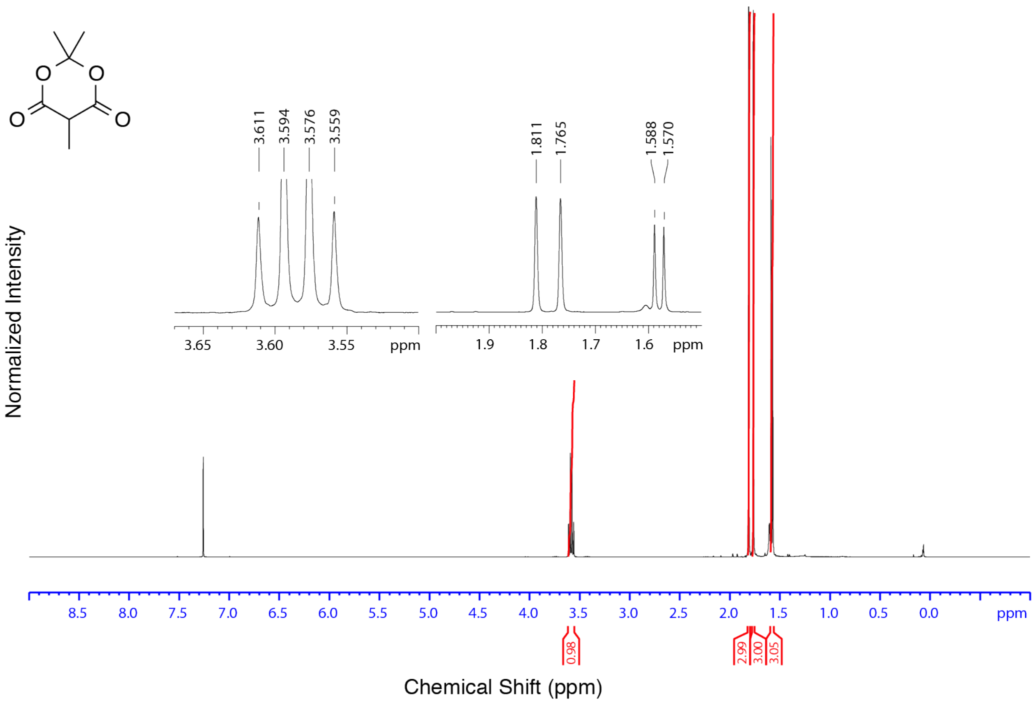

O
O
O
O
3.611
3.594
3.576
3.559
1.811
1.765
1.588
1.570
3.65
3.60
3.55
ppm
1.9
1.8
1.7
1.6
ppm
Normalized Intensity
8.5
8.0
7.5
7.0
6.5
6.0
5.5
5.0
4.5
4.0
3.5
3.0
2.5
2.0
1.5
1.0
0.5
0.0
ppm
0.98
2.99
3.00
3.05
Chemical Shift (ppm)

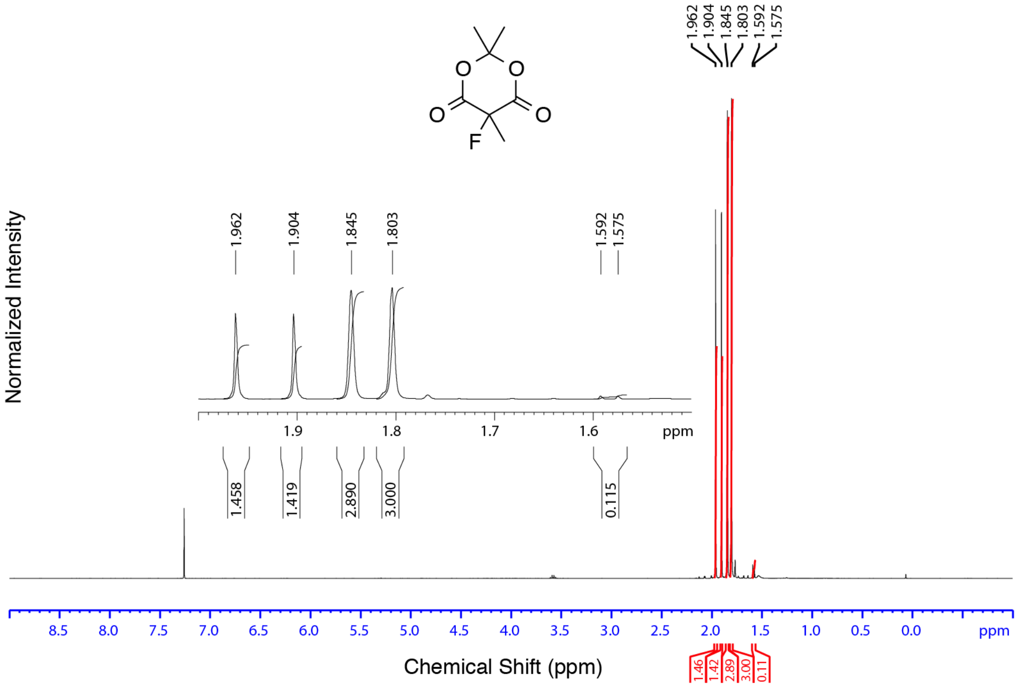

1.962
1.904
1.845
1.803
1.592
1.575
O
O
O
O
F
Normalized Intensity
1.9
1.8
1.7
1.6
ppm
1.458
1.419
2.890
3.000
0.115
8.5
8.0
7.5
7.0
6.5
6.0
5.5
5.0
4.5
4.0
3.5
3.0
2.5
2.0
1.5
1.0
0.5
0.0
ppm
Chemical Shift (ppm)
1.46
1.42
2.89
3.00
0.11

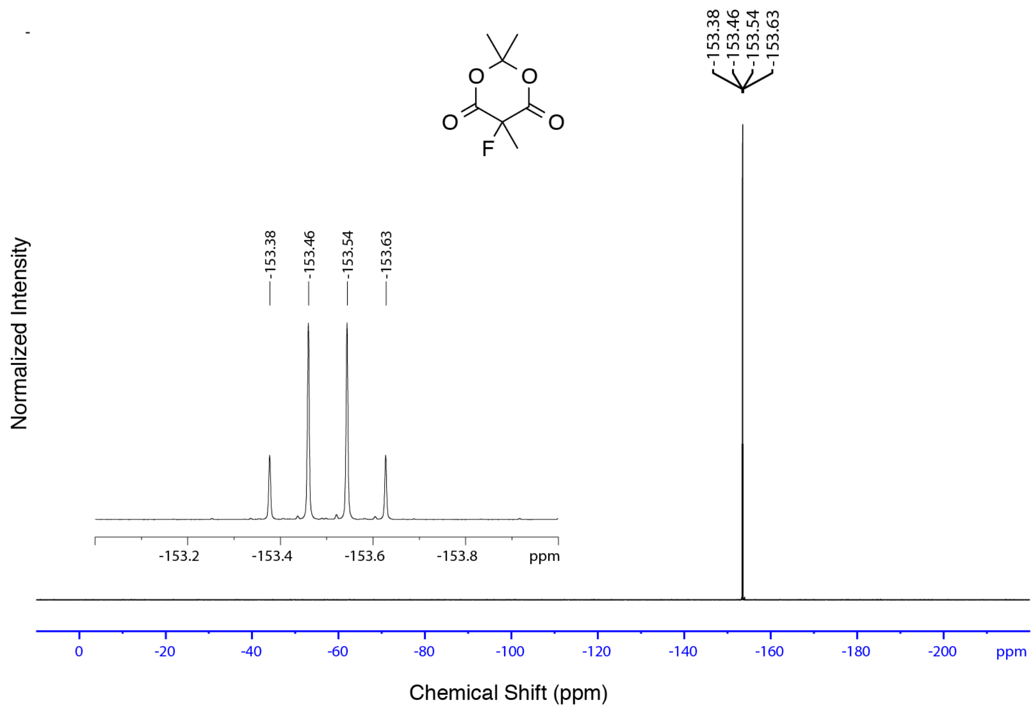
-153.38
-153.46
-153.54
-153.63
Normalized Intensity
-153.2
-153.4
-153.6
-153.8
ppm
0
-20
-40
-60
-80
-100
-120
-140
-160
-180
-200
ppm
Chemical Shift (ppm)

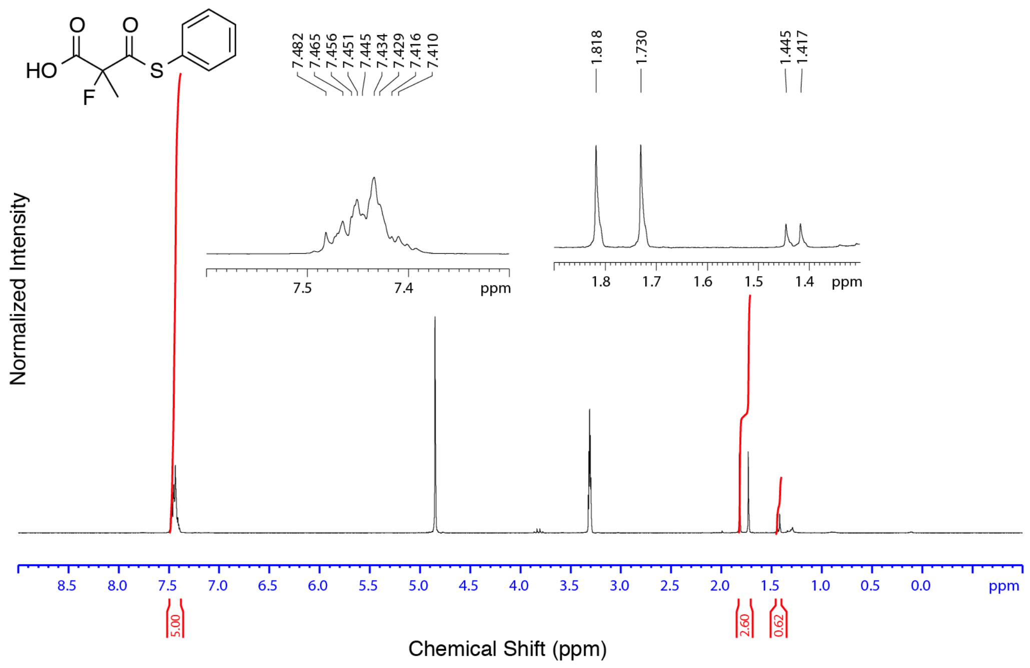
HO
O
O
S
F
7.482
7.465
7.456
7.451
7.445
7.434
7.429
7.416
7.410
1.818
1.730
1.445
1.417
Normalized Intensity
7.5
7.4
ppm
1.8
1.7
1.6
1.5
1.4
ppm
8.5
8.0
7.5
7.0
6.5
6.0
5.5
5.0
4.5
4.0
3.5
3.0
2.5
2.0
1.5
1.0
0.5
0.0
ppm
5.00
2.60
0.62
Chemical Shift (ppm)

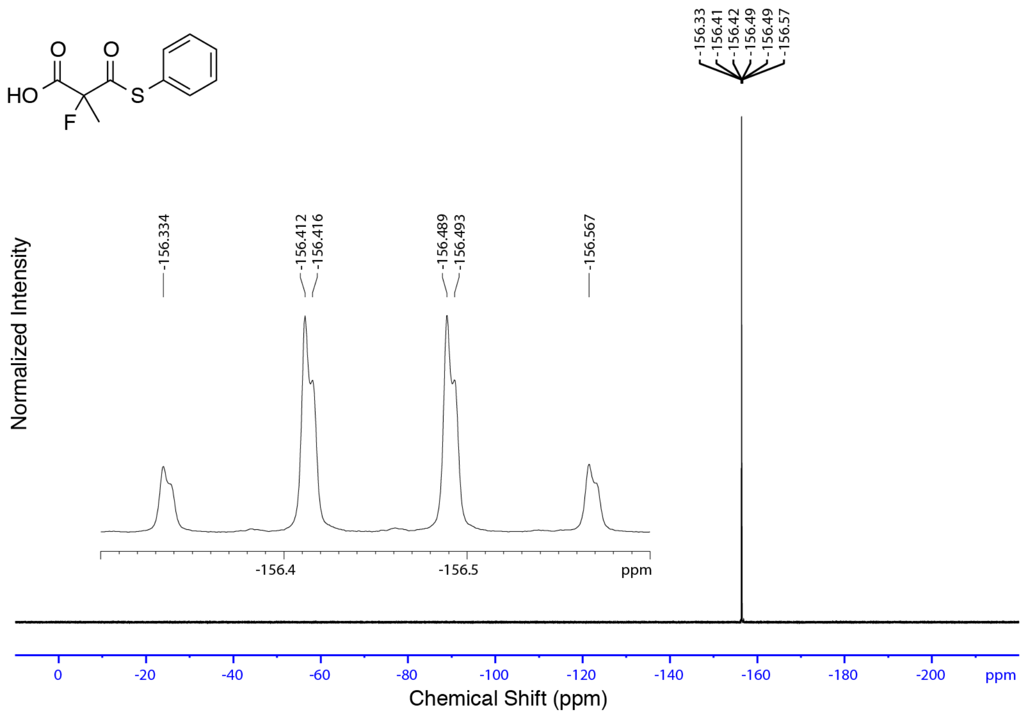
HO
O
O
S
F
-156.33
-156.41
-156.42
-156.49
-156.49
-156.57
Normalized Intensity
-156.334
-156.412
-156.416
-156.489
-156.493
-156.567
-156.4
-156.5
ppm
0
-20
-40
-60
-80
-100
-120
-140
-160
-180
-200
ppm
Chemical Shift (ppm)

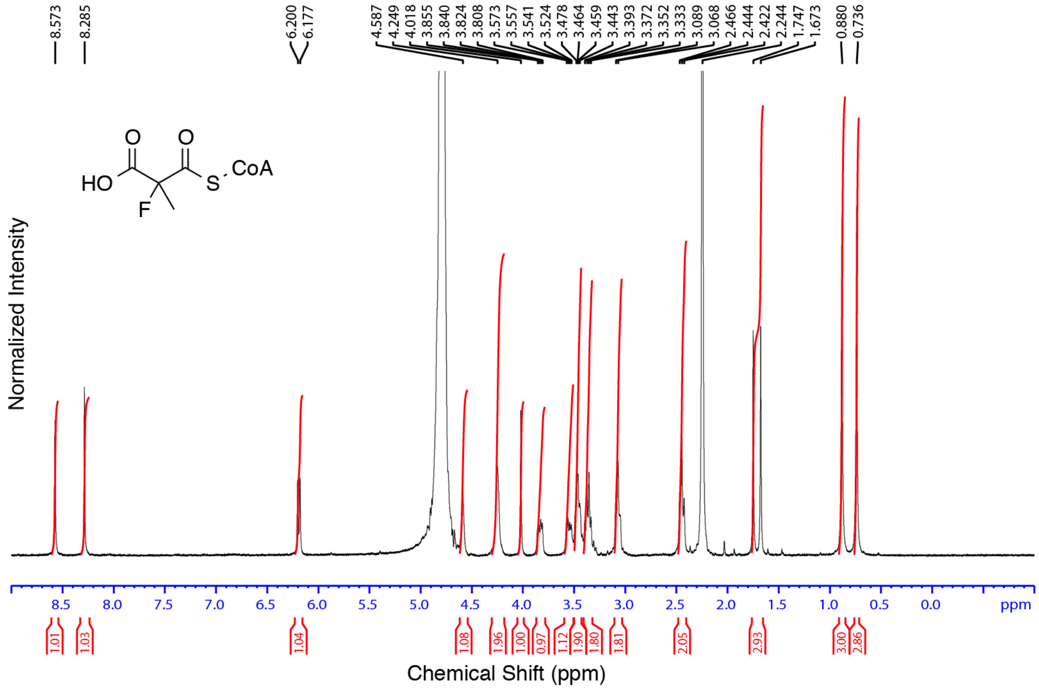
8.573
8.285
6.200
6.177
4.587
4.249
4.018
3.855
3.840
3.824
3.808
3.573
3.557
3.541
3.524
3.478
3.464
3.459
3.443
3.393
3.372
3.352
3.333
3.089
3.068
2.466
2.444
2.422
2.244
1.747
1.673
0.880
0.736
HO
O
O
S
CoA
F
Normalized Intensity
8.5
8.0
7.5
7.0
6.5
6.0
5.5
5.0
4.5
4.0
3.5
3.0
2.5
2.0
1.5
1.0
0.5
0.0
ppm
1.01
1.03
1.04
1.08
1.96
1.00
0.97
1.12
1.90
1.80
1.81
2.05
2.93
3.00
2.86
Chemical Shift (ppm)

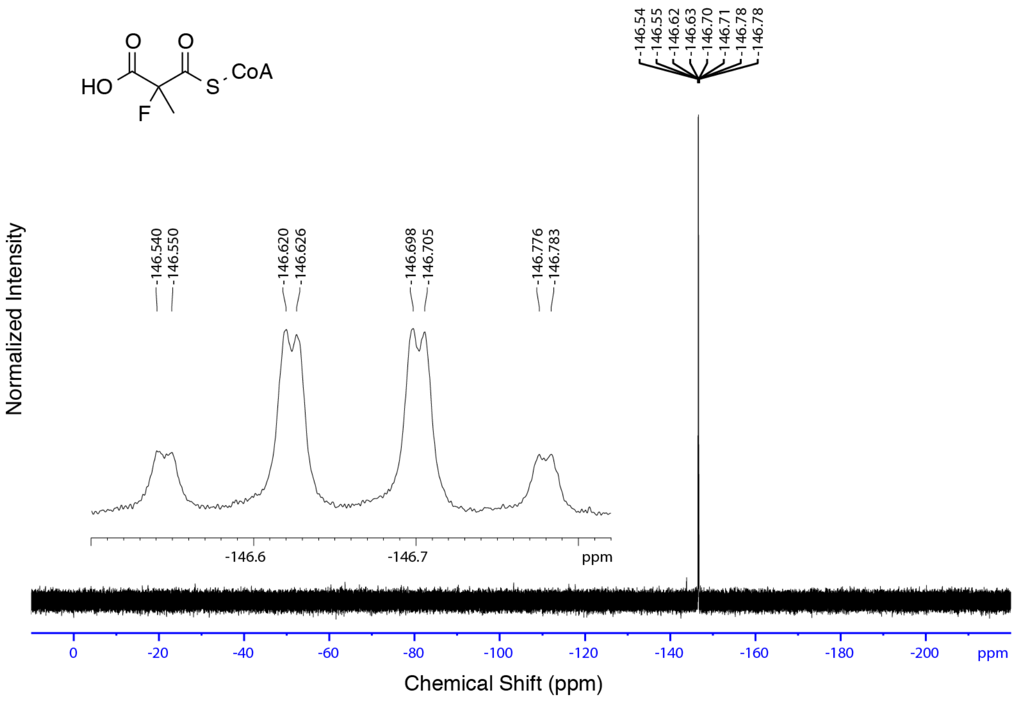

-146.54
-146.55
-146.62
-146.63
-146.70
-146.71
-146.78
-146.78
O
O
HO
S
CoA
F
Normalized Intensity
-146.540
-146.550
-146.620
-146.626
-146.698
-146.705
-146.776
-146.783
-146.6
-146.7
ppm
0
-20
-40
-60
-80
-100
-120
-140
-160
-180
-200
ppm
Chemical Shift (ppm)

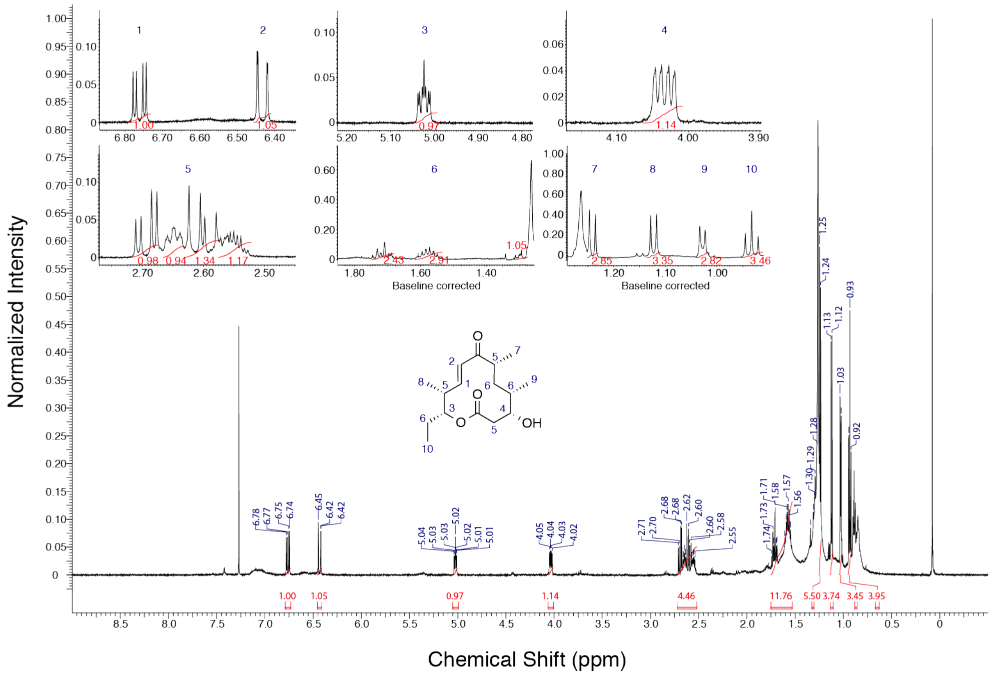
Normalized Intensity
Chemical Shift (ppm)
Baseline corrected
Baseline corrected
OH
1.00
1.05
0.97
1.14
4.46
11.76
5.50
3.74
3.45
3.95

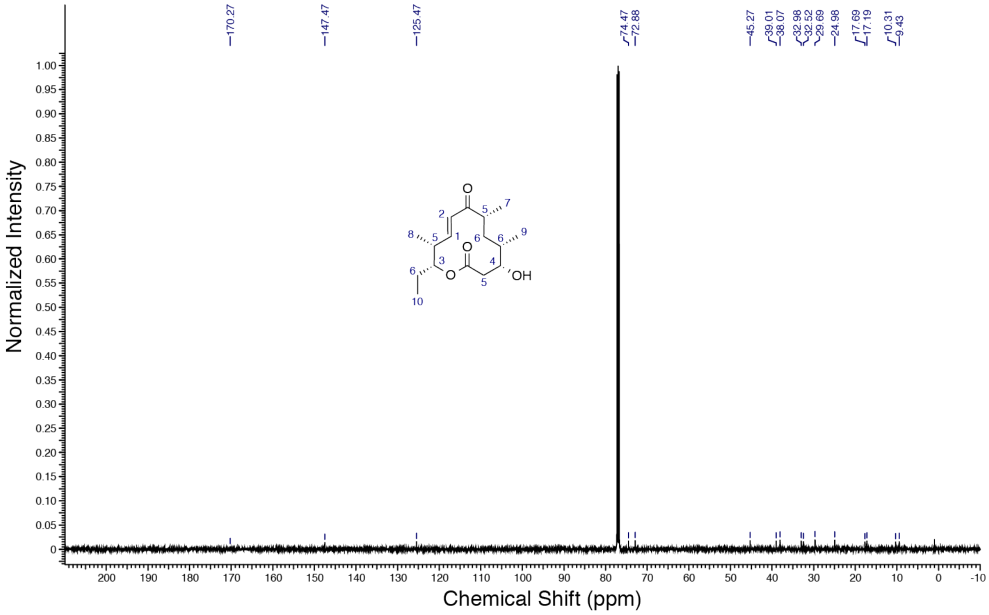
170.27
147.47
125.47
74.47
72.88
45.27
39.01
38.07
32.98
32.52
29.69
24.98
17.69
17.19
10.31
9.43
Normalized Intensity
Chemical Shift (ppm)
OH

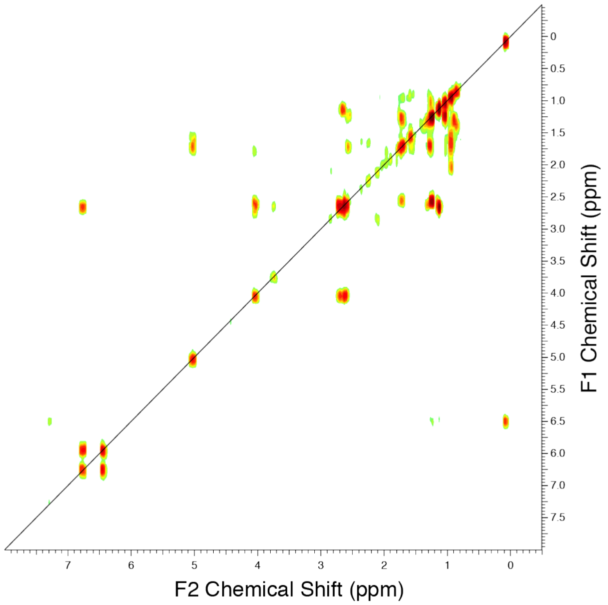
F1 Chemical Shift (ppm)
F2 Chemical Shift (ppm)

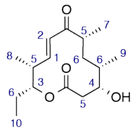

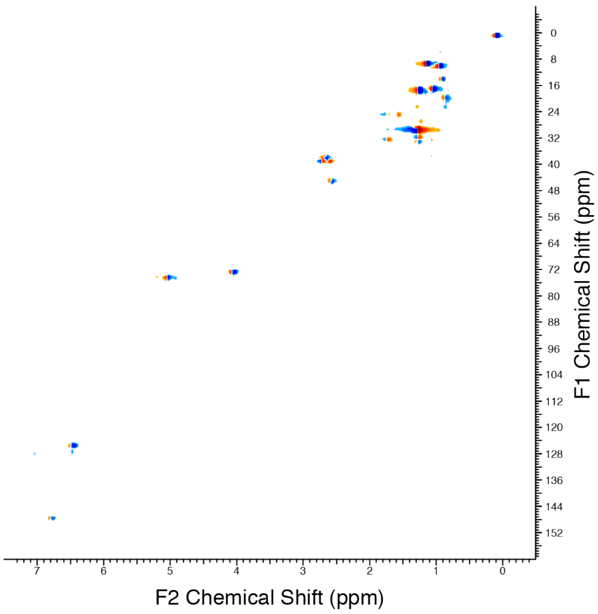
F1 Chemical Shift (ppm)
F2 Chemical Shift (ppm)

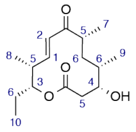

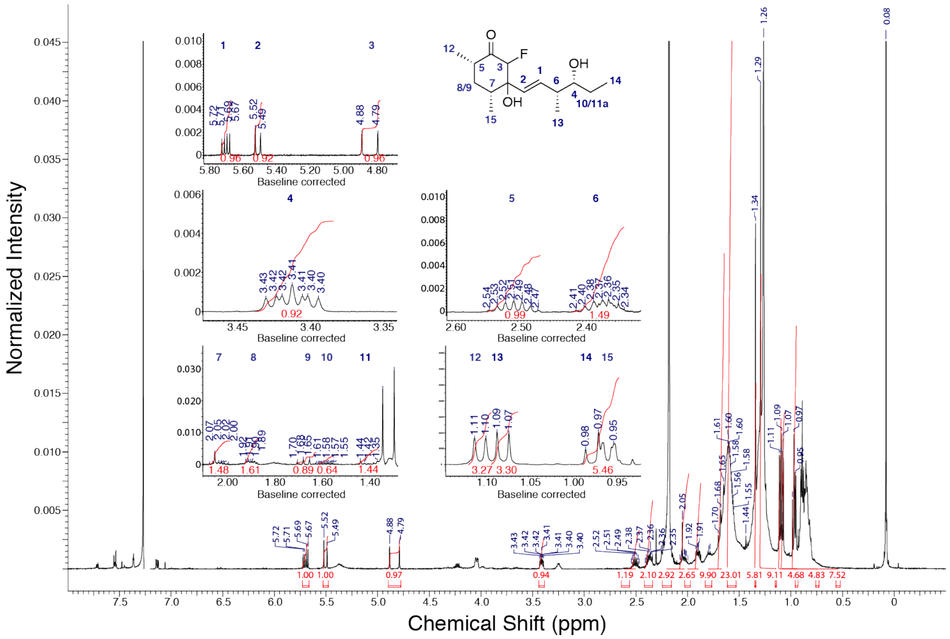

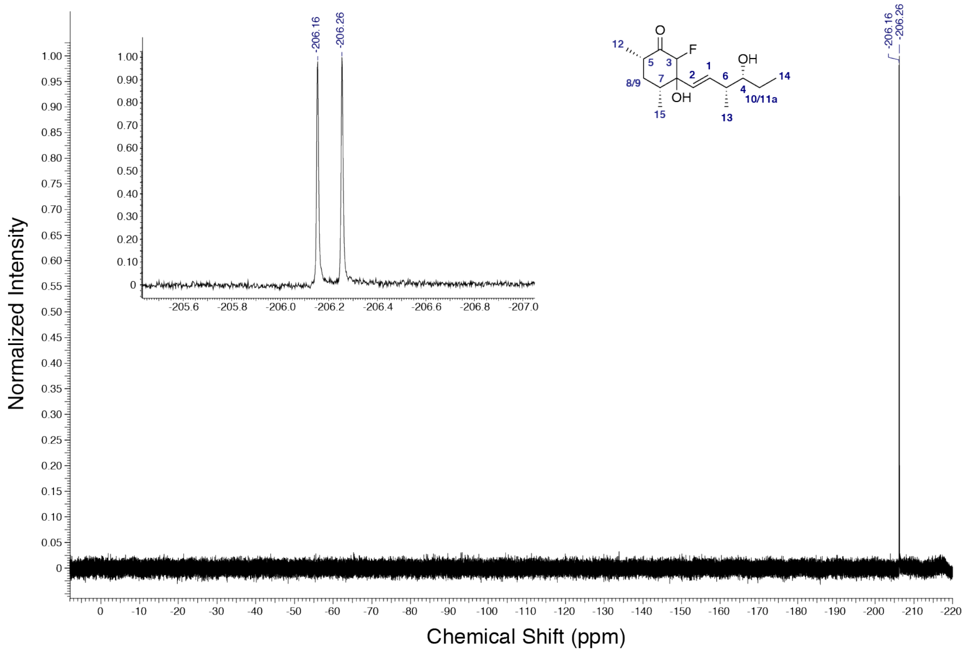

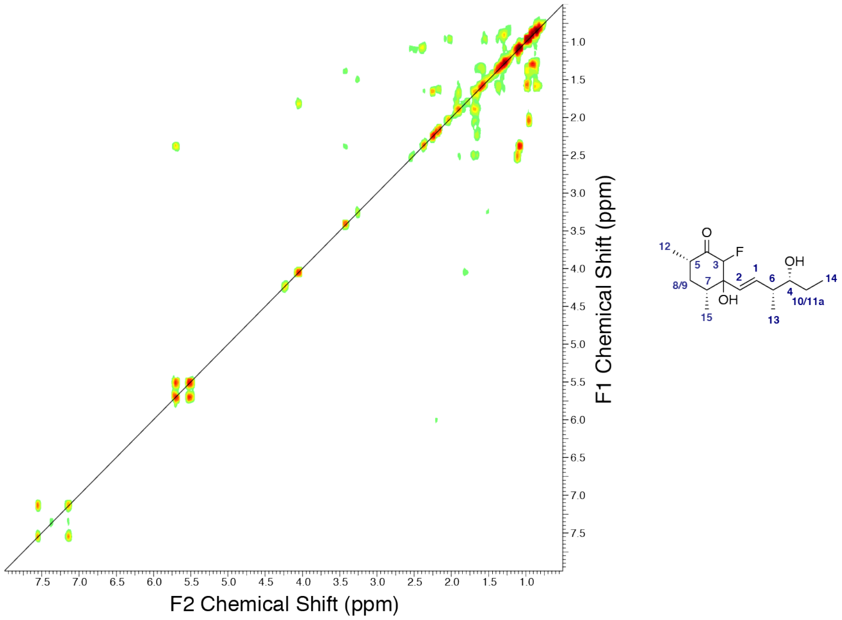
F2 Chemical Shift (ppm)
F1 Chemical Shift (ppm)
O
F
OH
OH
12
5
3
1
2
6
7
4
14
8/9
15
13
10/11a

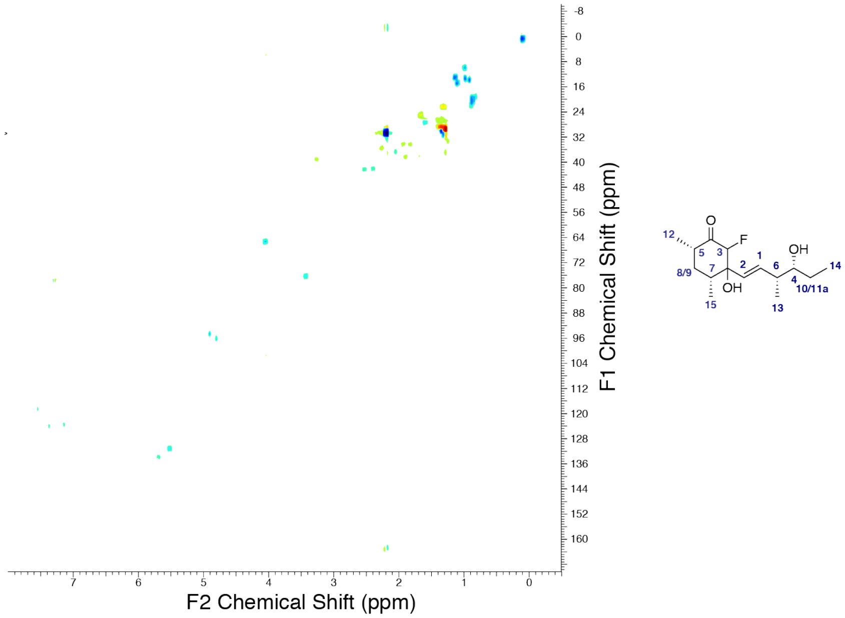
F2 Chemical Shift (ppm)
F1 Chemical Shift (ppm)
O
F
OH
OH
12
5
3
1
2
6
7
4
14
8/9
15
13
10/11a

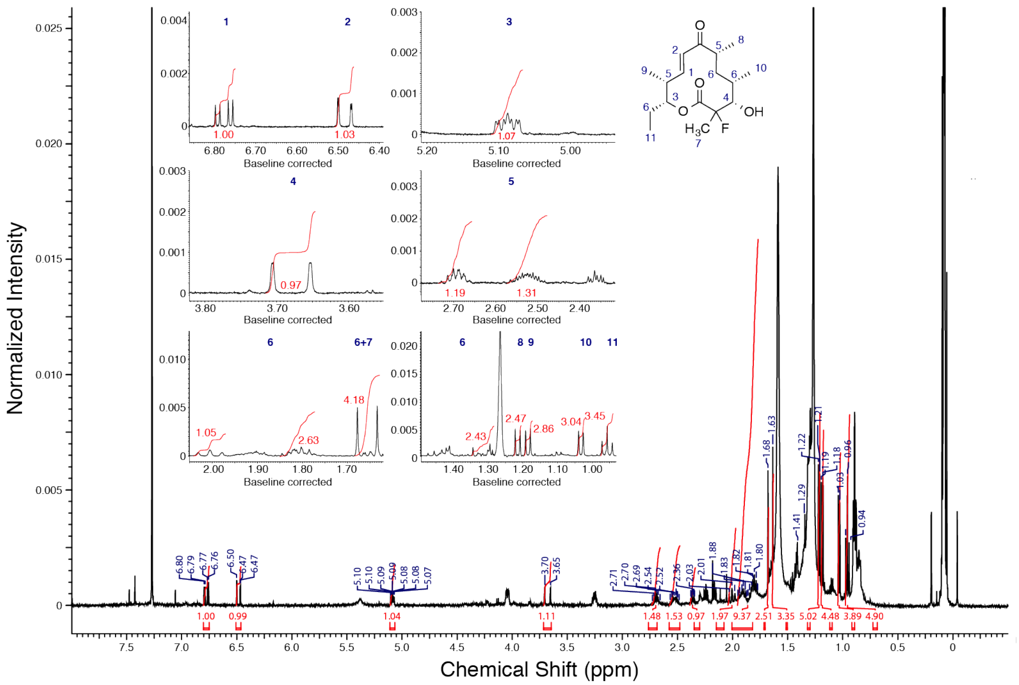
Normalized Intensity
Chemical Shift (ppm)
Baseline corrected
OH
F

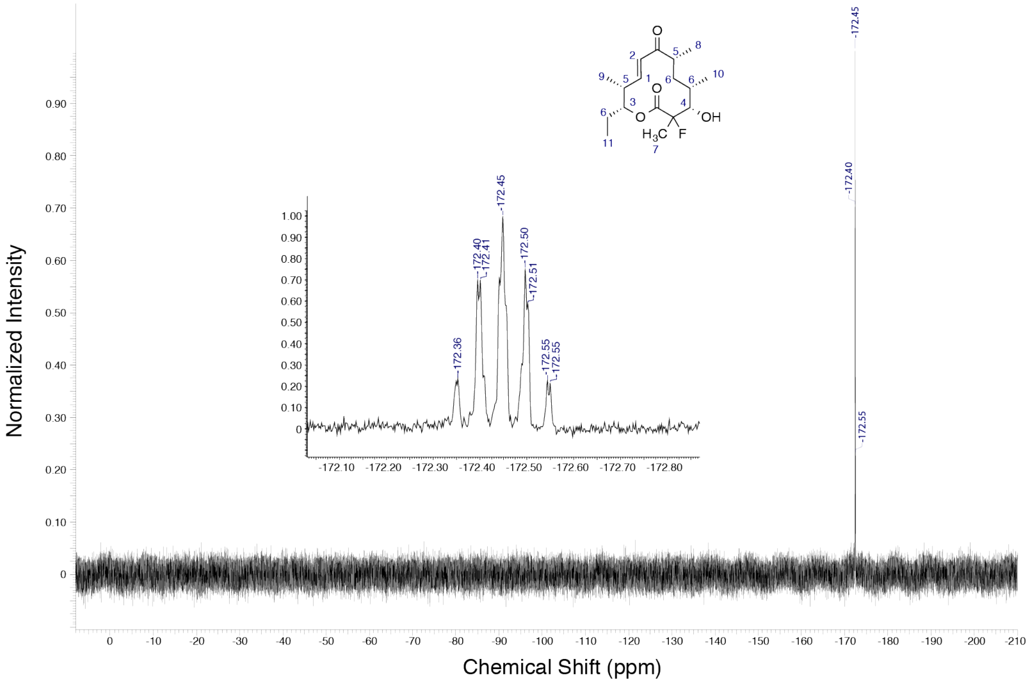
Normalized Intensity
Chemical Shift (ppm)
-172.45
-172.40
-172.55
OH
F

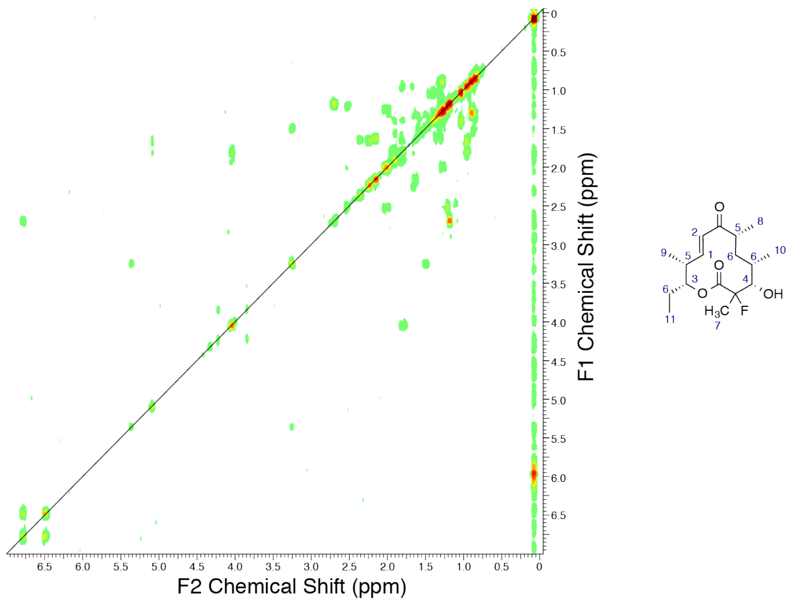

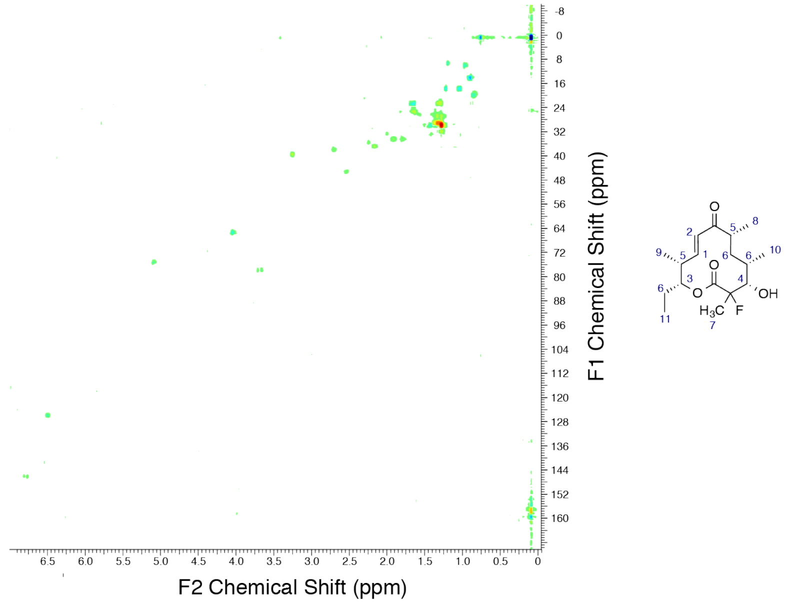
